# Discovery, characterisation and optimisation of bicyclic peptide inhibitors that disarm *Staphylococcus aureus* α-hemolysin

**DOI:** 10.64898/2026.03.09.710508

**Authors:** Jonathan R. Whiteside, Nick Lewis, Laura Díaz-Sáez, Hector Newman, Sarah Newell, Tazmin T. Martin, Jack Butler, Michael J. Skynner, Michael J Dawson, Paul Beswick, Christopher G. Dowson, Catherine E. Rowland

## Abstract

α-Hemolysin (Ahly) is a major *Staphylococcus aureus* virulence determinant implicated in tissue injury and immune dysregulation; antibody inhibitors have reached clinical trials but alternatives with improved ease of manufacture and tissue penetration are desirable. Here we demonstrate that phage-derived bicyclic peptides can serve as compact, chemically tractable Ahly neutralisers. Using TATB-scaffolded M13 phage libraries we identified WNP-motif containing bicyclic binders, with a lead hit of Peptide 14 (K_D_ = 1792nM) and progressed the lead by iterative affinity maturation to Peptide 20 (K_D_ = 609 nM) and by incorporation of strategically chosen non-canonical amino acids to yield Peptide 88 (KD = 96 nM). A 2.2 Å co-crystal structure with AhlyH35A locates the binding footprint on the rim domain and explains the critical role of the WNP motif in target engagement. Functional assays show that the Peptide 88 blocks Ahly mediated hemolysis, inhibits Ahly driven ADAM10 activation, and elucidate its inhibitory mechanism of preventing Ahly binding to human A549 epithelial cells. Peptide 88 protects A549 cells from recombinant toxin and attenuates cytotoxicity in *S. aureus* co-culture experiments, whilst showing no toxicity to A549 cells. Bicyclic peptides thus represent a new and promising anti-virulence modality: small, synthetically accessible molecules that mimic antibody recognition, with therapeutic potential against *S. aureus* infections.

## Introduction

Antimicrobial resistance (AMR) represents one of the most pressing threats to global public health, with the World Health Organisation (WHO) estimating 1.27 million deaths attributable to AMR in 2019, a figure projected to rise to 1.91 million deaths by 2050^1,2^. Among the pathogens contributing to this crisis, the Gram-positive bacterium *Staphylococcus aureus* is of particular concern, ranking as the second leading cause of mortality due to bacterial infection globally with approximately 1.1 million deaths in 2019^3^. Furthermore, reports consistently place *S. aureus* in the top 5 leading causes of death attributed to AMR, with this only expected to rise by 2050^1,2^.

*S. aureus* is a versatile pathogen capable of causing a range of infections from more common skin and soft tissue infections to severe invasive diseases such as bacteraemia, pneumonia, endocarditis and sepsis^4–6^. Although methicillin was introduced in 1959 to combat *S. aureus* infections, resistance emerged within a year, marking the beginning of the rapid spread of methicillin-resistant *Staphylococcus aureus* (MRSA) in both clinical and community settings^7^. Between 2001 and 2009 MRSA accounted for between 13% and 74% of global *S. aureus* cases depending on geographical region^8–10^. Resistance persists despite the development of new antimicrobial classes, largely due to the evolutionary exposure of *S. aureus* to natural product derived antibiotics and its capacity for resistance via genetic mutation and horizontal gene transfer^11^. These factors contribute to the ongoing challenge of managing MRSA infections, thus creating an urgent need for novel *S. aureus* counter measures.

An alternative approach lies in targeting bacterial virulence rather than viability. Bacteria that colonise the human body typically adopt one of two distinct niches: a commensal or mutualistic role, exemplified by members of the gut microbiota; or a pathogenic role. The latter is characterised by the expression of virulence factors that facilitate in immune system evasion, host tissue invasion and nutrient acquisition enabling the pathogen to establish a virulent infection^12–14^. Anti-virulence strategies aim to neutralise these factors, effectively disarming pathogens and enhancing clearance by the hosts immune system^15–17^. Unlike traditional broad-spectrum antibiotics, anti-virulence therapies selectively target pathogenic mechanisms, preserving beneficial microbiota and potentially reducing the selective pressure for resistance development^18^.

Pore forming toxins (PFTs) are a major class of virulence factors that compromise host cell membranes, contributing to immune system evasion, tissue invasion and intracellular survival^19^. These toxins are secreted as soluble monomers that bind to specific receptors on host cells, oligomerise and insert into the membrane to form large transmembrane pores^19–25^. The functional consequences of pore formation include osmotic dysregulation, ion flux and ultimately cell lysis as well as the activation of host cell death pathways such as necrosis, apoptosis and pyroptosis^26,27^. Genetic and pharmacological studies, including clinical trials, have highlighted PFTs as playing an important role in bacterial pathogenesis^28–30^. Among the best characterised PFTs is α-hemolysin (Ahly) which plays a central role in staphylococcal virulence^31^. Ahly is secreted as a monomer that binds to the host cell surface via the metalloprotease ADAM10 and assembles into a heptameric transmembrane β-barrel pore^22,31–33^. The mature pore permits the uncontrolled passage of water, ions, and small molecules (1-4 kDa), which can result in osmotic lysis of target cells^34^. Beyond direct cytolysis, Ahly modulates host-pathogen interactions through several non-lytic mechanisms. Notably, Ahly upregulates the protease activity of ADAM10, which is thought to aid in disruption of endothelial barriers and thus facilitate *S. aureus* tissue invasion^22,31,35,36^. Ahly also promotes intracellular survival and immune evasion of *S. aureus*^37,38^. Clinical data demonstrate that high anti-Ahly antibody titres correlate with protection against invasive *S. aureus* infections^39,40^. Its multifaceted role as a prominent *S. aureus* virulence factor has led to interest from pharmaceutical companies with various anti-Ahly antibodies entering clinical trials. Two agents, AR-301 (Tosatoxumab) and AR-320 (Suvratoxumab) have reached phase 3 clinical trials^41–43^.

Cyclic peptides are found abundantly in nature as antimicrobials produced by various bacterial species as part of the ‘evolutionary arms race’ of intraspecies competition. Several cyclic peptides have been developed into clinically approved antibiotics including daptomycin, telavancin, dalbavancin and oritavancin, all of which were derived from natural products^44^. The constrained conformation of cyclic peptides reduces the entropic penalty associated with target binding, enhancing affinity while minimising off-target binding by limiting alternative conformations^45,46^. Furthermore, this structural rigidity also confers improved resistance to proteolytic degradation^47–49^. Compared to antibodies, cyclic peptides offer advantages such as better tissue penetration, reduced immunogenicity and improved potential for cellular uptake and oral administration, whilst being more amenable to chemical modification and having a reduced manufacturing cost^50–58^.

Bicycle® molecules are bicyclic peptides which represent a promising subclass of constrained peptides. Bicycle molecules are formed by cyclisation of a linear peptide via covalent attachment of cysteine thiols to a central trivalent chemical scaffold. Using phage display technology, highly diverse genetically encoded libraries of bicyclic peptides can be generated and screened against target proteins *in vitro*^59^. Iterative rounds of selection enrich for high affinity binders, which are then identified by sequencing and chemically synthesised for further characterisation (Fig. 1). Unlike natural product cyclic peptide discovery, Bicycle Therapeutics’ phage display platform enables *de novo* generation of bicyclic peptides against virtually any protein target, without relying on the existence of a naturally evolved cyclic peptide ligands to said target. This synthetic approach greatly expands the accessible chemical space beyond those found in nature and allows novel ligands to be identified to specific targets. Targeting bacterial proteins with synthetic bicyclic peptides offers a key advantage in antimicrobial development: the absence of prior evolutionary exposure reduces the likelihood of innate or acquired resistance mechanisms, thereby potentially enhancing therapeutic efficacy. The Bicycle Therapeutics’ phage display platform has previously generated bicyclic peptides that have demonstrated therapeutic potential against key anti-infective targets such as the SARS-CoV-2 spike protein and *Escherichia coli* penicillin binding proteins^60,61^.In this study, we sought to evaluate whether bicyclic peptides could serve as a novel class of anti-virulence agents by targeting Ahly. We applied Bicycle Therapeutics’ phage display platform to discover and optimise bicyclic peptide inhibitors of Ahly, with the aim of developing potent neutralising molecules to this critical virulence factor.

**Figure 1.**
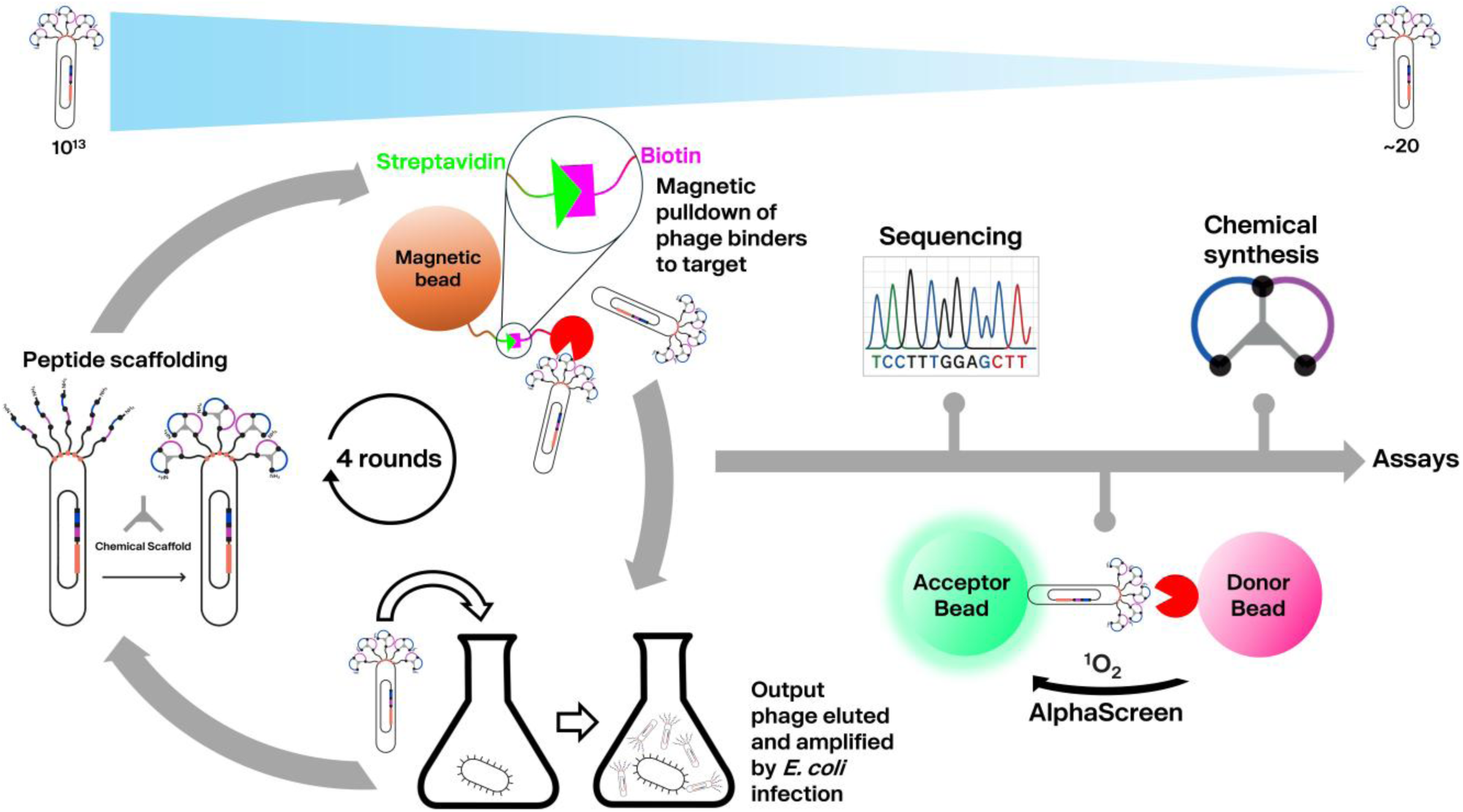
Schematic overview of bicyclic peptide phage display. M13 phage are genetically engineered to display peptide sequences on the pIII coat protein. Each displayed peptide contains three fixed cysteine residues flanking randomized sequences. These peptides are chemically cyclised via thioether bond formation with a trivalent chemical scaffold, yielding bicyclic peptides on the phage surface. Libraries (∼10^13^ diversity) are incubated with a biotinylated target protein, and then captured using magnetic streptavidin beads to isolate binding phage. Bound phage are eluted and amplified in *E. coli* TG1 cells to be used as the input to the next round of selection. After four rounds of selection, individual phage clones are sequenced to identify enriched peptide sequences and screened for target binding using an AlphaScreen assay. Approximately 20 sequences are then selected for solid-phase synthesis and chemical scaffolding to generate purified bicyclic peptides for downstream assays.

## Results

### Bicyclic peptides identified from phage display selections against AviTag-Ahly are capable of inhibiting Ahly toxicity in a hemolysis assay

A C-terminal AviTag was incorporated into the Ahly construct to enable biotinylation for use in Bicycle Therapeutics’ phage display platform. Streptavidin gel-shift analysis demonstrated complete AviTag biotinylation and that the AviTag-Ahly retained a structure and activity comparable to wild-type Ahly (Fig. S1). To discover novel Ahly bicyclic peptide binders, phase display selections were performed using AviTag-biotinylated Ahly (AviTag-Ahly) as the target protein. Following four rounds of phage display selection against AviTag-Ahly using 1,3,5-tris(bromoacetyl) hexahydro-1,3,5-triazine (TATB) scaffolded phage libraries, 245 phage were sequenced to identify their displayed peptide sequences. Binding of each phage clone to AviTag-Ahly was evaluated via AlphaScreen (Fig. 2c). Notably, 69% (170/245) of clones contained a conserved Trp-Asn-Pro (WNP) motif in their expressed peptide sequence. Based on copy number in sequencing output, AlphaScreen results, and attempting to maximise sequence diversity, 19 representative peptides were selected for chemical synthesis and further characterisation (Table 1). Binding affinities of the synthesised bicyclic peptides were measured by surface plasmon resonance (SPR). All but two of the bicyclic peptides that showed detectable binding to AviTag-Ahly contained the ‘WNP’ motif, with one of these two contained a similar Trp-Asn-Thr (WNT) motif (Peptide 7) (Table 1). Peptide 14 exhibited the highest affinity, with a K_D_ of 1.3µM; two more repeats were later performed giving an average K_D_ of 1.8µM over the 3 biological repeats (Table 1, Fig. 3). Due to the prevalence of the ‘WNP’ motif in the bicyclic peptides that exhibited detectable binding to Ahly in SPR we wished to investigate if they all bound to the same epitope on Ahly. To this end, a competition AlphaScreen was performed in a matrix format to test all bicyclic peptides with all respective phage clones (Fig 2c). The phage binding signal of each clone was reduced when ‘WNP’ containing bicyclic peptide was added, except for the phage clone expressing Peptide 14 on its surface (Fig 2b,c). These results support the conclusion that the ‘WNP’ containing bicyclic peptides target the same or an overlapping epitope and, that Peptide 14 exhibits superior affinity, which is in alignment with the SPR data. To investigate if cyclisation is a prerequisite for Ahly binding, binding of the phage clone expressing Peptide 14 to AviTag-Ahly was measured in AlphaScreen using both cyclised (TATB treated) and linear forms (Iodoacetamide treated). Only cyclised phage generated signal in the AlphaScreen assay (Fig. 2d), indicating that peptide cyclisation is critical for the binding interaction of Peptide 14 with AviTag-Ahly. To evaluate whether the highest affinity hit identified from selections was a functional inhibitor, we tested Peptide 14 in a hemolysis assay. We observed specific, dose-dependent inhibition of Ahly mediated erythrocyte lysis with an IC_50_ of 504µM (Fig. 2e, 4a). This inhibitory effect was specific to Ahly as no inhibition was seen against another PFT (pneumolysin) in the same assay (Fig. S2). Having established Peptide 14 as an inhibitor of Ahly pore forming function, we next sought to improve binding affinity, with the goal of enhancing inhibitory potency.

**Figure 2.**
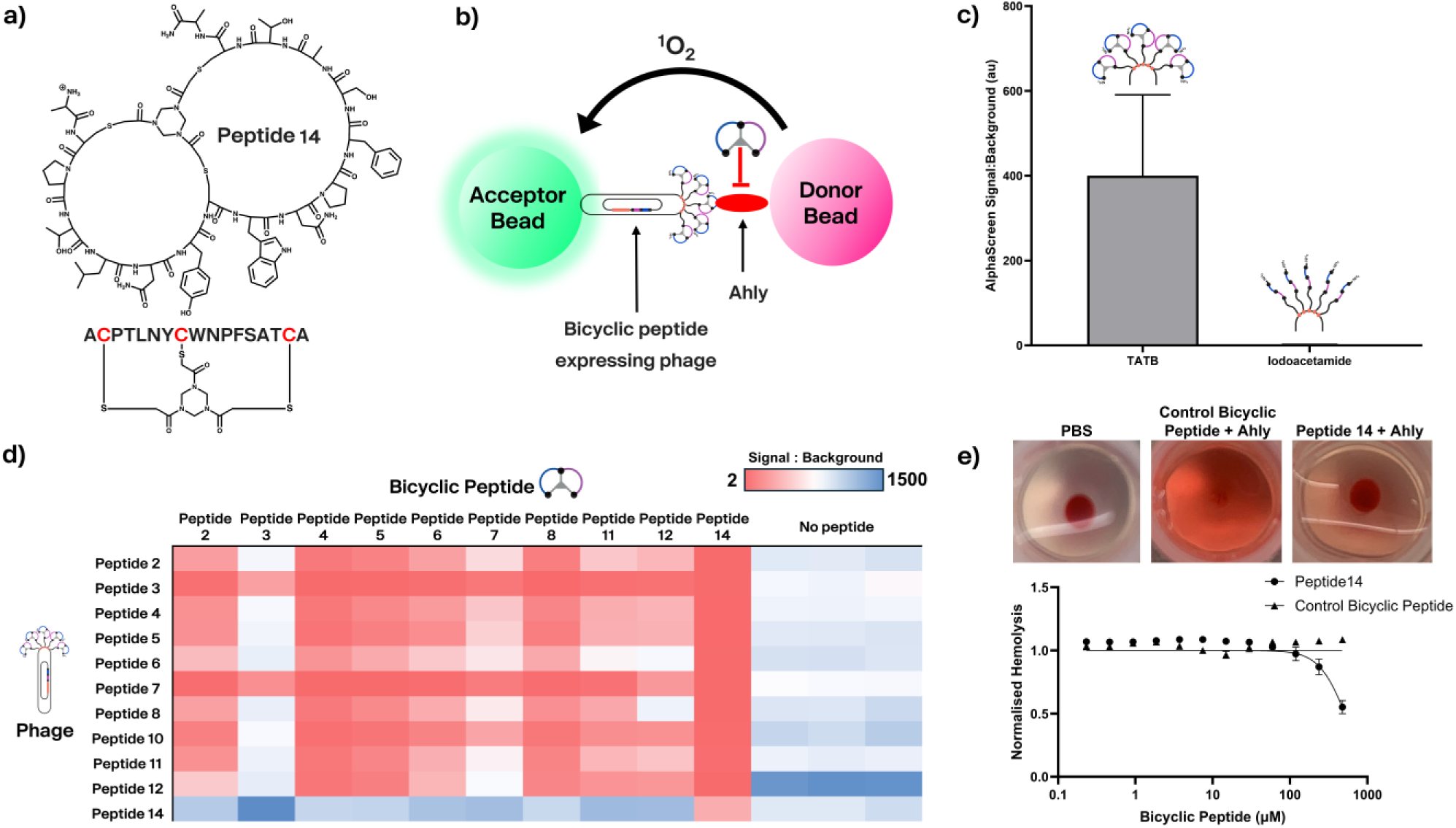
Identification of bicyclic peptides that inhibit Ahly mediated hemolysis using phage display. **a)** Chemical structure of Peptide 14 **b)** AlphaScreen assay for measuring monoclonal phage displayed bicyclic peptide binding to AviTag-Ahly. The assay was also performed in the presence of chemically synthesised bicyclic peptides to evaluate competition for binding sites (competition alpha screen), indicated by a decrease in phage AlphaScreen signal upon addition of free bicyclic peptide. **c)** AlphaScreen signal:background for phage clone expressing Peptide 14, displayed either as a 1,3,5-tris(bromoacetyl) hexahydro-1,3,5-triazine (TATB) scaffolded bicyclic peptide or a linear peptide with iodoacetamide capped cysteines. Values represent the ratio of signal in the presence of AviTag-Ahly to signal in the absence of AviTag-Ahly (streptavidin beads only). Mean of 3 technical replicates with SD error bars. **d)** Signal:background ratios for each phage clone in the presence of the corresponding chemically synthesised bicyclic peptide (10 µM). Values represent the ratio of signal in the presence of target to signal in the absence of target (streptavidin beads only). Heat map shading corresponds to the full range observed on the plate (2-1475): red indicates the lowest signal:background ratios (strong competition), and blue indicates the highest ratios (weak or no competition). **e)** Inhibition of Ahly (20nM) induced lysis of rabbit red blood cells by Peptide 14. Data are normalized to positive (Ahly only) and negative (PBS only) controls, DMSO concentration kept constant across all conditions. Mean of 3 technical replicates with SD error bars, data representative of 2 biological repeats.

**Table 1.**
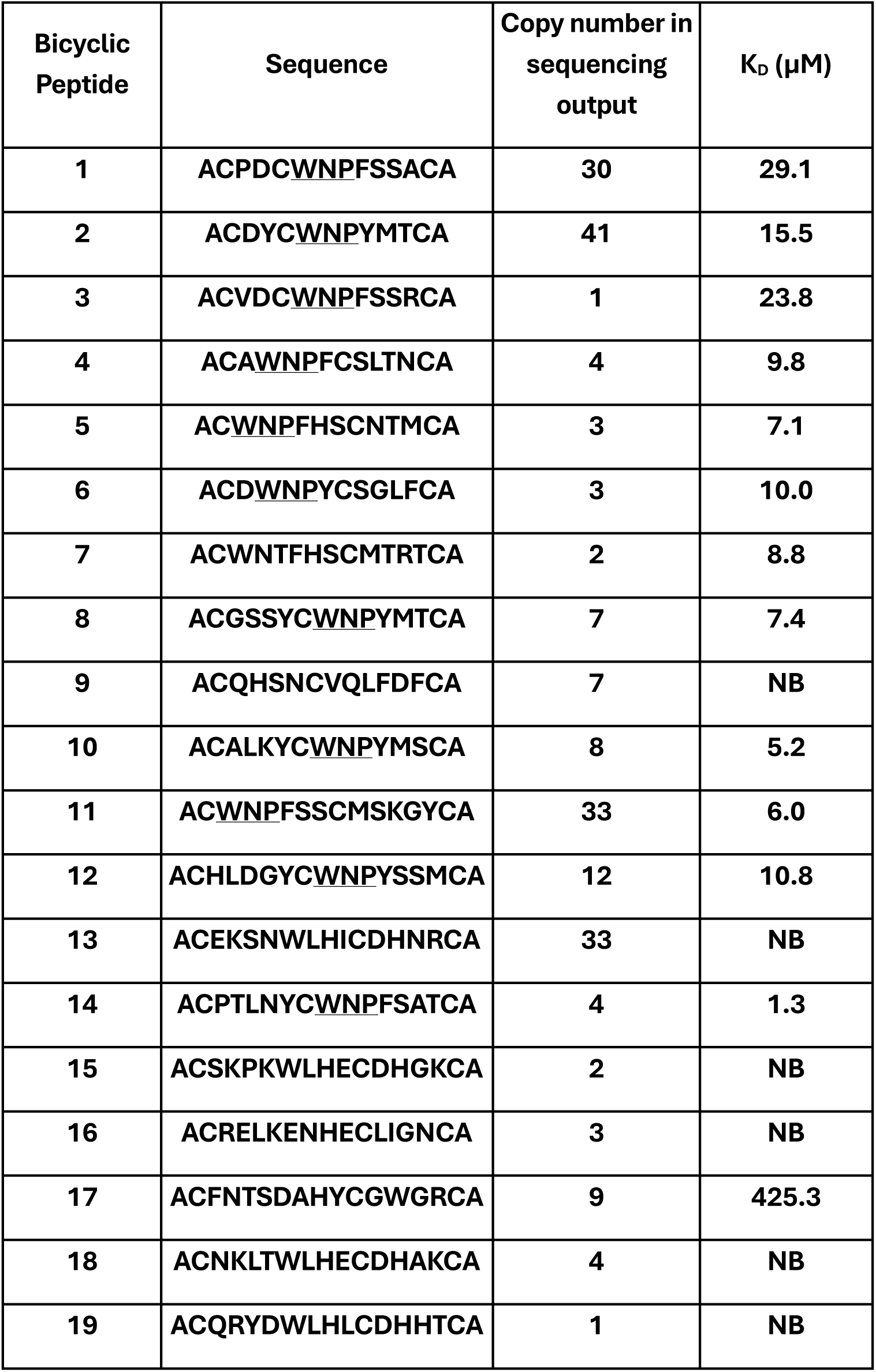
Bicyclic peptides identified from four rounds of selection against AviTag-Ahly. Peptide sequences chosen for chemical synthesis from sequencing output following four rounds of phage display selection using bicyclic peptide libraries scaffolded on 1,3,5-tris(bromoacetyl) hexahydro-1,3,5-triazine (TATB) against AviTag-Ahly. Conserved sequence motif is underlined. Binding affinities (K_D_) to AviTag-Ahly were determined by surface plasmon resonance (SPR) (n=1); NB = no detectable binding (compounds tested at 20μM maximum concentration).

**Figure 3.**
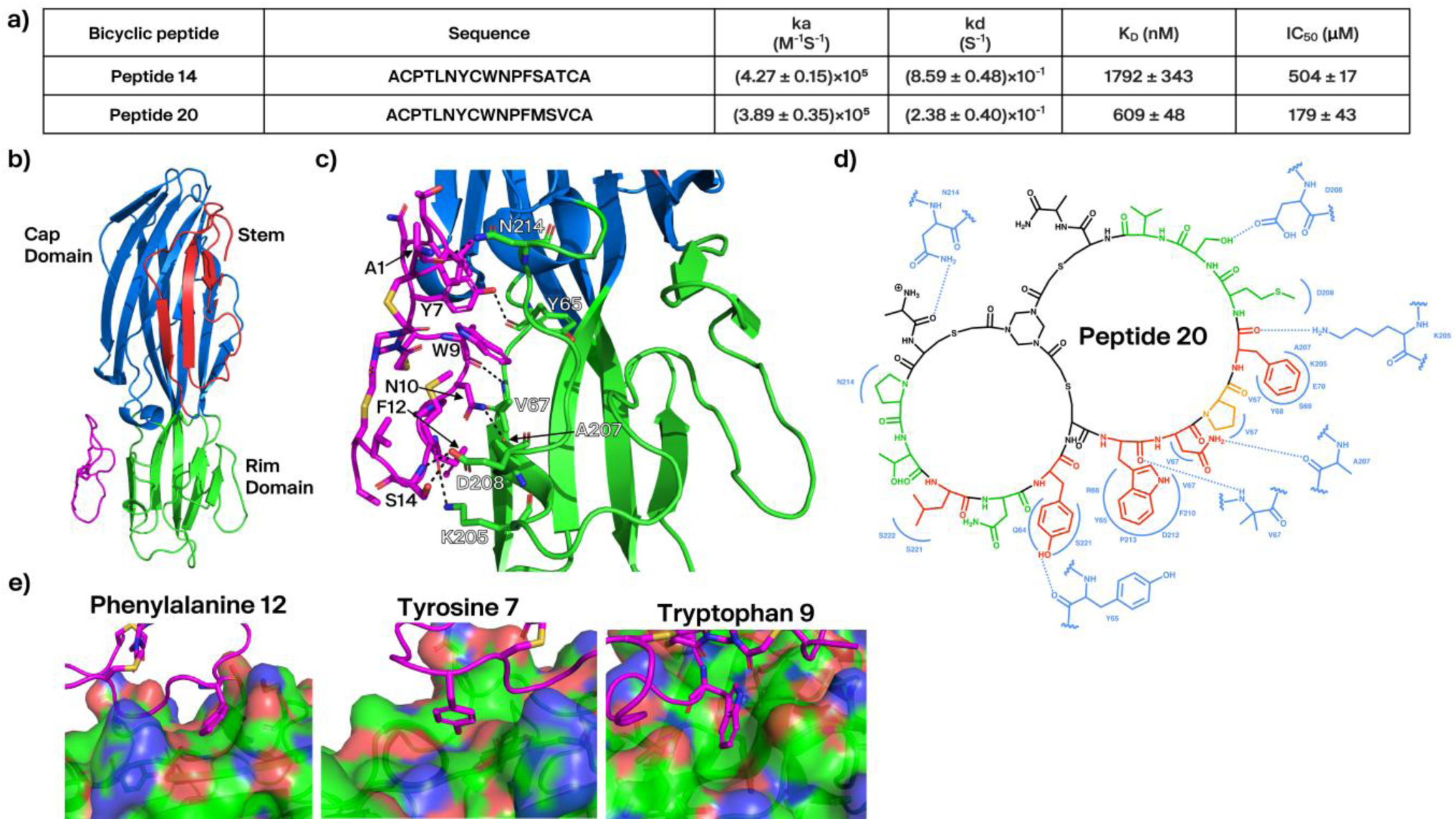
An X-ray crystallography co-crystal structure reveals structural insights into the interaction between Peptide 20 and AhlyH35A. **a)** Summary of binding kinetics determined via surface plasmon resonance (SPR) (mean ± standard deviation from ≥2 biological replicates) and hemolytic IC₅₀ values for the initial selection hit (Peptide 14) and the affinity matured bicyclic peptide (Peptide 20). **b)** X-ray crystallography co-crystal structure of Peptide 20 (magenta) bound to AhlyH35A **c)** Polar contacts between Peptide 20 and AhlyH35A depicted as dashed black lines, peptide residues are numbered from the N-terminal alanine. **d)** Representation of the binding interactions between Peptide 20 and AhlyH35A, as observed from the x-ray crystallography co-crystal structure, with blue indicating the interacting residues from AhlyH35A. Peptide 20 residues coloured based on their fold reduction in affinity to AviTag-Ahly (as determined by surface plasmon resonance) upon mutation to alanine; green, no reduction in affinity; orange, <5 fold reduction in affinity and red ώ5-no binding. **e)** Surface representation of AhlyH35A showing the positioning of aromatic residues from Peptide 20 at the binding interface, blue = positive charge, red = negative charge.

### X-ray crystallography co-crystal structure of affinity matured Peptide 14 in complex with Ahly monomer

Phage selections were used to optimise Peptide 14 in a process known as affinity maturation. Libraries were constructed consisting of phage expressing the peptide sequence of Peptide 14 or variants thereof, with a small number of residues in the peptide sequence randomised in each library. By creating multiple libraries, in which different residues are randomised at a time, we ensured broad sequence coverage and enabled testing of all amino acids at every position in the Peptide 14 sequence. Iterative rounds of selection against AviTag-Ahly using these libraries identified beneficial amino acid substitutions from the parent sequence that enhanced binding affinity to AviTag-Ahly. Sequences identified by affinity maturation were chemically synthesised and binding to AviTag-Ahly evaluated by SPR. This process produced an improved binder, Peptide 20, with a K_D_ of 609nM. The gain in affinity was driven by changes to the C-terminal residues in the peptide sequence. This increase in affinity also translated to greater inhibition of Ahly-mediated toxicity in a hemolysis assay (IC_50_, 179µM - Fig. 3a).

To gain insights into the binding interactions and the mechanism of inhibition, a co-crystal structure of Peptide 20 with an Ahly monomer was generated. A mutant Ahly was used containing the mutation H35A, to prevent the spontaneous formation of pores in crystallography conditions^62,63^. A co-crystal structure at 2.2Å resolution was produced of Peptide 20 bound to Ahly H35A, showing good electron density coverage for Peptide 20 (Fig. 3,S3, Table S1) (PDB: 9SVZ). The structure showed Peptide 20 bound to a slightly concave pocket on the rim domain of Ahly (Fig. 3a,S4). As the rim domain is responsible for the interaction with the phospholipid head groups of mammalian cell membranes, we proposed that Peptide 20 may exert an anti-hemolytic effect through inhibition of lipid interactions with Ahly^33,64^. Structural analysis revealed a mixture of polar and hydrophobic interactions, with the aromatic residues of Peptide 20 occupying distinct hydrophobic grooves on the surface of Ahly (Fig. 4d,e). The ‘WNP’ motif of Peptide 20 plays a central role in the binding interaction, with side chain amine of Asn10 forming a hydrogen bond with the carbonyl of Ala207 in Ahly. Pro11 appears to play a supporting role by introducing a kink into the peptide chain to allow Asn10 to protrude into Ahly, better positioning it to enable it to form a hydrogen bond (Fig. 3c, S5). Trp9 sits in a hydrophobic cleft at the centre of the binding footprint (Fig. 3e). Interestingly, the binding epitope of Peptide 20 closely resembles that of the antibody LTM14, also derived from phage display selections against Ahly^65^. Structural comparison revealed that both Peptide 20 and LTM14 engage Ahly in a strikingly similar manner, with LTM14 containing a ‘WRP’ motif that mirrors the ‘WNP’ sequence of Peptide 20. The two motifs align with an RMSD of 0.134Å across the Cα atoms of the three residues (Fig. S4). In LTM14, however, the arginine residue forms a salt bridge with Asp208, whereas in Peptide 20 the corresponding asparagine establishes a hydrogen bond with Ala207, despite the two residues occupying an almost identical binding pose on Ahly (Fig S6). Given that salt bridges generally provide stronger stabilization than hydrogen bonds, we attempted to substitute Asn10 with arginine to mimic the LTM14 interaction. The substitution abolished detectable binding, indicating that such an interaction cannot be recapitulated in Peptide 20 and that precise hydrogen bonding at this position is critical for binding of Peptide 20 to Ahly.

**Figure 4.**
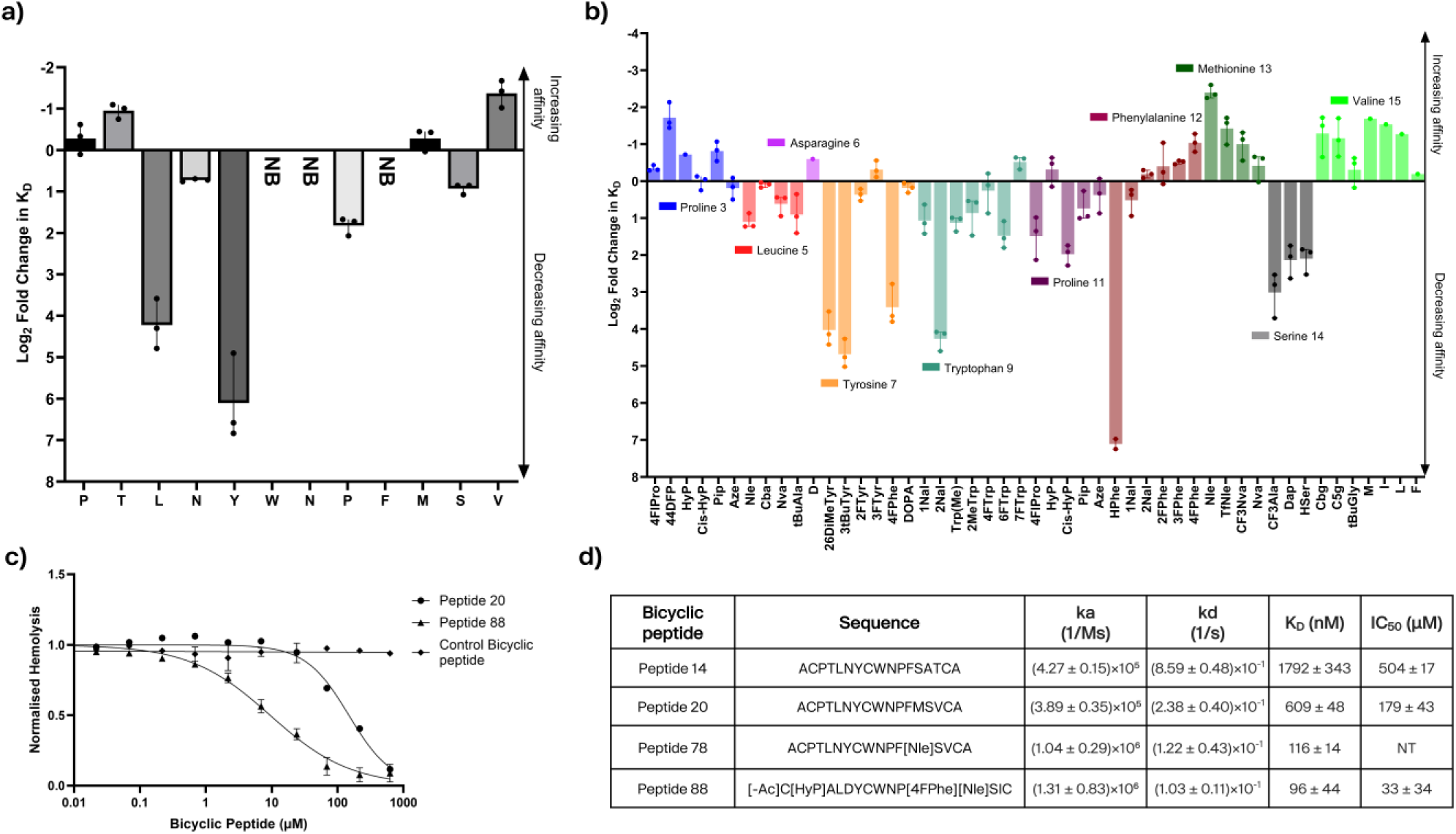
Effect of canonical and non-Canonical amino acid substitutions on Peptide 20 affinity and inhibitory potency. **a)** Fold change in binding affinity to Avitag-Ahly of Peptide 20 upon alanine substitution of individual residues, as measured by surface plasmon resonance (SPR). Data represent mean of three biological replicates; NB = no detectable binding (compounds tested at 2 μM maximum concentration). **b)** Fold change in binding affinity to Avitag-Ahly following substitution of selected residues in Peptide 20 with structurally similar non-canonical analogues or rationally chosen canonical amino acids. Data represent mean of three biological replicates. **c)** Inhibition of Ahly (30nM) induced lysis of rabbit red blood cells by Peptide 20 and Peptide 88, assessed in a hemolysis assay. Data are normalized to positive (Ahly only) and negative (PBS) controls, DMSO concentration kept constant across all conditions. Mean of three technical replicates with error bars indicating standard deviation (SD). **d)** Summary of binding kinetics as determined by SPR (mean ± SD from ≥2 biological replicates) and hemolytic IC₅₀ values (n=2) for the initial selection hit (Peptide 14), the affinity matured bicyclic peptide (Peptide 20), the non-canonical variant (Peptide 78), and the peptide incorporating multiple non-canonical residues (Peptide 88).

### Addition of non-canonical amino acids produced Peptide 88 with improved affinity and inhibitory potency

To understand the contribution of individual residues to target binding, we performed an alanine scan of Peptide 20 in which each residue in the peptide sequence was systematically substituted to an alanine (Figure 4a). As expected from sequence data and the co-crystal structure, the core ‘WNP’ binding motif could not be substituted without loss of binding. Replacement of Trp9 or Asn10 with alanine abolished any detectable binding, highlighting these residues’ crucial role in Ahly binding as demonstrated by the co-crystal structure. Phe12 to alanine resulted in no detectable binding and alanine in replacement of Leu5 and Tyr7 also significantly reducing binding affinities supported by the co-crystal structure which shows these residues engage in key hydrophobic interactions. The impact of ala substitutions at these positions demonstrated that they contribute markedly to Ahly binding, and underscored the importance of the hydrophobic interactions mediated by the aromatic residues of Peptide 20 (Fig. 4a, TableS2). In an effort to seek further gains in binding affinity, a panel of peptides incorporating a series of systematic non-canonical amino acid substitutions at positions within the peptide sequence of Peptide 20 were evaluated in SPR. These modifications were designed to explore whether introducing additional chemical diversity could further enhance binding affinity and thereby improve inhibitory potency as previously reported^66^. The full panel of non-canonical substitutions, including their structures and corresponding positions within the peptide sequence of Peptide 20 are shown in Fig. S7. The results showed most substitutions had either a negative or negligible effect on binding affinity compared to the parent Peptide 20 (Fig. 4b, Table S3). However, Peptide 68, with substitution of methionine with norleucine, had a 5.3-fold increase in affinity (K_D_ = 116nM). Truncation of the N- and C-terminal alanines, which are often sites of proteolytic degradation by exopeptidases, also showed a negligible effect on binding as did acetylation of the N-terminus (Table S4). A further panel of peptides was then synthesised, incorporating combinations of the tolerated changes in Peptide 20. These changes were chosen with the primary goal of improving binding affinity as well as increasing solubility and proteolytic stability. This set produced Peptide 88, which had the highest affinity to AviTag-Ahly of all bicyclic peptides tested (K_D_ = 96nM - Fig. 4d, Table S5). We observed a corresponding increase in inhibitory potency compared to Peptide 20 (IC_50_ = 179µM) with Peptide 88 displaying an IC_50_ = 33µM (Fig. 4c, 4d).

### Peptide 88 inhibits Ahly by blocking binding to cells

The co-crystal structure, together with the observed epitope similarities to the characterised antibody inhibitor LTM14, suggested that the inhibitory mechanism of Peptide 88 may be through blocking Ahly binding to cell membranes. To test this hypothesis, we performed a competition flow cytometry experiment in which we measured binding of Alexa Fluor 647-labelled Ahly H35A to A549 human alveolar epithelial cells in the presence of Peptide 88 or a control bicyclic peptide. A549 cells were chosen because they model human alveolar epithelium relevant to *S. aureus* lung infections and express robust levels of ADAM10 making them highly susceptible to toxin mediated damage^36^. The Ahly H35A mutant was used as this construct retains the ability to bind to cells but is incapable of oligomerising to form lytic pores. In the absence of bicyclic peptide we observed a clear shift in cellular Alexa Fluor 647 signal when A549 cells were incubated with fluorescent Ahly H35A, indicating binding of AhlyH35A to the cell membrane (Fig. S8a,b). Peptide 88, but not a control bicyclic peptide, was able to block this interaction in a dose dependent manner (Fig. 5a,b). These data suggested that, similar to LTM14, Peptide 88 inhibits Ahly by blocking binding to cells.

**Figure 5.**
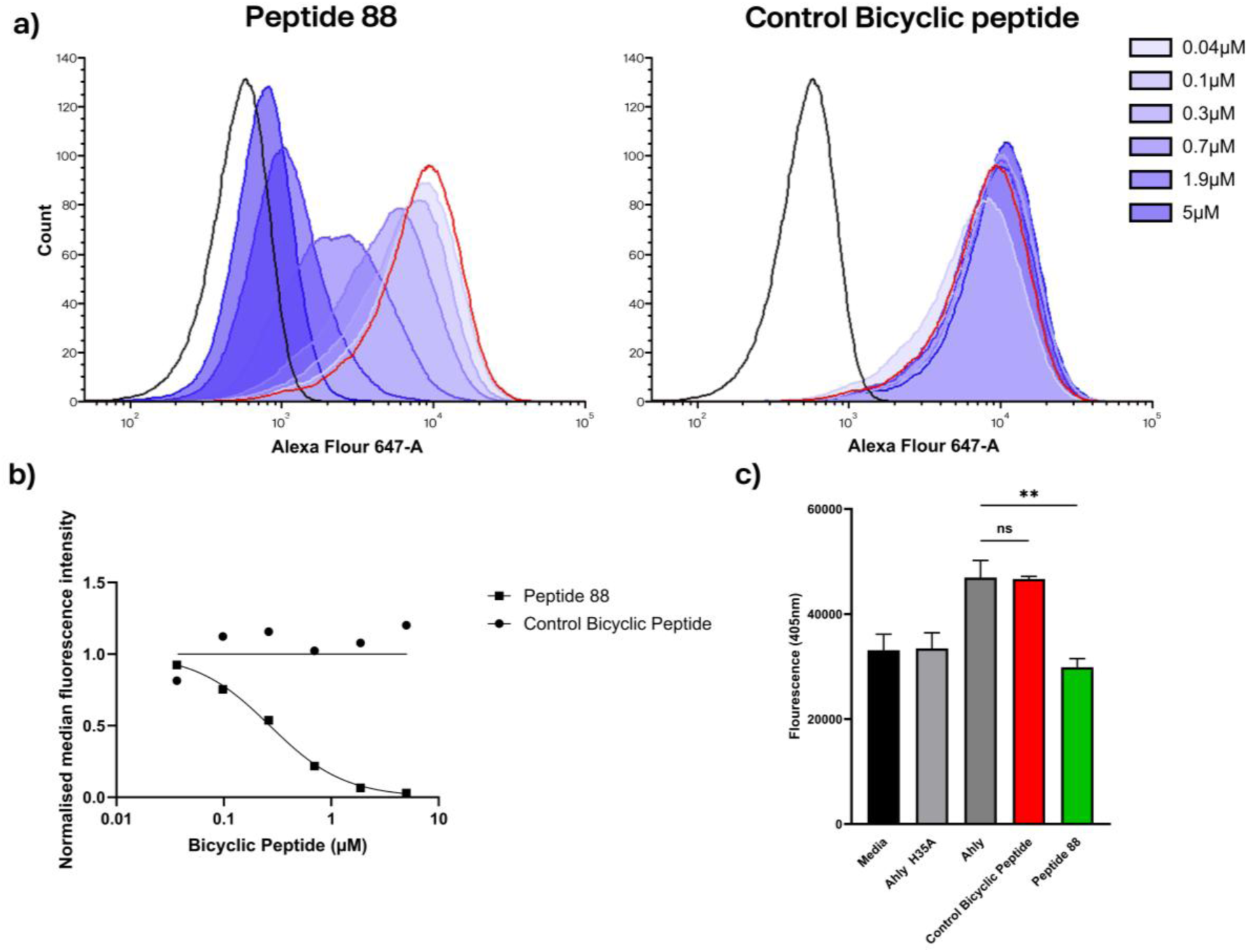
Peptide 17 inhibits Ahly binding to host cells. **a)** Alexa Fluor 647-labeled AhlyH35A (7.5nM) was incubated with A549 cells in the presence of increasing concentrations of Peptide 88 or a control bicyclic peptide. Cell-associated fluorescence was quantified by flow cytometry and shown as histogram overlays. Negative control (cells only) shown in black; positive control (AhlyH35A without peptide) shown in red. **b)** Quantification of median fluorescence intensity plotted against peptide concentration. Data are normalized to the negative and positive controls. **c)** ADAM10 protease activation by Ahly (6µM) was measured using a whole-cell FRET peptide cleavage assay in the presence of Peptide 88 or a control bicyclic peptide (900µM). Mean of two biological replicates; error bars indicate standard deviation. Data were analysed using one-way ANOVA with Dunnett’s test: ns = not significant; ** = P < 0.01.

Ahly has also been shown to upregulate ADAM10 metalloprotease activity on cells, which can be measured through the use of a fluorogenic peptide substrate^35,67,68^. Peptide 88, but not a control bicyclic peptide, was able to block the stimulating effect of Ahly on A549 cellular ADAM10 activity (Fig. 5c). Pore formation by Ahly underlies the stimulation of ADAM10, as demonstrated by the lack of ADAM10 stimulation by the non-pore forming mutant AhlyH35A (Fig. 5c)^35^. Therefore, inhibition of this stimulating effect can be caused by inhibiting either membrane binding or pore formation. Taken together we demonstrate that Peptide 88 has inhibitory effects on Ahly outside of pore formation, likely mediated through blocking cell binding.

### Peptide 88 protects A549 cells from Ahly induced cytotoxicity

Although hemolysis assays are commonly used to quantify the activity of Ahly inhibitors, they have limited relevance for evaluating therapeutic potential in humans, as Ahly exhibits minimal lytic activity toward human erthyrocytes^69^. To better model the cytotoxic effects of Ahly on human cells, we performed a cell-based assay using A549 cells. A549 cells were incubated with Ahly in the presence of Peptide 88 or a control bicyclic peptide, and cytotoxicity measured based on dead cell protease activity using a commercial reagent. Treatment with Ahly alone induced significant cell death, with no impact from the addition of a control bicyclic peptide. In contrast, Peptide 88 was able to abolish the toxic effects of Ahly on the A549 cells (Fig. 6a,b). We next investigated whether Peptide 88 could protect A549 cells from the cytotoxicity induced by co-incubation with a clinical isolate *S. aureus* strain 8325-4. Significant cytotoxicity was observed when A549 cells were co-cultured with *S. aureus* 8325-4, however a non-Ahly producing *S. aureus* strain (DU1090) was associated with negligible cytotoxicity (Fig 6c). The results showed Peptide 88, but not a control bicyclic peptide, statistically significantly inhibited 8325-4 toxicity in a dose dependent manner (Fig. 6d). Peptide 88 was not associated with *in vitro* toxicity to A549 cells at these concentrations (Fig. S9). Despite both assays producing a similar overall level of cytotoxicity (∼60%), higher concentrations of Peptide 88 were needed to produce an inhibitory effect in the co-infection assay in comparison to the recombinant Ahly assay.

**Figure 6.**
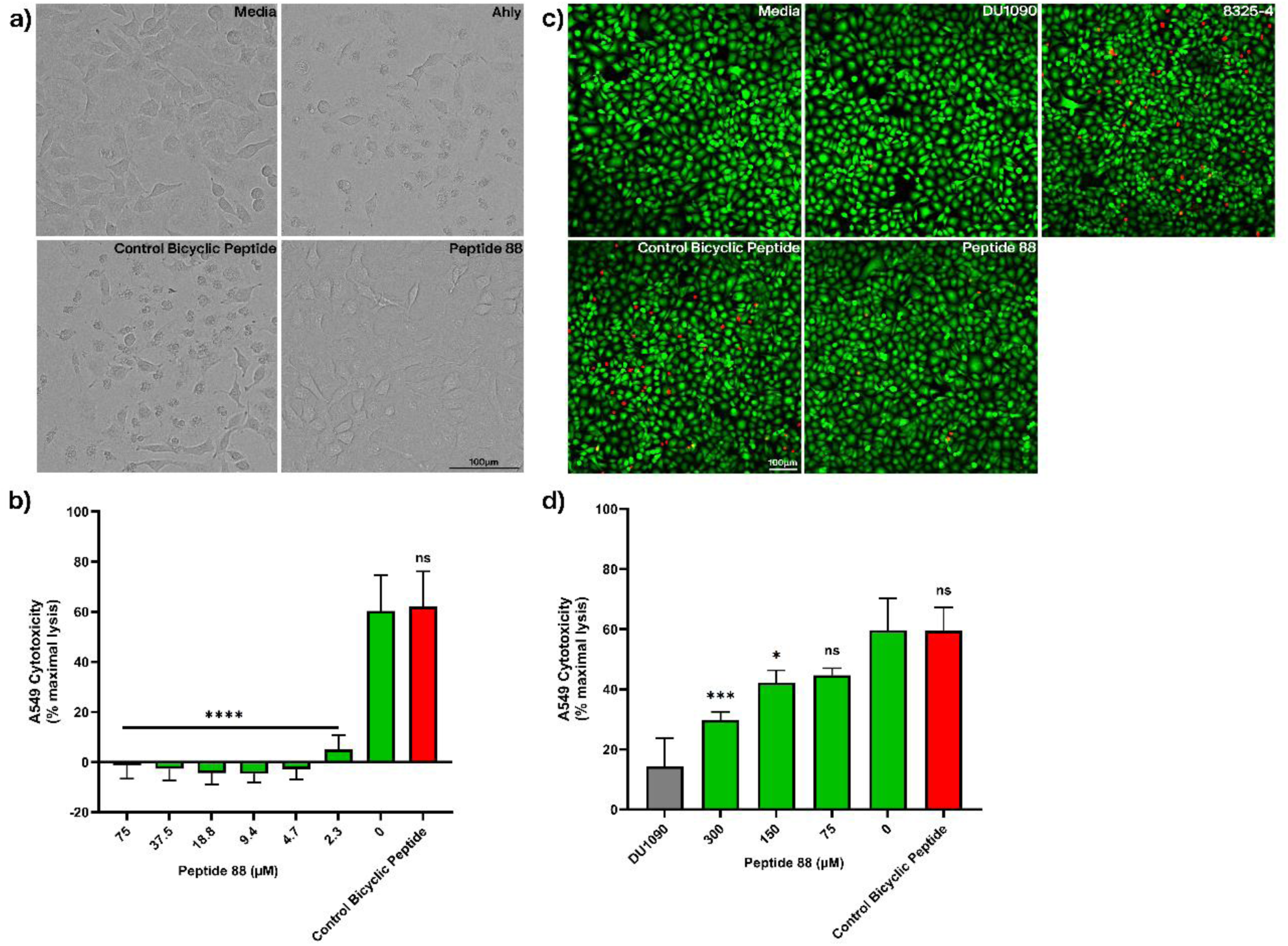
Peptide 88 protects A549 cells from Ahly mediated cytotoxicity. **a)** Representative brightfield images of A549 cells after 5-hour incubation with media, Ahly (1.5μM), or Ahly with Peptide 88 or a control bicyclic peptide (75μM). DMSO concentration was kept constant across all conditions. **b)** Cytotoxicity on A549 cells of Ahly (1.5μM) in the presence of indicated concentrations Peptide 88 or a control bicyclic peptide (75μM), measured after 5 hours by LDH release. Data are normalized to positive (1% Triton X-100) and negative (media) controls of cytotoxicity, DMSO concentration was constant across all conditions. Mean of three technical replicates with error bars indicating standard deviation (SD), data representative of 2 biological repeats. Data were analysed using one-way ANOVA with Dunnett’s test: ns = not significant; **** = P < 0.0001. **c)** Representative fluorescent microscopy images of A549 cells after 3.5 hour incubation with media, S. aureus 8325-4, or S. aureus 8325-4 with Peptide 88 or a control bicyclic peptide (300μM), the non-Ahly producing strain DU1090 used as a negative control for Ahly activity. DMSO concentration was kept constant across all conditions. Cells were stained using the live cell stain calcein-AM (green) and a dead cell stain ethidium homodimer-1 (red) before being imaged using confocal fluorescence microscopy. **d)** A549 cells were co-cultured for 4 hours with *S. aureus* 8325-4 in the presence of indicated concentrations of Peptide 88 or a control bicyclic peptide (300μM). The non-Ahly producing strain DU1090 served as a negative control for Ahly activity. LDH release was quantified as in panel b, data are normalized to positive (1% Triton X-100) and negative (vehicle only) controls of cytotoxicity. DMSO concentration was kept constant across all conditions. Mean of three technical replicates with SD error bars, data representative of 2 biological repeats. Data were analysed using one-way ANOVA with Dunnett’s test: ns = not significant:* = P < 0.05; *** = P < 0.001.

We hypothesised this loss of potency in the whole cell context may be due to degradation of Peptide 88 by proteases released from *S. aureus*. To investigate this, both Peptide 20 and Peptide 88 were incubated with *S. aureus* 8325-4 supernatant and subsequently analysed by liquid chromatography-mass spectrometry (LC-MS). Multiple hydrolysis products were detected for both Peptide 20 and Peptide 88 (Table 2, Fig. S11). Although accurate quantification of cleavage products could not be performed, the amount of cleavage products detected in relation to intact bicyclic peptide was minimal suggesting low levels of bicyclic peptide degradation (Fig. S11). No hydrolysis products were detected with M9 media only or when protease inhibitors were added to the *S. aureus* supernatant, suggesting that the bicyclic peptide degradation was protease dependent. To help identify specific cleavage sites, the alanine scan series of Peptide 20 was screened for susceptibility to proteolytic degradation in the same assay. Potential positions where alanine substitution showed no detectable cleavage sites may highlight key protease recognition or cleavage sites. Such data could enable rational design of protease resistant analogues by introducing tolerated alanine substitution or incorporating similar non-canonical amino acids at these critical positions in the hopes of circumventing degradation to enhance bicyclic peptide stability. The results revealed cleavage products were detected for all alanine substitutions indicating that multiple cleavage sites within the peptide sequence were present.

**Table 2.**
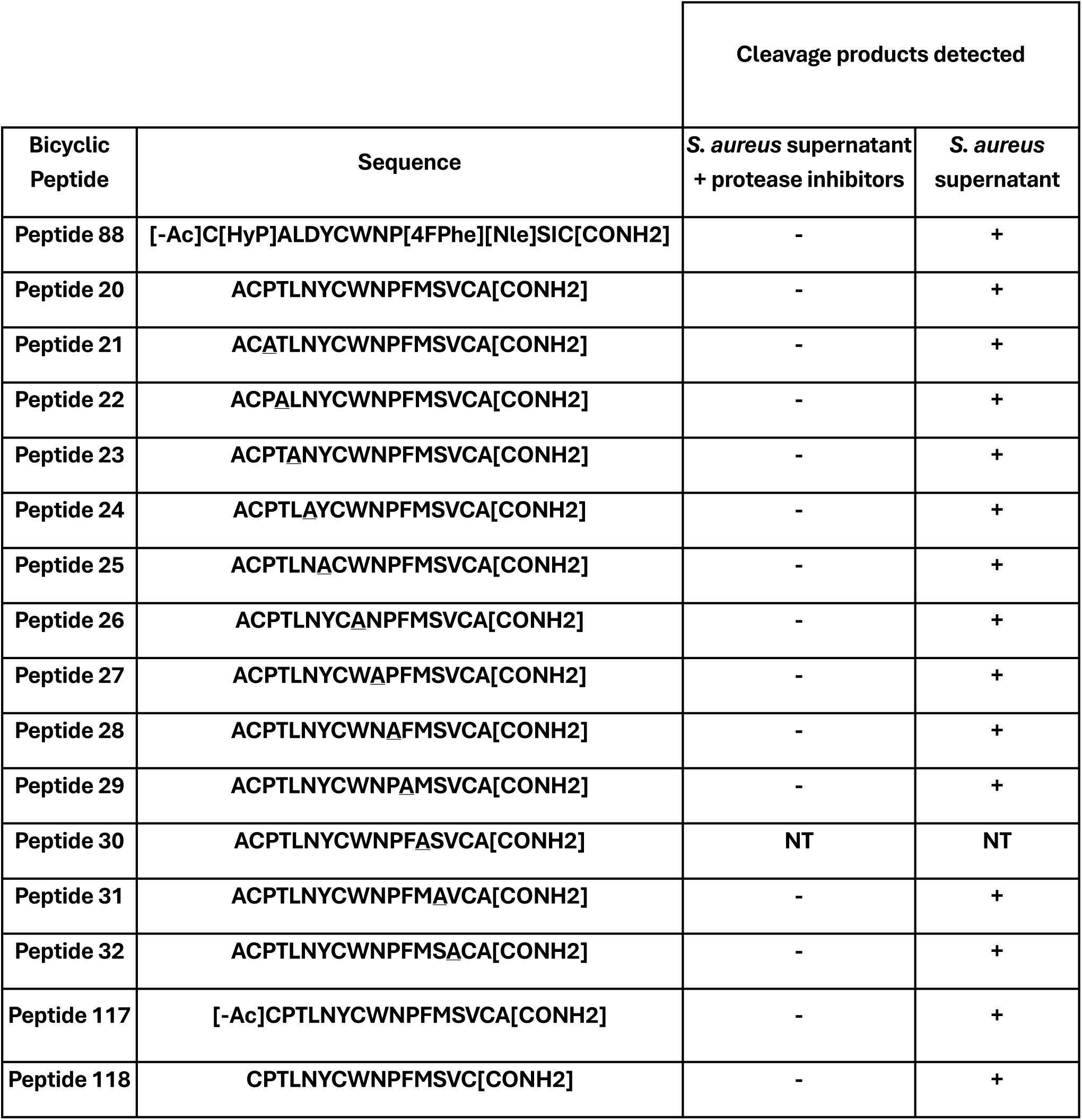
Proteolytic stability of bicyclic Peptide 20 and its alanine substituted variants after incubation with *S. aureus* supernatant. An alanine scan set of Peptide 20 was used to assess susceptibility to proteolytic cleavage by factors present in *S. aureus* 8325-4 culture supernatant. Each variant was incubated overnight with bacterial supernatant (grown in M9 minimal media) with or without protease inhibitors, and the presence of cleavage products detected by liquid chromatography mass spectrometry. Positions of alanine substitution are underlined.

An additional hypothesis for the reduced potency of peptide 88 in the whole cell context was that the high proportion of hydrophobic amino acids could result in the bicyclic peptide being sequestered in the membrane of *S. aureus,* thus reducing the effective concentration of peptide. Hydrophobicity has been shown to play an important role in antimicrobial peptides (AMPs) which associate with bacterial membranes^70–72^. We did not observe growth inhibition of *S. aureus* strain 8325-4 by peptide 88, suggesting that any membrane-associating effect was below the threshold for potent whole cell killing (Fig. S10).

## Discussion

To date, antibody Ahly inhibitors have been the sole modality to have progressed into clinical trials as anti-virulence *S. aureus* therapies. These trials focused on prophylaxis for those at risk or severe infections such as *S. aureus* pneumonia in hospitalised patients ^73,74^. Here, we show that through phage display we can rapidly identify bicyclic peptides which offer a potential alternative modality for systemic therapy. The peptides presented show binding to Ahly and inhibition of the pore-forming function of the toxin. Further work in the development of these peptides for systemic delivery would include optimising their drug-like properties, such as blood stability and exposure following IV administration; and evaluating their efficacy in mouse models of infection by *S. aureus*. Pre-clinical studies of anti-Ahly antibody efficacy in mouse have been performed using readouts such as lung CFU and lesion size following pneumonia and dermonecrosis models, respectively^75,76^ The efficacy of our peptides will be based upon a combination of the target engagement (i.e. the ability to inhibit pore formation by Ahly) and drug-like properties, and so further work could also encompass additional enhancement of the inhibitory potency of our peptides to the low nanomolar – picomolar K_D_ range reported with antibody inhibitors^65,75,77,78^. In particular, we have shown proteolytic instability of the peptides presented here. Enhancement of protease stability, guided by metabolite identification (metID) studies in *S. aureus* culture and relevant biological matrices (such as whole blood) could result in both improved inhibitory potency in co-culture and exposure upon systemic dosing.

While the development of therapies for life threatening *S. aureus* infections remains essential, there is also a pressing need for treatments that address more prevalent, less severe infections. In this regard, *S. aureus* is the leading cause of skin and soft tissue infections (SSTIs) in the United States^79^. One such example is the superficial bacterial skin infection impetigo which is estimated to effect in excess of 162 million children worldwide at any given time^80^. *S. aureus* is one of the primary causes of impetigo and a Cochrane review indicated that topical antibiotics are equally, or more, effective than oral antibacterials in the treatment of impetigo^81^. A stable and easily manufactured anti-Ahly therapy to apply topically to surface skin infections such as impetigo in conjunction to topical antibiotics could be highly advantageous. Bicyclic peptides could be such a modality, combining facile solid phase peptide synthesis with enhanced stability conferred by their rigid chemical scaffold and a reduced manufacturing cost in comparison to monoclonal antibodies.

Bicyclic peptide phage display selections against Ahly yielded a striking convergence on a core ‘WNP’ motif present in most TATB-scaffolded phage clones after 4 rounds of selection. Binding studies confirmed that only bicyclic peptides containing this motif exhibited detectable binding in competition AlphaScreen or SPR. Furthermore, substitution of either Trp9 or Asn10 in this motif to alanine abolished the interaction, underscoring the critical role of the motif in binding to Ahly. Cyclisation was a pre-requisite to target engagement as Peptide 14 exhibited binding to Ahly in AlphaScreen only when constrained into a bicyclic peptide with no binding detected when left in a linear format. Cyclic peptides have a high degree of structural preorganisation into stable conformations that enable high affinity binding. Linear peptides, in contrast, often fail to adopt such stabilized conformations due to their inherent flexibility making such binding entropically unfavourable. The essentiality of the ‘WNP’ motif was further underscored by structural data, which revealed that Asn10 forms a crucial hydrogen bond and Pro11 contorts the peptide back bone to support this interaction with the target. Further interactions were formed by the non-substitutable aromatic residues Trp9, Tyr7, and Phe12, which were embedded in the hydrophobic cleft at the centre of the binding interface on Ahly. Intriguingly, independent antibody selections to Ahly utilising a human-derived phage display library produced LTM14, an Ahly inhibiting antibody possessing a similar ‘WRP’ motif within its paratope. Alignment of the co-crystal structures from both studies revealed that this shared motif engages Ahly through an almost identical binding pose and a similar set of intermolecular interactions. This remarkable convergence across distinct molecular scaffolds (cyclic peptides and antibodies) suggests a highly privileged interaction site on Ahly. Using flow cytometry, we showed that despite a far smaller molecular footprint than LTM14, Peptide 88 was able to inhibit the pore forming function of Ahly through the same inhibitory mechanism, by blocking Ahly membrane binding.

An extensive campaign to improve the binding affinity of our initial hit Peptide 14 was undertaken with the affinity to Ahly improving twenty-fold from 1792nM to 96nM. Peptide 20 was identified through affinity maturation, and further improved by designing, synthesising and testing bicyclic peptides containing non-canonical amino acid substitutions. Both affinity maturation and incorporation of non-canonical amino acids improved affinity from position 13, substituted from serine in the phage clone to methionine, and ultimately norleucine. Structural analysis shows that the methionine side chain sits between the electron rich TATB scaffold and a hydrophobic surface on Ahly. Replacing the sulphur atom in methionine with a less electron rich carbon, as in norleucine, may reduce repulsion forces from the scaffold while enhancing hydrophobic interactions with the protein. This process of optimisation resulted in Peptide 88 with an affinity to Ahly of 96nM and an IC_50_ of 33µM. These results serve to highlight both the substantial gains in binding affinity and inhibitory potency achievable through peptide optimisation, and the broad range of tools and methods available to accomplish it.

Peptide 88 showed potent protection of A549 cells from recombinant Ahly at close to equimolar concentrations to the toxin. This protection was also seen in the whole cell context with Peptide 88 inhibiting *S. aureus* 8325-4 toxicity when co-cultured with A549 cells, albeit at higher concentrations. We initially proposed and investigated 2 hypotheses to explain this asymmetry in potency between the two assay. First, the high proportion of hydrophobic residues in Peptide 88 may lead to sequestration within *S. aureus* membranes, reducing its availability to bind to Ahly. Second, proteases released by *S. aureus* may degrade Peptide 88. Whilst the addition of Peptide 88 had no effect on the growth of *S. aureus* strain 8325-4 this does not rule out membrane sequestration, as it may occur without sufficient membrane disruption to impact bacterial proliferation. Therefore, we cannot exclude bacterial membrane association of Peptide 88. However, this would be surprising given Peptide 88 lacks the characteristic cationic or amphipathic properties typically associated with bacterial membrane partitioning AMPs. To assess proteolytic degradation, Peptide 88 was incubated with *S. aureus* 8325-4 supernatant, after which LC-MS analysis showed clear evidence of peptide cleavage albeit at low levels and thus unlikely to completely account for the loss in potency. Systematic alanine substitution was able to indicate potential key residues that are susceptible to proteolysis namely Leu5, Try9 and Phe12. However, this enhanced protease stability comes with a trade off as alanine at these positions either significantly reduces or abolishes binding to Ahly. Rather, strategies to mitigate this degradation should focus on the use of non-canonical amino acids, N-methylation, peptoid residues or the use of D-amino acids to name a few.

In summary, this study presents bicyclic peptides, identified via phage display, as a novel class of Ahly inhibitors that target a conserved and functionally important epitope on Ahly. Through affinity maturation, structural analysis and functional assays we developed our initial hit into our lead bicyclic peptide with nanomolar affinity, validated binding site and mechanism of action as well as demonstrated its capability of protecting human cells from toxicity induced by *S. aureus* infection. These findings highlight the value and potential of bicyclic peptides as novel therapeutic modalities for the development of next generation anti-virulence therapeutics.

### Experimental Section

#### Expression and Purification of Proteins

The gene encoding Ahly was purchased from IDT with a N- 6x histidine tag followed by a 3C protease cleavage tag to facilitate tag removal. The gene also contained NdeI and XhoI restriction enzyme sites at the N- and C-terminus respectively. The gene was cloned into a pET47b vector using NdeI and XhoI restriction digest following manufacturer instructions (New England Biolabs). Plasmids were transformed into chemically competent BL21 (DE3) cells, a single colony added to 100mL of lysogeny broth (LB) supplemented with Kanamycin (50 µg/ml) and grown overnight at 37°C with shaking at 180rpm. Afterwards, 10-15mL of overnight culture was diluted into 1L of Terrific Broth (TB) supplemented with Kanamycin at 50 µg/ml for a total of 6L and incubated 37°C 180rpm. Once the culture reached optical density of 600nm (OD_600_) of 0.6-0.8, Isopropyl-β-D-thiogalactopyranoside (IPTG) was added to a final concentration of 1mM after which the temperature was dropped to 20°C for overnight incubation. The following day bacteria were pelleted at 3,500xg for 20 minutes and the pellets flash frozen in liquid nitrogen and stored at -80°C. For purification, pellets were resuspended in 100ml Buffer A (30mM HEPES, 150mM NaCl, 1.5mM TCEP, 2% Glycerol (w/v), pH 7.4) with the addition of cOmplete™ EDTA-free Protease Inhibitor tablets (Roche) on ice. Cells were then lysed either using sonication or a cell disruptor and cell debris pelleted by centrifugation at 40,000xg for 1h. Cell lysate was then loaded onto a HisTrap HP column (Cytiva) pre-equilibrated with buffer A and eluted with a gradient of 0-100% buffer B (Buffer A + 500mM imidazole). Fractions containing protein (as determined by absorbance at 280nm) were pooled and dialysed overnight with stirring at 4°C in dialysis tubing (3kDa cutoff, Fischer) suspended in 2L buffer A with his-tagged 3C protease added for tag cleavage. The His-tag cleaved protein was then further purified by reverse IMAC on a HisTrap HP column (Cytiva) using buffer A and B. Protein containing fractions were then pooled and concentrated on a 3kDa molecular weight cut off centrifugal concentrator (Vivaspin) until a volume <5ml before being purified further via size exclusion chromatography (Superdex 16/60 75pg, Cytiva) in buffer A. Finally, protein was pooled and dialysed overnight against 2L storage buffer (30mM HEPES, 150mM NaCl, 2mM TCEP, pH 7.6, 50% glycerol (w/v)), aliquoted and flash frozen in liquid nitrogen and stored at -80°C prior to use.

For AviTag proteins, constructs were transformed into chemically competent BL21 (DE3) cells pre-transformed with a BirA expression vector. Expression and purification were performed as described previously, with the exception of TB media supplementation with 700µl of 0.1mg/ml D-biotin.

AhlyH35A was expressed and purified as described above and labelled with AF647 using Alexa Fluor™ 647 NHS Ester (Thermo Fischer) following manufacturer’s instructions, unreacted AF647 was removed further via size exclusion chromatography (Superdex 16/60 75pg, Cytiva).

#### Selection of bicyclic peptides via phage display

Selections were performed as described previously^59^. Briefly, AviTag biotinylated Ahly was used as the input to phage display selections. After incubation with the phage library, Ahly was removed using magnetic streptavidin beads, washed to remove non-specific binding phage and Ahly bound phage eluted. Eluted phage were amplified overnight in TG1 *E. coli,* purified and their peptide sequences cyclised by reacting the trivalent chemical scaffold with the 3 fixed cysteines present in each sequence, prior to the subsequent selection round. After 4 rounds of selection using a decreasing concentration of Ahly at each round, the eluted phage were allowed to infect TG1 *E. coli* for 1 hour before being plated on tetracycline agar plates (12.5 mg/ml). The following day, colonies were selected for phage plasmid sanger sequencing to determine the peptide sequences expressed on the phage. Individual phage clones were then assessed in their ability to bind Avitag-Ahly using an AlphaScreen assay. Select binders were made individually by solid phase peptide synthesis for further characterization.

#### Bicyclic peptide synthesis and purification

All peptides were synthesized on Rink amide resin using standard Fmoc (9-fluorenylmethyloxycarbonyl) solid phase peptide synthesis using 2 automated systems. Peptide synthesis at 25μmol was run on a Biotage SyroII automated synthesizer. Peptide synthesis (80–240 μmol) was carried out with a Gyros Symphony X automated synthesizer. Bicyclic molecules were synthesized with a C-terminal lysine(pentynoyl) for use in copper(I)-catalysed azide alkyne cycloaddition (CuAAC). Following cleavage from the resin using a cocktail of 95% TFA, 2.5% triisopropylsilane, and 2.5% H2O with 25 mg of dithiothreitol (DTT) per mL, peptides were precipitated with diethyl ether and dissolved in 50:50 acetonitrile/water. Linear vector peptides were lyophilized after cleavage. Peptides for bicyclization were diluted to 2 mM in 50:50 acetonitrile:water, 2.6 mM scaffold solution, and 200 mM ammonium bicarbonate to give final concentrations of 1, 1.3, and 100 mM respectively. Completion of cyclization was determined by matrix assisted laser desorption ionization time-of-flight (MALDI-TOF) or LC–MS. Once complete, the cyclization reaction was quenched using N-acetyl cysteine (10 equiv of 1 M solution over peptide) and lyophilized. Standard Fmoc amino acids, as well as nonproteinogenic Fmoc amino acids, were obtained from Sigma-Aldrich, Iris Biotech GmbH, Apollo Scientific, ChemImpex and Fluorochem.

Crude peptides, following lyophilization, were dissolved in an appropriate solvent system and filtered through a 0.45 μm PES filter before loading onto a 5 μm, 100 Å, 21.2 × 100 mm Kinetex XB-C18 column (Phenomenex). Prep HPLC gradients using 0.1% TFA in H2O (solvent A) and 0.1% TFA in acetonitrile (solvent B) were selected based on the retention time of samples analysed either after cleavage or during cyclization on a 2.6 μm, 100 Å, 2.1 × 50 mm Kinetex XB-C18 analytical column on a gradient of 5–95% 0.1% TFA in acetonitrile.

Peptide fractions of sufficient purity and correct molecular weight (verified MALDI-TOF and HPLC or LC–MS) were pooled and lyophilized. Concentrations were determined by UV absorption using the extinction coefficient at 280 nm, which was based on Trp/Tyr content.

#### Antibody AlphaScreen

Antibody binding to Ahly was assessed using an AlphaScreen assay. AviTag-Ahly (25 nM) was added to wells of a 384-well OptiPlate, followed by addition of increasing concentrations of suvratoxumab in assay buffer (30mM HEPES, 150mM NaCl, 1.5mM TCEP, pH 7.4). Under subdued lighting, 20 µg/mL streptavidin-coated donor beads (PerkinElmer) and 20 µg/mL acceptor beads conjugated to an anti-IgG antibody (PerkinElmer) were added to each well. The plate was incubated for 1 h at room temperature before being read on a CLARIOstar plate reader (BMG Labtech) with excitation at 680 nm and emission measured at 570 nm.

#### Hemolysis assay

Ahly was preincubated with serial dilutions of bicyclic peptide for 30 minutes in a final volume of 50µl in a v-bottom 96 well plate (Corning). After which, 50µl of 5% (v/v) ice cold PBS washed rabbit erythrocytes were then added and the plate incubated at 37°C for 1h. Following incubation, 100µl of PBS was added to all wells and the plate was centrifuged at 1000xg for 20 minutes. Finally, 100µl of supernatant from each well was transferred to a flat bottom 96 well plate (Corning) and the amount of haemoglobin released was measured by absorbance at 520nm in a CLARIOstar plate reader (BMG Labtech). Data was normalised to a positive control of Ahly only and a negative control of PBS only, data plotted and fitted in GraphPad Prism 10 to generate IC_50_ values. Reported IC_50_ values represent an average of 2 biological replicates.

#### X-ray crystallography

The plasmid encoding Ahly H35A was produced from the original WT Ahly plasmid via site directed mutagenesis (New England Biolabs) and expressed and purified via the same method as described previously. The protein was concentrated to 10mg/ml before being aliquoted and flash frozen in liquid nitrogen. An aliquot of protein was thawed and Peptide 20 added at a 3:1 molar ratio of bicyclic peptide to protein. After a 30 minute incubation on ice the protein and bicyclic peptide mixture was subject to sitting-drop crystallisation screening trials at 290K against 6 commercial screens in 96 well crystallography plates using a Formulatrix NT8. Three different ratios of protein to crystallisation condition (2:1, 1:1 and 1:0.5) were used for each condition in the screens. Crystals appeared in a 1:0.5 ratio of protein (30mM HEPES pH 7.6, 150mM NaCl and 1mM TCEP) to crystallisation buffer (0.2M calcium chloride dihydrate, 0.1M sodium acetate and 20% w/v PEG6000). Crystals were cryoprotected in crystallisation buffer + 20% glycerol (v/v) and flash frozen in liquid nitrogen. Diffraction data was collected from the diamond synchrotron in Oxford, United Kingdom and analysed with AIMLESS, PHASER, Coot and refmac5. The Ahly H35A structure PDB code: 4YHD was used as the model for molecular replacement. Structural figures were made in PyMol and ChemDraw.

#### Surface plasmon resonance (SPR)

The affinity and rates of association and dissociation between bicyclic peptides and Ahly were measured using either a BIAcore T200 or a BIAcore 8K (Cytiva). Avitag-Ahly was immobilised by flowing the protein in PBSP+ (0.2 M phosphate buffer with 27 mM KCl, 1.37 M NaCl and 0.5% Tween 20 (v/v), Cytiva) over a streptavidin sensor chip (Cytiva). Bicyclic peptides were dispensed into a 384 well plate using a D300e compound dispenser (Tecan) before being diluted in PBSP+ spanning a typical concentration range 10µM-0.04µM. DMSO was normalised ensuring all wells contained 1% (v/v) DMSO final concentration. Bicyclic peptides at each concentration were flowed over the sensor as well as a control streptavidin chip surface with no immobilised protein as a control in PBSP+, 1% DMSO. The signal from the streptavidin-only control surface was subtracted from the signal from the protein surface to produce the final sensorgrams. Affinity constants were estimated by curve fitting of sensorgram plots using a 1:1 binding model in BIAcore evaluation software. Reported K_D_ values generally represent an average of 3 biological replicates, unless otherwise stated.

#### Flow cytometry

A549 cells at 1×10^6^ cells/ml were incubated with 7.5nM of AF647 labelled Ahly H35A for 2 hours in the presence or absence of bicyclic peptides in PBS at 4°C. Control wells of cells only and cells + AF647 Ahly H35A only were included. Cells were then washed 3 times in PBS to remove any unbound protein before staining of dead cells with DAPI at 1µg/ml prior to analysis by flow cytometry using a an Attune NxT flow cytometer (Fischer) at excitation at 650nM and emission at 671nM. Data was analysed using FCS express software and median fluorescent intensities plotted in GraphPad Prism 10 normalised to cell only negative and cells + Ahly H35A only positive controls. The results are representative of three independent experiments.

#### ADAM10 activity assay

A549 cells were seeded at a density of 1-2 x 10^4^ cells per well in a tissue culture treated 96 well plate (Corning) and incubated overnight at 37°C with 5% CO_2_. On the following day, bicyclic peptides were pre-incubated with Ahly (6µM) in cell culture medium for 10 minutes at room temperature. The culture medium was aspirated from each well and replaced with 100µl of the prepared Ahly-bicyclic peptide mixture. Cells were incubated for 15 minutes at 37°C with 5% CO_2_. Following incubation, cells were washed once with 25mM Tris buffer, pH 8.0 and a fluorogenic ADAM10 substrate peptide (Mca-PLAQAV-Dpa-RSSSR-NH_2_; R&D Systems) was added at a final concentration of 10µM. Plates were incubated for an additional 15 minutes under the same conditions. Substrate cleavage was then quantified using a CLARIOstar plate reader (BMG Labtech) with excitation at 320nm and emission at 405nm.

#### Ahly Cytotoxicity assay

A549 cells were seeded at a density of 1-2 x 10^4^ cells per well in tissue culture treated 96 well plates (Corning) and incubated overnight at 37°C with 5% CO_2._ The following day, serial dilutions of bicyclic peptides were prepared and incubated with 1.25µM Ahly for 30 minutes in cell culture medium (DMEM + 10% FBS (v/v)) at room temperature. Subsequently, the cell culture medium from each well was aspirated and replaced with 100µl of the prepared peptide-Ahly mixtures. The plate was incubated for 5 hours at 37°C with 5% CO_2._ After incubation, brightfield images were taken at 40x magnification using a Cytation 7 microscope (BioTek). Cytotoxicity was quantified using the CytoTox-Glo cytotoxicity assay (Promega) according to the manufacturer’s instructions. Luminescence was measured using a luminometer plate reader (Promega). Data were normalised to 100% for a positive control of cells treated with Ahly only and 0% for a negative control of cells treated with medium only. The results are representative of two independent experiments.

#### Bicyclic peptide cytotoxicity assay

A549 cells were seeded at 1×10^4^ cells per well and incubated overnight at 37°C before the addition of Peptide 88 and incubated together for 72h. Following this, cytotoxicity was measured using the CellTiter-Glo® cell viability assay (Promega) following the manufacturer’s instructions. Cytotoxicity was normalised to a positive control of 1% (v/v) Triton X-100 and a negative control of DMSO only.

1. *S. aureus* Cytotoxicity assay.

A549 cells were seeded at a density of 1-2 x 10^4^ cells per well in tissue culture treated 96 well plates and incubated overnight at 37°C with 5% CO_2_. On the following day, *S. aureus* strain 8325-4 was cultured in tryptic soy broth to an OD_600_ of 0.5. A 5ml aliquot of the culture was washed with PBS and made up to a final volume of 10ml in cell culture medium (DMEM + 10% FBS (v/v)). Serial dilutions of Peptide 88 were prepared in this bacterial suspension and incubated together for 30 minutes. The cell culture medium was aspirated from each well and replaced with 100µl of the peptide-*S. aureus* mixture. Plates were incubated for 4 hours at 37°C with 5% CO_2_. Cytotoxicity was then measured using the CyQUANT™ LDH cytotoxicity assay (ThermoFisher) according to the manufacturer’s instructions, or cells were stained using a LIVE/DEAD stain (Proteintech) and imaged on a Zeiss LSM 880 confocal microscope. The non-Ahly producing strain DU1090 was used as an Ahly negative control. Data was normalised to 100% for a cytotoxicity positive control of cells treated 2% (v/v) Triton X-100 and 0% for a negative control of cells treated with medium only. The results are representative of two independent experiments.

#### Bicyclic peptide cleavage assay

*S. aureus* 8325-4 was grown in M9 minimal media at 37°C until an OD of 1.5, after which the supernatant was removed and filtered twice through a 0.22µM filter (Merck) to remove any remaining *S.aureus.* Bicyclic peptides were added to the supernatant with or without protease inhibitors (Merck) and EDTA (0.124% (w/v)) at 256 µg/ml (123µM) in a final volume of 600µl before incubation at 37°C overnight and flash frozen in liquid nitrogen the following day. Samples were analysed by Waters TQD LC-MS on a run using a gradient of 5-95% over 9 minutes on a 2.6 µm, 100 Å, 2.1×30 mm Kinetex XB-C18 analytical column in 0.1% formic acid in water (solvent A) and 0.1% formic acid in acetonitrile (solvent B). The run was performed at a temperature of 21°C and flow rate of 0.6 mL/min. The results were analysed at 220 nm and peak identity was confirmed using the correlating mass spectra for each peak. The number of cut sites were generated by observing the number of peaks with the mass of (M+18). For an example, with 2 cut sites another peak was also observed with a mass of [M+2(18)]. Peaks were integrated in relation to base line absorbance and the total area under the peaks corresponding to hydrolysis products was used in comparison to the area under the peak corresponding to the intact peptide to calculate the percentage hydrolysis.

#### MICs and bacterial growth curves

Minimal inhibition concentration (MIC) assays were performed using S. aureus 8325-4 in accordance with CLSI standards, using cation-adjusted Mueller Hinton broth (caMHB; Sigma Aldrich). For bacterial growth curve monitoring, cultures were prepared as for MIC assays and OD_600_ was measured constantly over time in a CLARIOstar plate reader (BMG Labtech), with incubation at 37°C.

## Acknowledgments

This work was by funded by a studentship from the MRC Doctoral Training Partnership in Interdisciplinary Biomedical Research at the University of Warwick (grant number MR/R015910/1) and Bicycle Therapeutics. We thank Professor David Roper for supplying the chemically competent BL21 (DE3) cells pre-transformed with a BirA expression vector. The media preparation team of Cerith Harries and Charlotte Curtis at the University of Warwick and the lab support team at Bicycle Therapeutics for producing all the agar plates, bacterial media, PBS and antibiotic stocks used in this work. The biophysics team at Bicycle Therapeutics for all their help and guidance with SPR as well as the chemistry team for producing the bicyclic peptides that made this work possible.. The *S. aureus* strains 8325-4 and DU1090 were kindly donated by Professor Simon Foster. Finally, many thanks to Valentin Dospinescu, Paulina Michór and Jack Butler for their generous hospitality.

## Supporting Information

**Figure S1:** C-terminal AviTag fusion does not perturb Ahly structure or function

**Figure S2:** Peptide 14 is a specific inhibitor of Ahly

**Table S1:** Statistics of x-ray crystallography structure of AhlyH35A in complex with peptide 20

**Figure S3:** Peptide 20 electron density omit map

**Figure S4:** Binding epitope of Peptide 20 on Ahly

**Figure S5:** Similarities in Ahly binding of Peptide 88 and LTM14

**Figure S6:** Differing Ahly interactions of R07 in LTM14 and N10 in Peptide 20

**Figure S7:** Non-canonical amino acids tested in Peptide 20

**Table S2:** Effect of alanine substitution on Peptide 20 affinity to AviTag-Ahly as measured by SPR

**Table S3:** Effect of non-canonical substitution on Peptide 20 affinity to AviTag-Ahly as measured by SPR

**Table S4:** Effect of Peptide 20 truncation and N-terminal acetylation on affinity to AviTag-Ahly as measured by SPR

**Table S5:** Effect of combinations of multiple non-canonical amino acid substitutions on Peptide 20 binding affinity to AviTag-Ahly as measured by SPR

**Figure S8:** Flow cytometry gating strategy and median fluorescent intensity of controls

**Figure S9:** Peptide 88 is not toxic to A549 cells

**Figure S10:** Peptide 88 does not inhibit S. aureus growth

**Figure S11:** Supernatant Cleavage Assay LC/MS data

## Abbreviations

ADAM10: A disintegrin and metalloprotease 10;
Ahly: α-hemolysin;
AMR: antimicrobial resistance;
ANOVA: analysis of variance;
AviTag: Avi-tagged protein sequence;
caMHB: cation-adjusted Mueller Hinton broth;
DMSO: dimethyl sulfoxide;
DMEM: Dulbecco’s modified Eagle medium;
DTT: dithiothreitol;
EDTA: ethylenediaminetetraacetic acid;
FRET: Förster resonance energy transfer;
HEPES: 4-(2-hydroxyethyl)-1-piperazineethanesulfonic acid;
HPLC: high-performance liquid chromatography;
IC₅₀: half-maximal inhibitory concentration;
IPTG: isopropyl β-D-1-thiogalactopyranoside;
KD: equilibrium dissociation constant;
LB: lysogeny broth;
LC-MS: liquid chromatography–mass spectrometry;
MIC: minimum inhibitory concentration;
MRSA: methicillin-resistant Staphylococcus aureus;
OD600: optical density at 600 nm;
PBS: phosphate-buffered saline;
PDB: protein data bank;
PEG: polyethylene glycol;
PFT: pore-forming toxin;
RMSD: root mean square deviation;
SD: standard deviation;
SPR: surface plasmon resonance;
SSTI: skin and soft tissue infection;
TATB: 1,3,5-tris(bromoacetyl) hexahydro-1,3,5-triazine;
TB: terrific broth;
TCEP: tris(2-carboxyethyl)phosphine;
TFA: trifluoroacetic acid;
WT: wild type.

