## Supporting Information for "Discovery, characterisation and optimisation of bicyclic peptide inhibitors that disarm *Staphylococcus aureus* α-hemolysin"

**Contents**

**Figure S1:** C-terminal AviTag fusion does not perturb Ahly structure or function

**Figure S2:** Peptide 14 is a specific inhibitor of Ahly

**Table S1:** Statistics of x-ray crystallography structure of AhlyH35A in complex with peptide 20

**Figure S3:** Peptide 20 electron density omit map

**Figure S4:** Binding epitope of Peptide 20 on Ahly

**Figure S5:** Similarities in Ahly binding of Peptide 88 and LTM14

**Figure S6:** Differing Ahly interactions of R07 in LTM14 and N10 in Peptide 20

**Figure S7:** Non-canonical amino acids tested in Peptide 20

**Table S2:** Effect of alanine substitution on Peptide 20 affinity to Avitag-Ahly as measured by SPR

**Table S3:** Effect of non-canonical substitution on Peptide 20 affinity to AviTag-Ahly as measured by SPR

**Table S4:** Effect of Peptide 20 truncation and N-terminal acetylation on affinity to AviTag-Ahly as measured by SPR

**Table S5:** Effect of combinations of multiple non-canonical amino acid substitutions on Peptide 20 binding affinity to Avitag-Ahly as measured by SPR

**Figure S8:** Flow cytometry gating strategy and median fluorescent intensity of controls

**Figure S9:** Peptide 88 is not toxic to A549 cells

**Figure S10:** Peptide 88 does not inhibit S. aureus growth

**Figure S11:** Supernatant Cleavage Assay LC/MS data

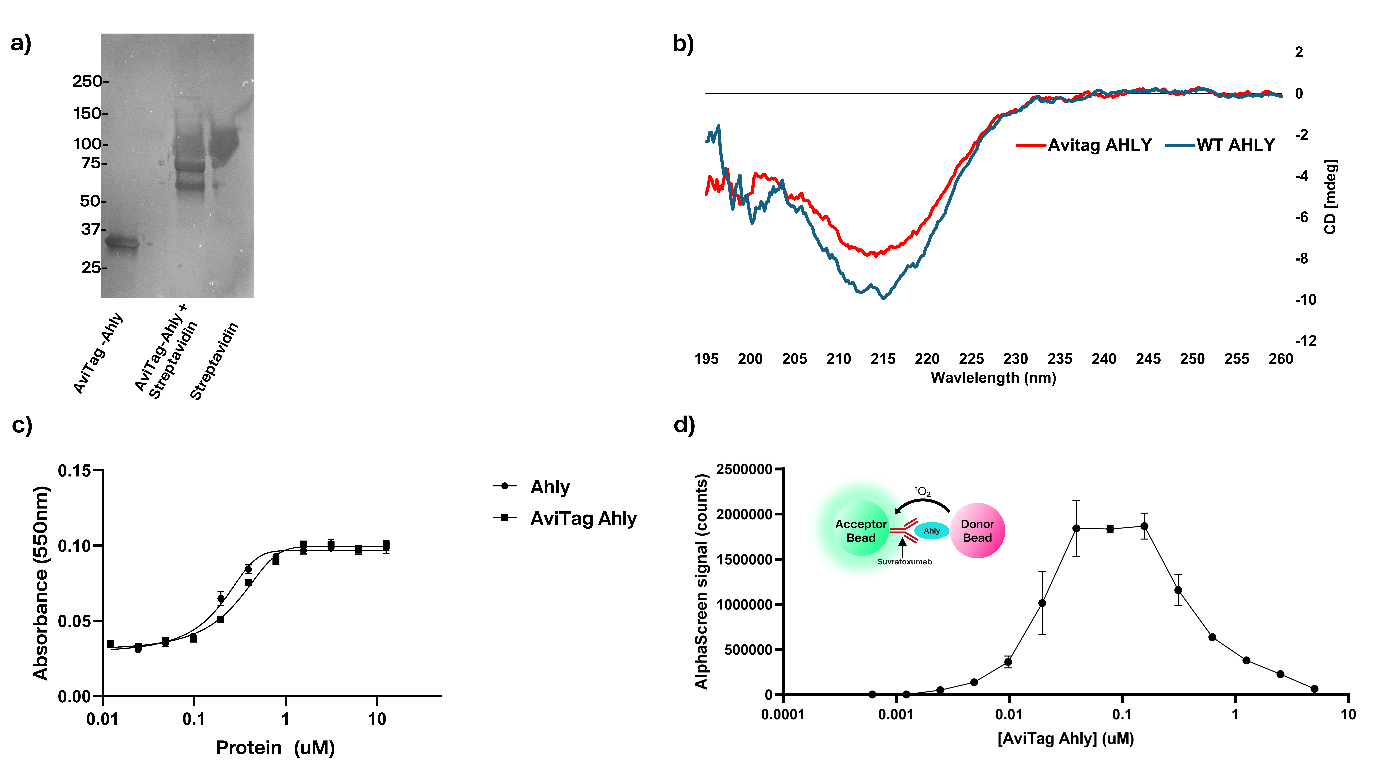

**Supplementary Figure 1: C-terminal AviTag fusion does not perturb Ahly structure or function**. **a)** The approximate percentage of biotinylation was estimated from the % reduction in free Avitag-Ahly band intensity, quantified in FIJI, upon pre-incubation with streptavidin in a gel shift assay. **b)** Circular dichroism traces of Ahly and Avitag-Ahly. **c)** WT Ahly and Avitag-Ahly lytic activity measured in a hemolysis assay. **d)** Suvratoxumab binding to AviTag-Ahly measured by AlphaScreen assay. AviTag-Ahly was titrated against a fixed concentration of Suvratoxumab (3nM), generating a characteristic hook effect. Mean of 3 technical replicates; error bars represent standard deviation (SD), au = arbitrary units.

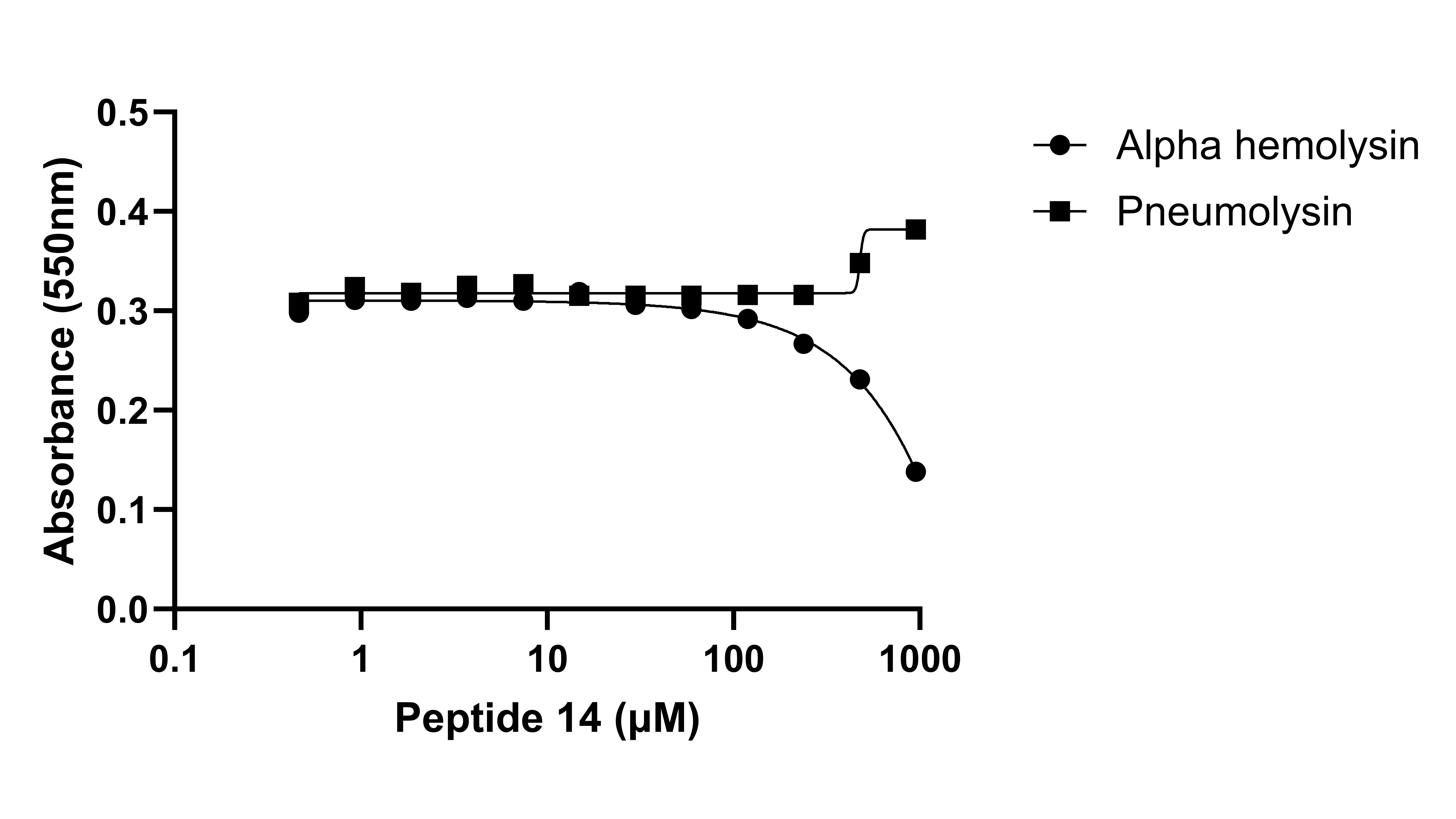

**Supplementary Figure 2: Peptide 14 is a specific inhibitor of Ahly**. Inhibition of Ahly (20 nM) and pneumolysin (Ply) (2nM) induced lysis of rabbit red blood cells by Peptide 14 in a hemolysis assay. Release of haemoglobin measured by absorbance at 550nm.

| **Table S1: Statistics of x-ray crystallography structure of AhlyH35A in complex with peptide 20**  **Structure** | **AhlyH35A:Peptide 20 (PDB: 9SVZ)** |
| --- | --- |
| **Space group** | P1 |
| **Wavelength (Å)** | 0.95373 |
| **Unit cell dimensions**  **a, b, c (Å) / α, β, γ (°)** | 53.8388, 59.0409, 72.4730, 71.1499, 86.3643, 70.5841 |
| **Resolution range^a^ (Å)** | 52.63 – 1.98 |
| **No. Reflections** | 47389 |
| **Unique reflections** | 32677 |
| **Completeness (%)** | 98.6 (98) |
| **R_merge_** | 0.085 (0.534) |
| **R_pim_** | 0.085 (0.534) |
| **CC_1/2_** | 0.995 (0.825) |
| **Redundancy** | 3.2 (3.5) |
| **< I/σ(I)>** | 7.9 (2.3) |
| **Wilson B (Å^2^)** | 32.177 |
| **R_work_/R_free_** | 0.1747 / 0.2137 |
| **DPI ^g^ (Å)** | 0.2797 |
| **Bond lengths (Å) / angles (º)** | 0.0116 / 1.8210 |
| **Average protein atoms B-factors (Å^2^)** | 37.17 |
| **Protein atoms (average B-factors, Å^2^)** | 4503 (37.17) |
| **Water molecules (average B-factors, Å^2^)** | 154 (35.268) |
| **Ligand (average B-factors, Å)** | 146 (36.747) |
| **Ramachandran analyses** |  |
| **Favoured regions (%)** | 94.2 |
| **Allowed regions (%)** | 99.5 |

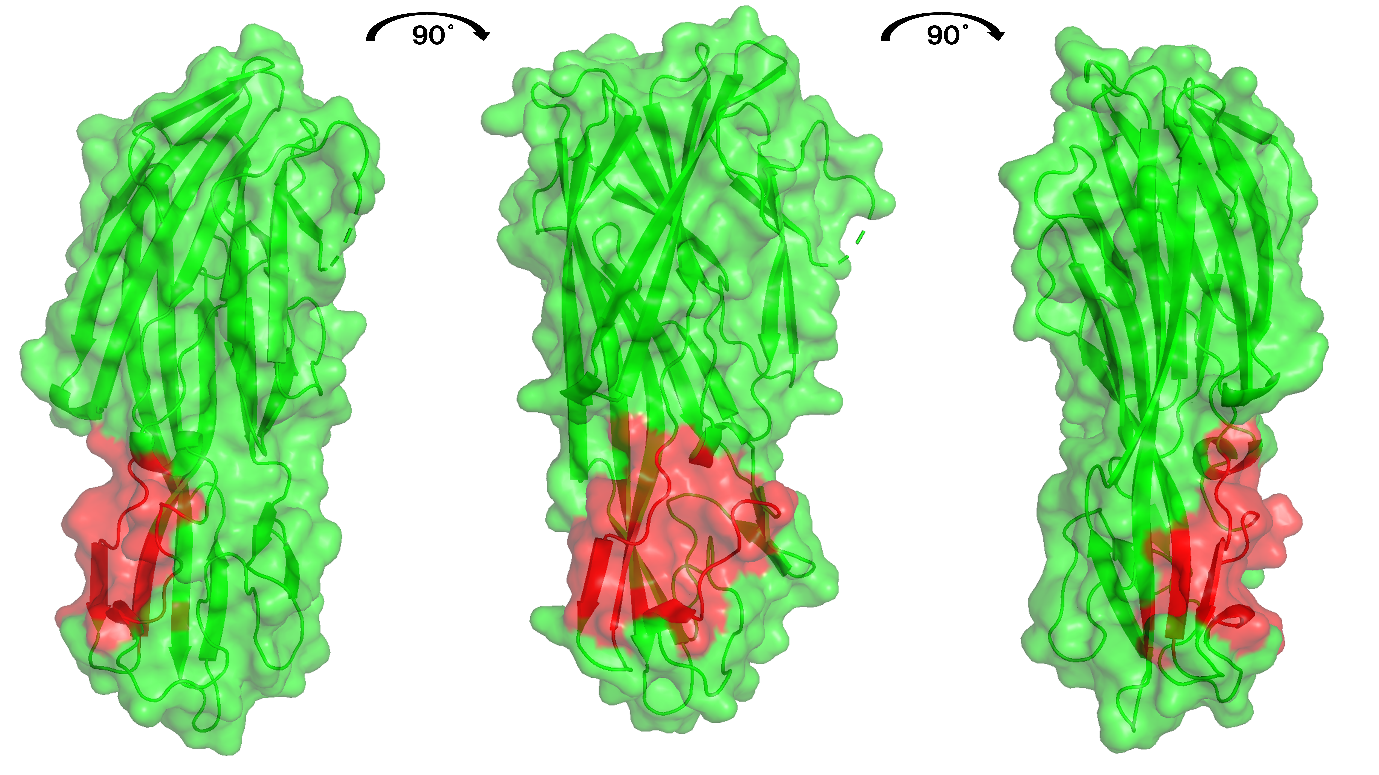

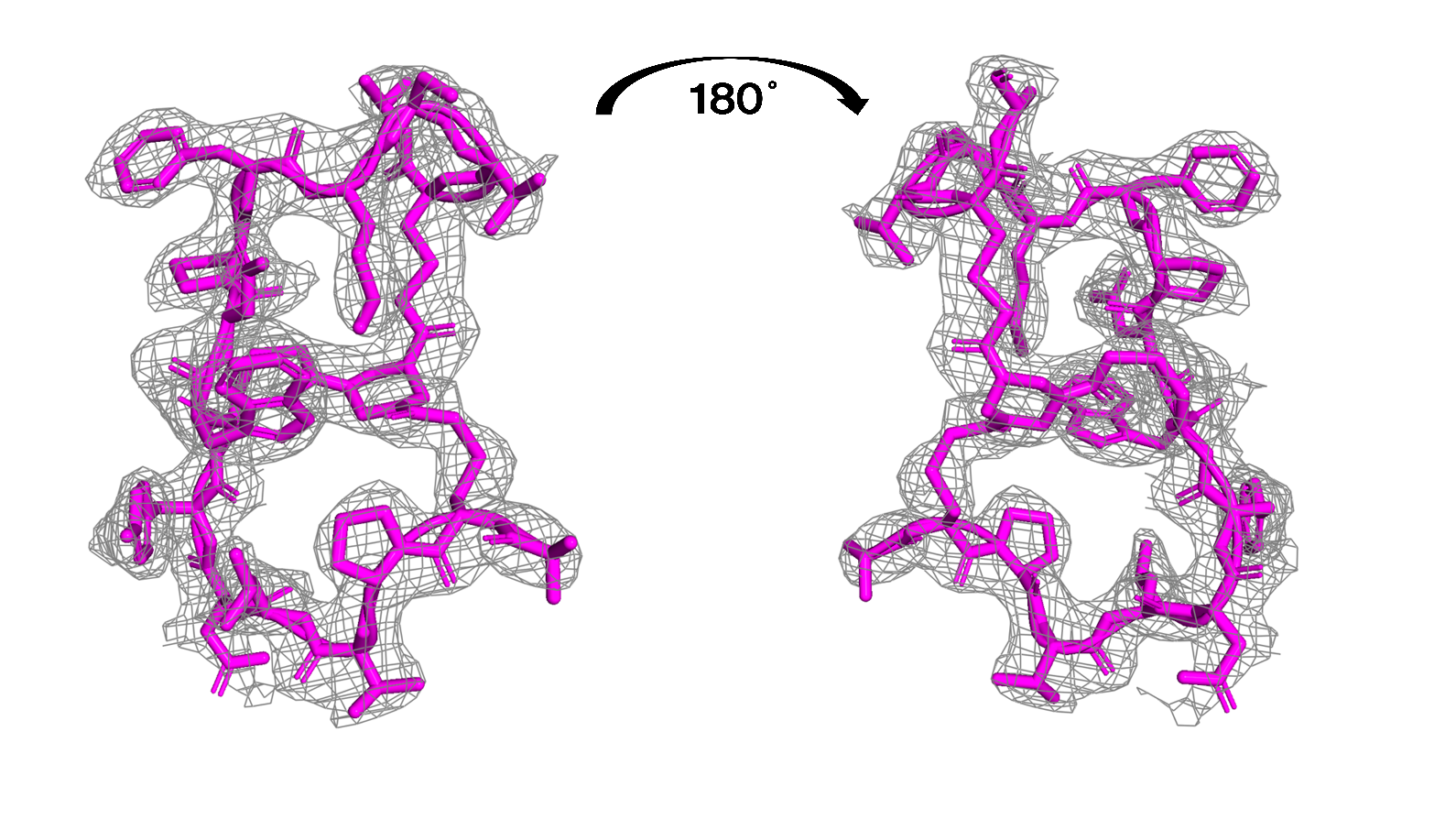

**Figure S4: Binding epitope of Peptide 20 on Ahly.** Peptide 20 binding site on AhlyH35A shown in red from AhlyH35A co-crystal structure.

**Figure S3: Peptide 20 electron density omit map.** Electron density omit map of Peptide 20 (grey) from co-crystal structure with Ahly with the structural model fitted to that density (pink). Electron density F_O_-F_C_ omit map for Peptide 20 was generated using REFMAC and visualised in PyMOL.

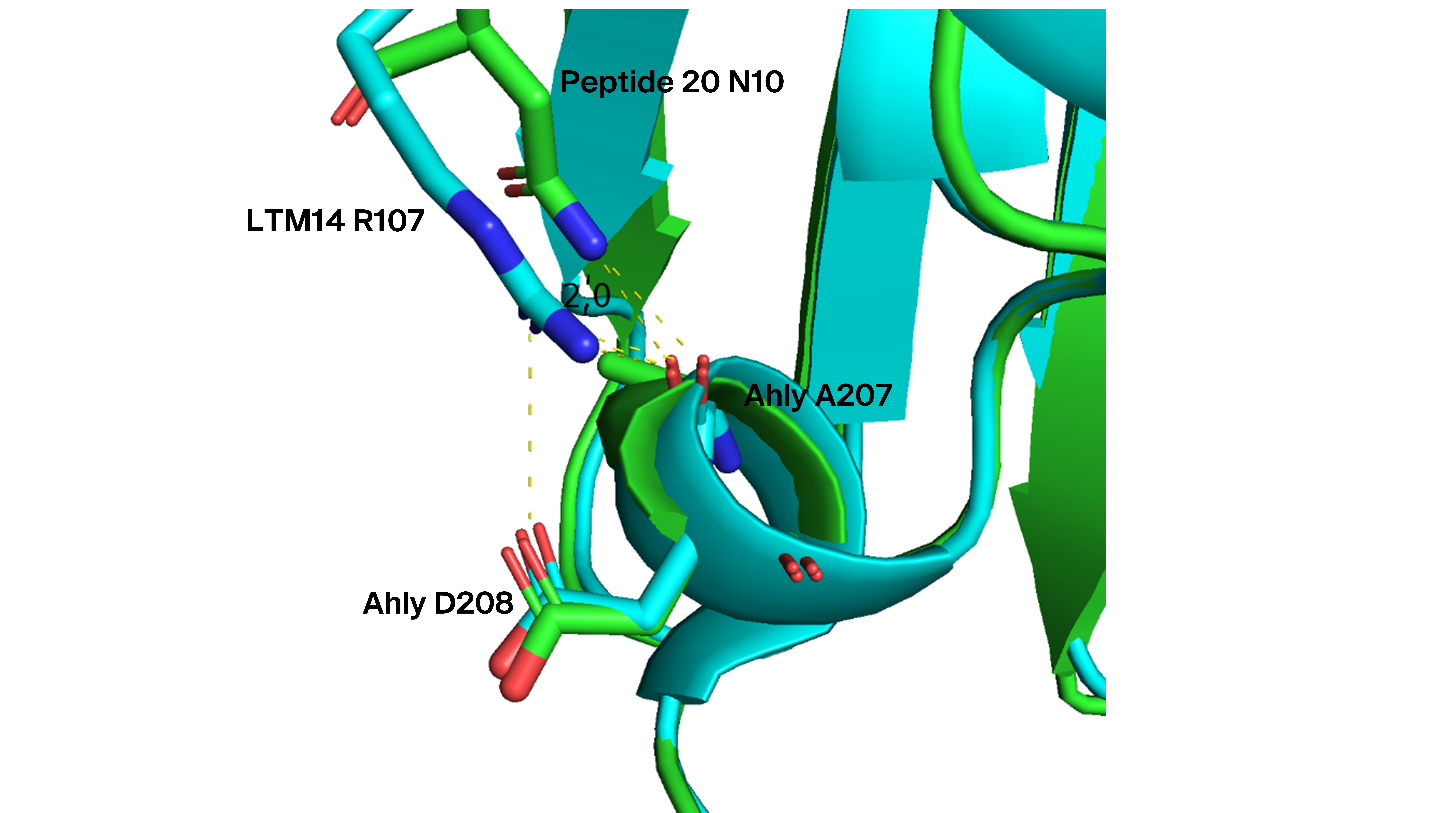

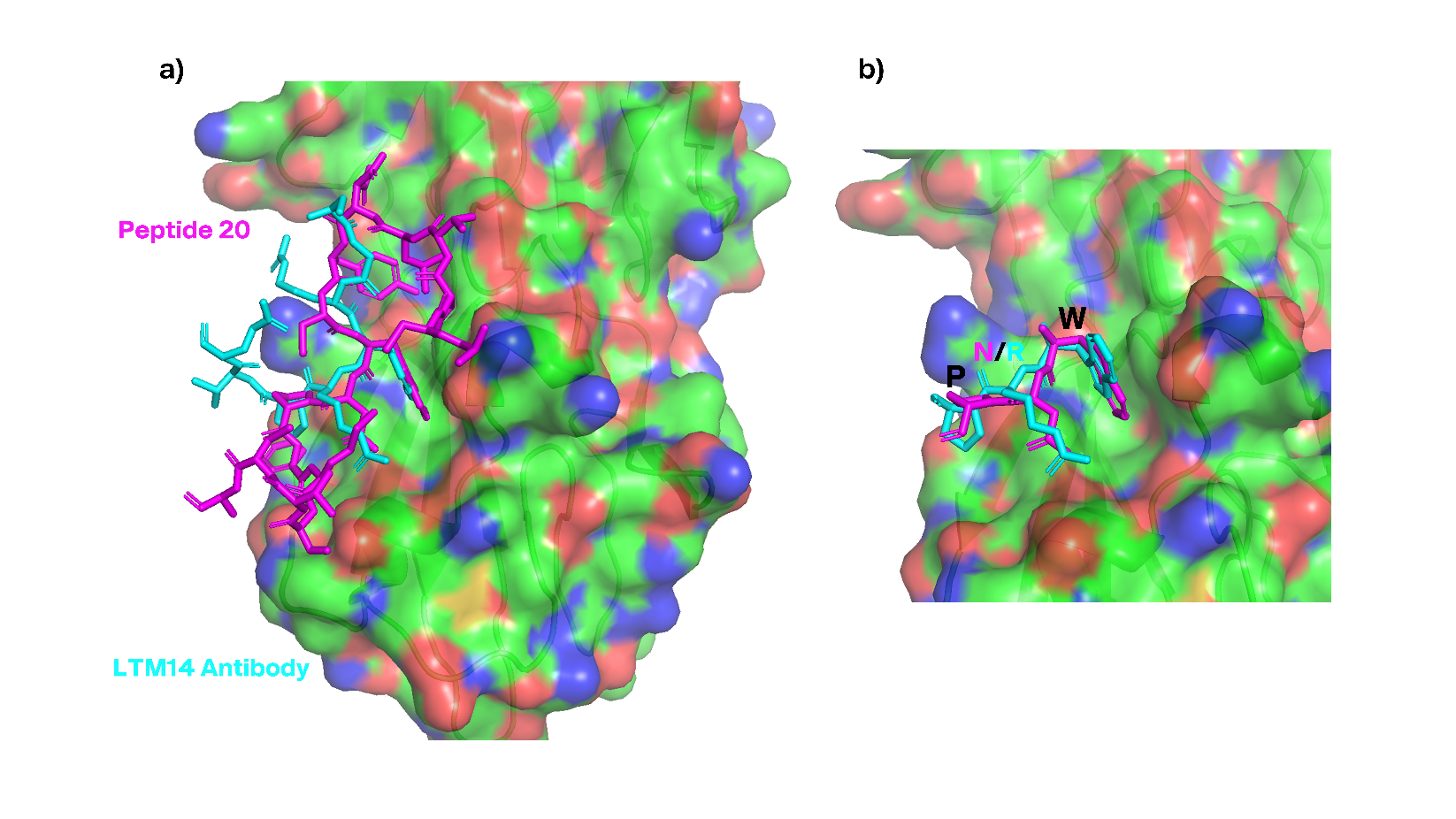

**Figure S5: Similarities in Ahly binding of Peptide 88 and LTM14.** **a)** Structure of Peptide 20 (magenta) bound to AhlyH35A, aligned with a portion of the antibody LTM14 from its co-crystal structure to AhlyH35L (Cyan) (PDB: 4IDJ). **b)** Alignment of ‘WNP’ motif of Peptide 20 with the corresponding ‘WRP’ motif of LTM14. Ahly surface positive charges shown in blue and negative charges shown in Red.

**Figure S6: Differing Ahly interactions of R07 in LTM14 and N10 in Peptide 20.** Structural alignment of LTM14 (blue) and Peptide 20 (green) co-crystal structures with Ahly, R107 of LTM14 forms a salt bridge with D208 of Ahly, while N10 of Peptide 20 forms a hydrogen bond with A207. Bonds are shown by dashed yellow lines.

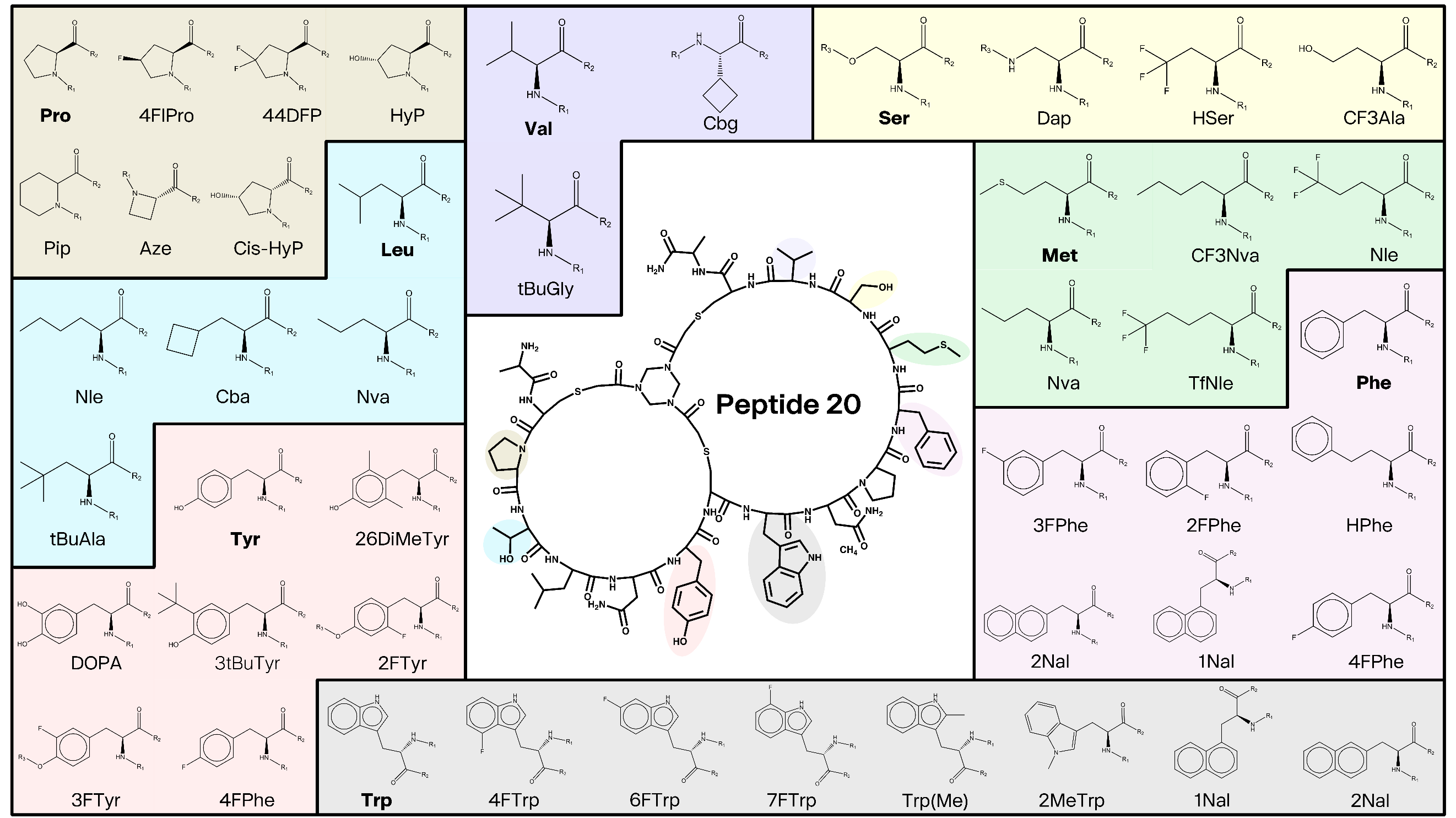

**Table S2: Effect of alanine substitution on Peptide 20 affinity to AviTag-Ahly as measured by SPR**

**Figure S7: Non-canonical amino acids tested in Peptide 20.** Chemical structure of Peptide 20, amino acids highlighted and the corresponding non canonical amino acids tested at these positions colour coded.

| **Bicyclic Peptide** | **Residue Replaced with Alanine** | **Geometric mean K_D_ (nM) (N=3)** | **SD** |
| --- | --- | --- | --- |
| Peptide 21 | P | 500 | 131 |
| Peptide 22 | T | 315 | 40 |
| Peptide 23 | L | 11367 | 4757 |
| Peptide 24 | N | 1005 | 24 |
| Peptide 25 | Y | 41887 | 26941 |
| Peptide 26 | W | NB |  |
| Peptide 27 | N | NB |  |
| Peptide 28 | P | 2150 | 335 |
| Peptide 29 | F | NB |  |
| Peptide 30 | M | 501 | 100 |
| Peptide 31 | S | 1158 | 104 |
| Peptide 32 | V | 235 | 55 |

**Table S3: Effect of non-canonical substitution on Peptide 20 affinity to AviTag-Ahly as measured by SPR**

| **Bicyclic Peptide** | **Residue in Peptide 20** | **Non-** **canonical/canonical amino acid susbstitution** | **Geometric mean K_D_ (nM) (N=3)** | **SD** |
| --- | --- | --- | --- | --- |
| 33 | Pro3 | HyP | 369 |  |
| 34 |  | 4FlPro | 479 | 25 |
| 35 |  | 44DFP | 185 | 45 |
| 36 |  | Cis-HyP | 618 | 88 |
| 37 |  | Pip | 346 | 64 |
| 38 |  | Aze | 693 | 145 |
| 39 | Leu5 | Nle | 1306 | 176 |
| 40 |  | Cba | 650 | 32 |
| 41 |  | Nva | 931 | 197 |
| 42 |  | tBuAla | 1138 | 417 |
| 43 | Asp6 | D | 402 |  |
| 44 | Tyr7 | 26DiMeTyr | 9923 | 3027 |
| 45 |  | DOPA | 689 | 62 |
| 46 |  | 3tBuTyr | 15622 | 4073 |
| 47 |  | 2FTyr | 785 | 87 |
| 48 |  | 3FTyr | 489 | 75 |
| 49 |  | 4FPhe | 6466 | 2277 |
| 50 | Trp9 | 1Nal | 1272 | 344 |
| 51 |  | 2Nal | 11706 | 2458 |
| 52 |  | Trp(Me) | 1326 | 197 |
| 53 |  | 2MeTrp | 1106 | 459 |
| 54 |  | 4FTrp | 727 | 306 |
| 55 |  | 6FTrp | 1694 | 417 |
| 56 |  | 7FTrp | 424 | 54 |
| 57 | Pro11 | 4FlPro | 1706 | 762 |
| 58 |  | HyP | 488 | 152 |
| 59 |  | Cis-HyP | 2397 | 480 |
| 60 |  | Pip | 1016 | 272 |
| 61 |  | Aze | 791 | 269 |
| 62 | Phe12 | HPhe | 83991 | 11172 |
| 63 |  | 1Nal | 869 | 244 |
| 64 |  | 2Nal | 532 | 39 |
| 65 |  | 2FPhe | 461 | 175 |
| 66 |  | 3FPhe | 426 | 15 |
| 67 |  | 4FPhe | 297 | 50 |
| 68 | Met13 | Nle | 115 | 14 |
| 69 |  | TfNle | 227 | 65 |
| 70 |  | CF3Nva | 304 | 90 |
| 71 |  | Nva | 456 | 131 |
| 72 | Ser14 | CF3Ala | 4914 | 2372 |
| 73 |  | Dap | 2679 | 896 |
| 74 |  | HSer | 2605 | 727 |
| 75 | Val15 | Cbg | 249 | 110 |
| 76 |  | C5g | 272 | 98 |
| 77 |  | tBuGly | 491 | 159 |
| 78 |  | M | 189 |  |
| 79 |  | I | 210 |  |
| 80 |  | L | 252 |  |
| 81 |  | F | 532 |  |

**Table S4: Effect of Peptide 20 truncation and N-terminal acetylation on affinity to AviTag-Ahly as measured by SPR**

| **Bicyclic Peptide** | **Sequence** | | | | | | | | | | | | | | | | | | | **K_D_ (nM)**  **(N=1)** |
| --- | --- | --- | --- | --- | --- | --- | --- | --- | --- | --- | --- | --- | --- | --- | --- | --- | --- | --- | --- | --- |
| 82 |  |  | C | P | T | L | N | Y | C | W | N | P | F | M | S | V | C | A | [CONH2] | 358 |
| 83 |  |  | C | P | T | L | N | Y | C | W | N | P | F | M | S | V | C | [CONH2] |  | 701 |
| 84 |  | [-Ac] | C | P | T | L | N | Y | C | W | N | P | F | M | S | V | C | A | [CONH2] | 435 |
| 85 |  | [-Ac] | C | P | T | L | N | Y | C | W | N | P | F | M | S | V | C | [CONH2] |  | 439 |
| 86 | [-Ac] | A | C | P | T | L | N | Y | C | W | N | P | F | M | S | V | C | A | [CONH2] | 389 |
| 87 | [-Ac] | A | C | P | T | L | N | Y | C | W | N | P | F | M | S | V | C | [CONH2] |  | 566 |

**Table S5: Effect of combinations of multiple non-canonical amino acid substitutions on Peptide 20 binding affinity to AviTag-Ahly as measured by SPR**

| **Bicyclic Peptide** | Sequence | | | | | | | | | | | | | | | | | | K_D_ (nM) (N=1) |
| --- | --- | --- | --- | --- | --- | --- | --- | --- | --- | --- | --- | --- | --- | --- | --- | --- | --- | --- | --- |
| Peptide 89 | [-Ac] | C | P | T | L | D | Y | C | W | N | P | F | [Nle] | S | M | C | A | [CONH2] | 118 |
| Peptide 90 | [-Ac] | C | P | A | L | D | Y | C | W | N | P | [4FPhe] | [Nle] | S | M | C | [CONH2] |  | 121 |
| Peptide 91 | A | C | P | T | L | N | Y | C | W | N | P | F | [Nle] | S | V | C | A | [CONH2] | 125 |
| Peptide 92 | [-Ac] | C | P | T | L | D | Y | C | W | N | P | F | [Nle] | S | A | C | A | [CONH2] | 133 |
| Peptide 93 | [-Ac] | C | [HyP] | A | L | D | Y | C | W | N | P | F | [Nle] | S | M | C | [CONH2] |  | 148 |
| Peptide 94 | [-Ac] | C | [HyP] | A | L | D | Y | C | W | N | P | [4FPhe] | [Nle] | S | M | C | [CONH2] |  | 152 |
| Peptide 95 | [-Ac] | C | P | A | L | D | Y | C | W | N | P | [4FPhe] | [Nle] | S | V | C | [CONH2] |  | 165 |
| Peptide 96 | [-Ac] | C | [HyP] | A | L | D | Y | C | W | N | P | [4FPhe] | [Nle] | S | M | C | [K(PYA)] | [CONH2] | 181 |
| Peptide 97 | [-Ac] | C | P | T | L | D | Y | C | W | N | P | F | [Nle] | S | [Cbg] | C | A | [CONH2] | 209 |
| Peptide 98 | [-Ac] | C | P | L | L | D | Y | C | W | N | P | F | [Nle] | S | V | C | [CONH2] |  | 228 |
| Peptide 99 | [-Ac] | C | [HyP] | A | L | D | Y | C | W | N | P | F | [Nle] | S | V | C | [CONH2] |  | 250 |
| Peptide 100 | [-Ac] | C | [HyP] | A | L | D | Y | C | W | N | [HyP] | [4FPhe] | [Nle] | S | M | C | [CONH2] |  | 253 |
| Peptide 101 | [-Ac] | C | P | M | L | D | Y | C | W | N | P | F | [Nle] | S | V | C | [CONH2] |  | 253 |
| Peptide 102 | [-Ac] | C | [HyP] | A | L | D | Y | C | [7FTrp] | N | P | [4FPhe] | [Nle] | S | M | C | [CONH2] |  | 270 |
| Peptide 103 | [-Ac] | C | P | A | L | D | Y | C | W | N | [HyP] | [4FPhe] | [Nle] | S | V | C | [CONH2] |  | 296 |
| Peptide 104 | [-Ac] | C | P | D | L | D | Y | C | W | N | P | F | [Nle] | S | V | C | [CONH2] |  | 326 |
| Peptide 105 | [-Ac] | C | P | F | L | D | Y | C | W | N | P | F | [Nle] | S | V | C | [CONH2] |  | 342 |
| Peptide 106 | [-Ac] | C | P | A | L | D | Y | C | W | N | [HyP] | F | [Nle] | S | V | C | [CONH2] |  | 420 |
| Peptide 107 | [-Ac] | C | [HyP] | A | L | D | Y | C | W | N | P | [4FPhe] | [Nle] | S | V | C | [CONH2] |  | 483 |
| Peptide 108 | [-Ac] | C | [HyP] | P | L | D | Y | C | W | N | P | [4FPhe] | [Nle] | S | M | C | [CONH2] |  | 522 |
| Peptide 109 | [-Ac] | C | [HyP] | A | L | D | Y | C | [7FTrp] | N | [HyP] | [4FPhe] | [Nle] | S | M | C | [CONH2] |  | 570 |
| Peptide 110 | [-Ac] | C | G | A | L | D | Y | C | W | N | P | [4FPhe] | [Nle] | S | M | C | [CONH2] |  | 866 |
| Peptide 111 | [-Ac] | C | [Dap] | A | L | D | Y | C | W | N | P | [4FPhe] | [Nle] | S | M | C | [CONH2] |  | 989 |
| Peptide 112 | [-Ac] | C | [Dab] | A | L | D | Y | C | W | N | P | [4FPhe] | [Nle] | S | M | C | [CONH2] |  | 1084 |
| Peptide 113 | [-Ac] | C | P | T | L | D | Y | C | W | N | P | F | [Nle] | S | [C5g] | C | A | [CONH2] | 2248 |
| Peptide 114 | [-Ac] | C | [HyP] | A | L | D | Y | C | W | N | P | [4FPhe] | [Nle] | [Dab] | M | C | [CONH2] |  | 4093 |
| Peptide 115 | [-Ac] | C | P | T | L | D | Y | C | W | N | P | F | [Nle] | S | L | C | A | [CONH2] | 9434 |
| Peptide 116 | [-Ac] | C | M | A | L | D | Y | C | W | N | P | F | [Nle] | S | V | C | [CONH2] |  | 45310 |

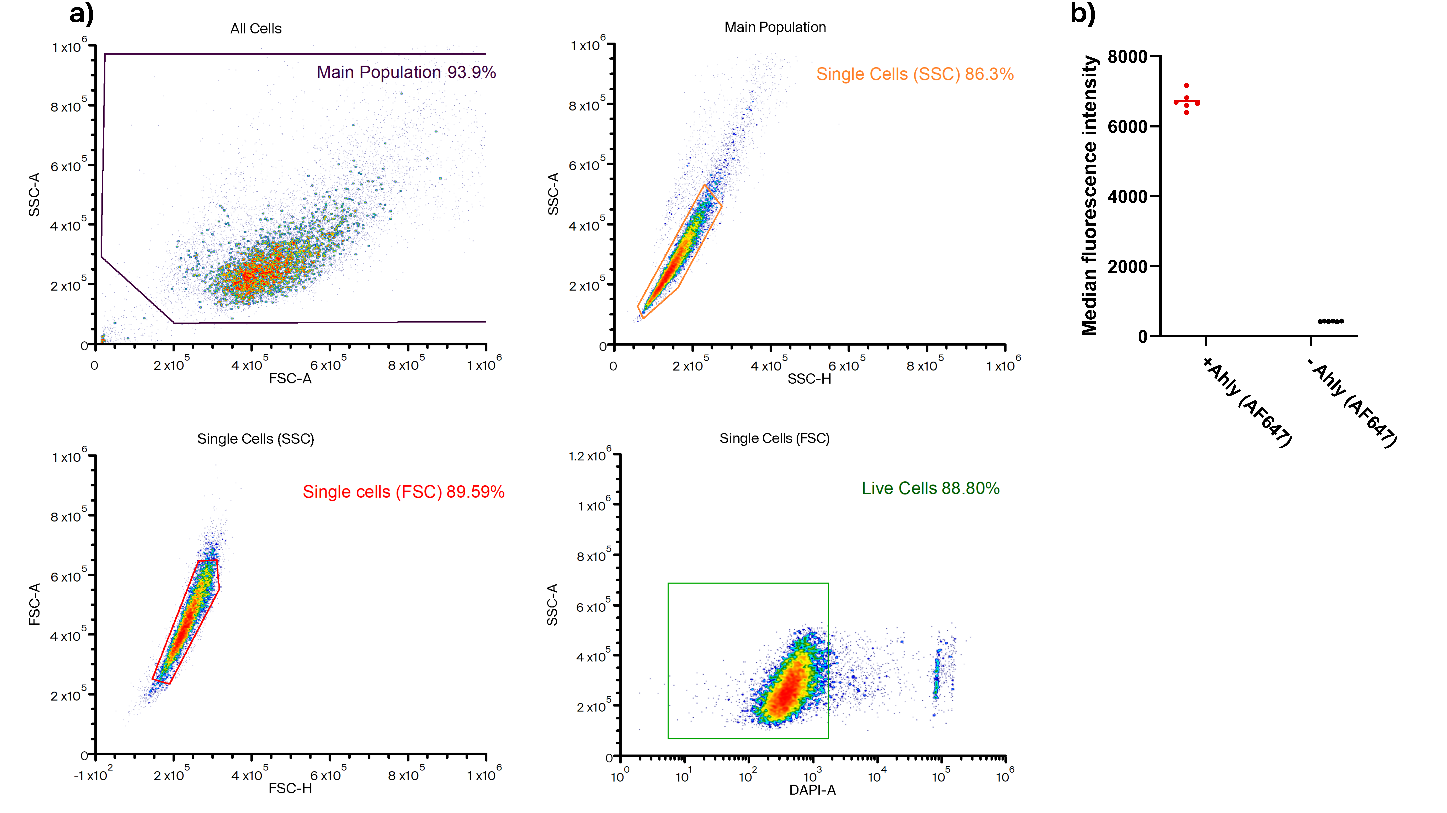

**Figure S8: Flow cytometry gating strategy and median fluorescent intensity of controls. a)** Flow cytometry gating strategy for bicyclic peptide inhibition of Alexa Flour 647 labelled AhlyH35A binding to A549 cells. Cells were sequentially gated as follows: (Top left) main cell population selected based on forward scatter area (FSC-A) vs side scatter area (SSC-A) to exclude cells debris. Followed by doublet exclusion by comparing side scatter area vs height (SSC-A vs SSC-H) and then by comparing forward scatter area vs height (FSC-A vs FSC-H) (top right and bottom left, respectively). Finally, cells were gated for live cells by exclusion of DAPI positive (dead) cells in the SSC-A vs DAPI-A plot (bottom right). **b)** Average median fluorescent intensity for A549 cells with or without incubation with of Alexa Fluor 647 labelled AhlyH35A, average of 6 technical repeats.

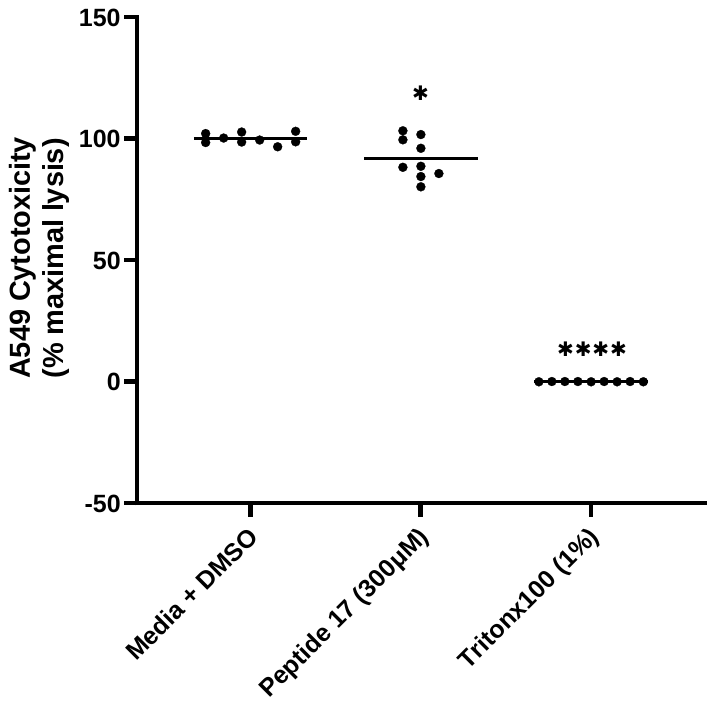

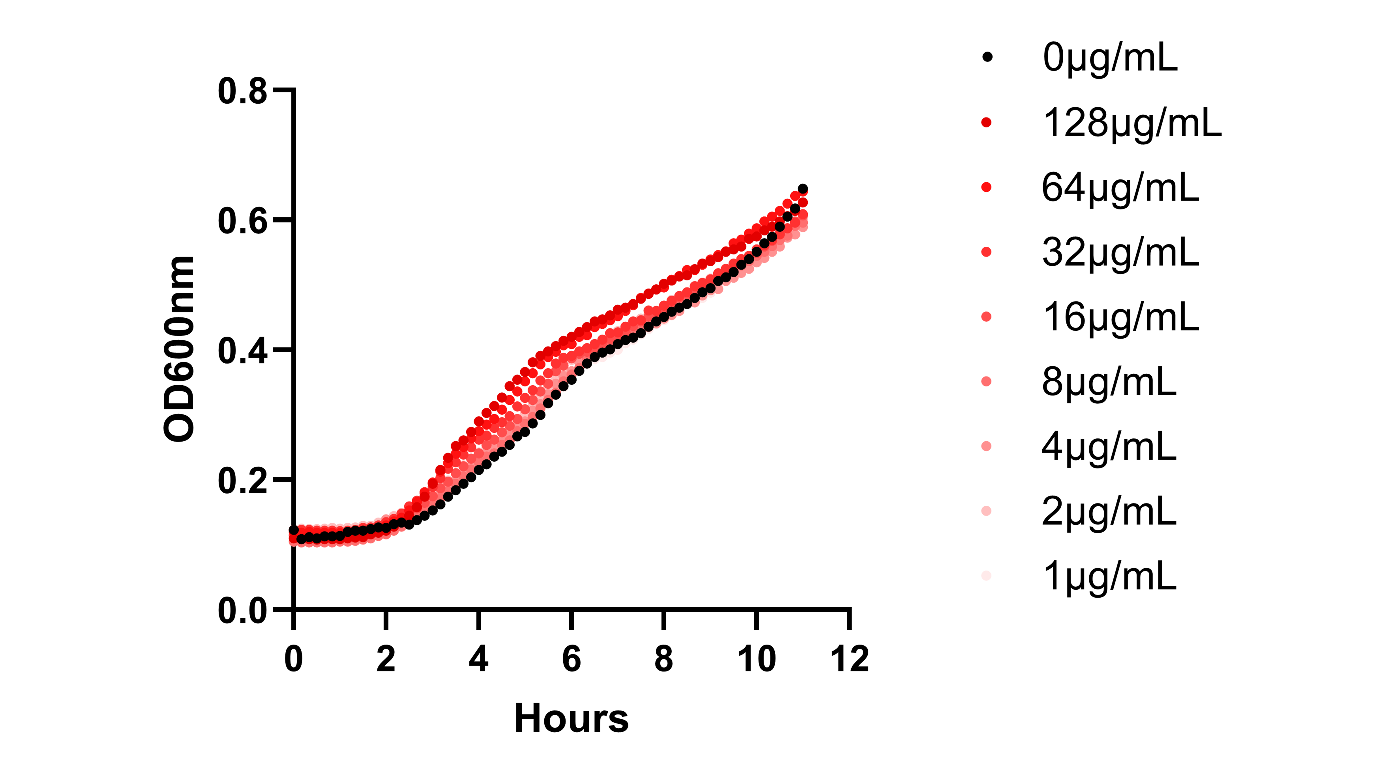

**Figure S9: Peptide 88 shows negligible toxicity to A549 cells.** A549 cell cytotoxicity measured after 72h incubation with vehicle (DMSO), Peptide 88 or 1% (v/v) Triton x100, DMSO concentrations were kept constant across all samples. Each point represents a technical replicate from 3 biological repeats with average shown. Data were analysed using one-way ANOVA with Dunnett’s test; * = P < 0.05; **** = P < 0.0001.

**Figure S10: Peptide 88 does not inhibit *S. aureus* growth.** Growth curve of *S. aureus* strain 8325-4 in the presence of increasing concentrations of Peptide 88.

**S11 Supernatant Cleavage Assay LC/MS data**

Media only :

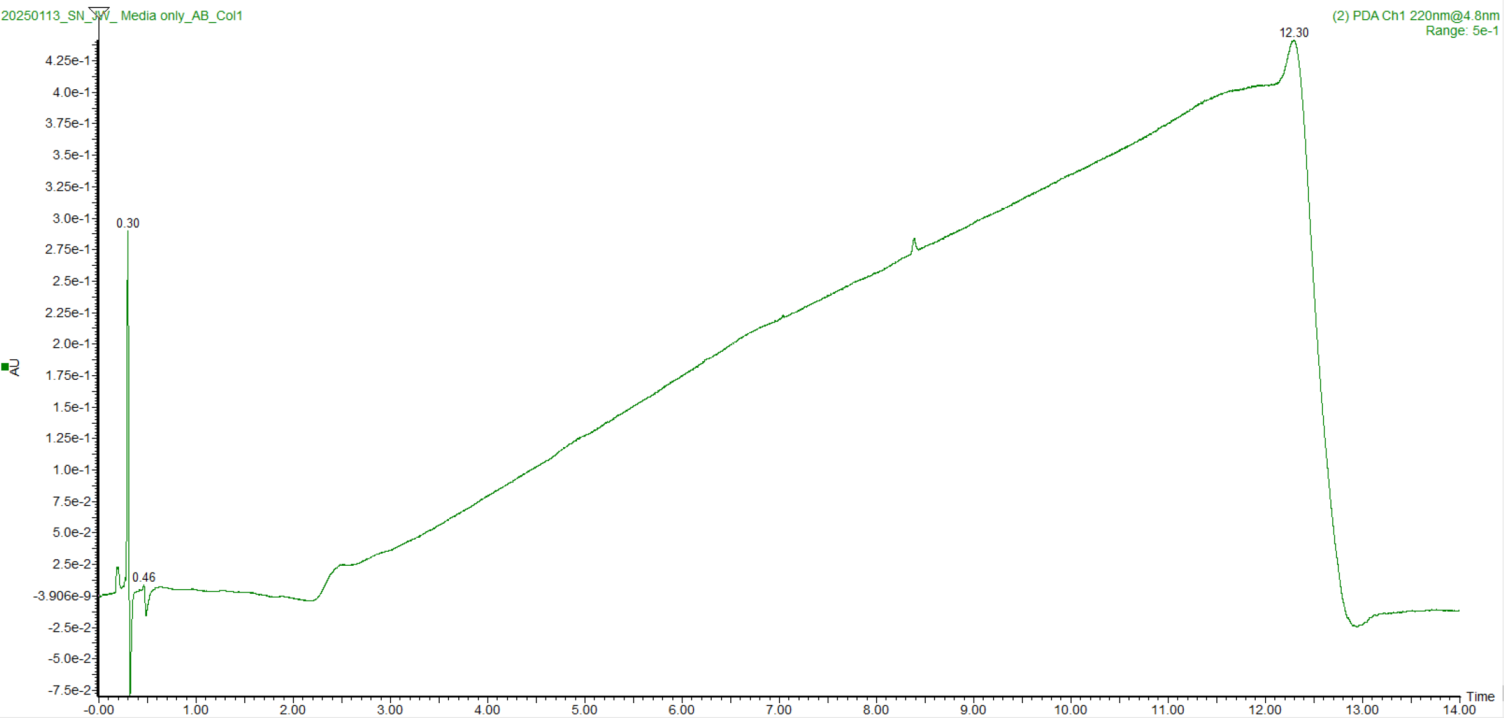

*S. aureus* supernatant only

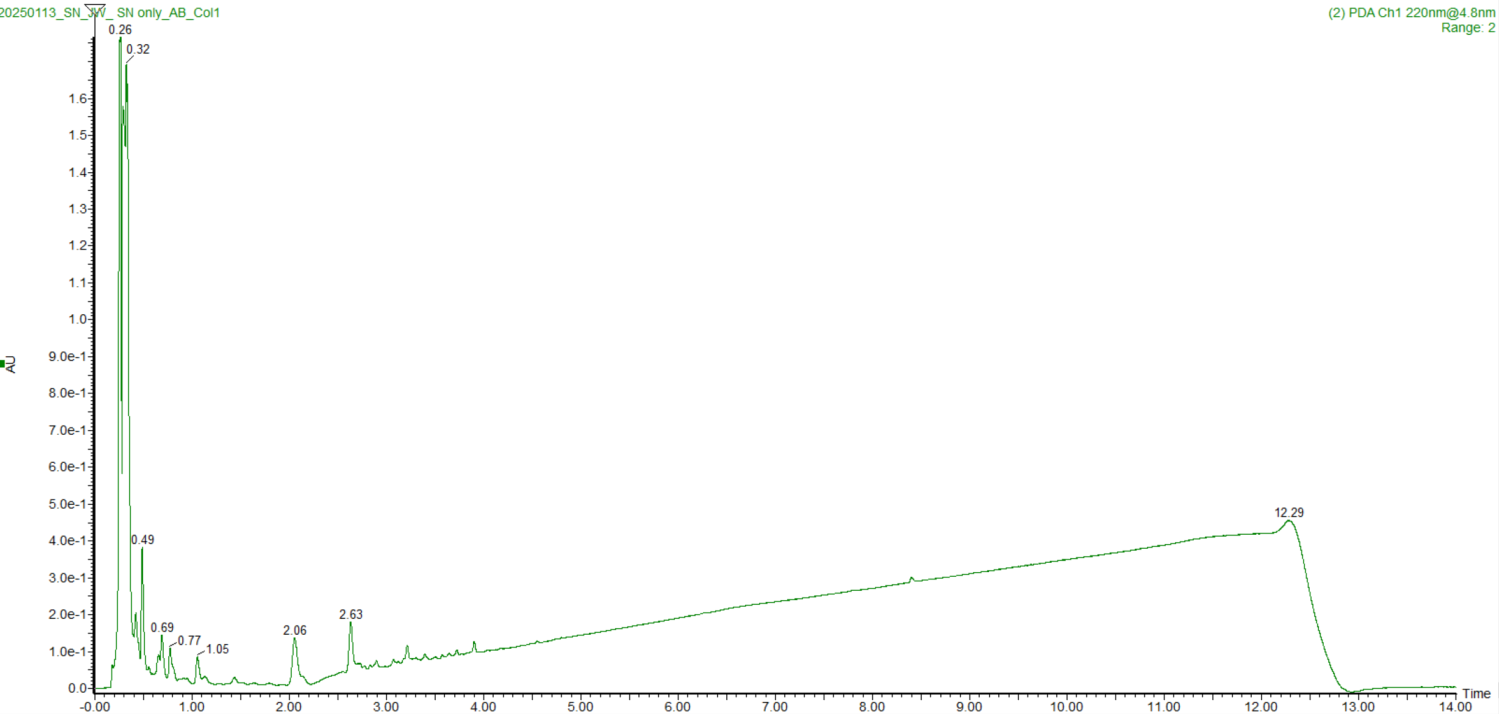

*S. aureus* supernatant + protease inhibitors

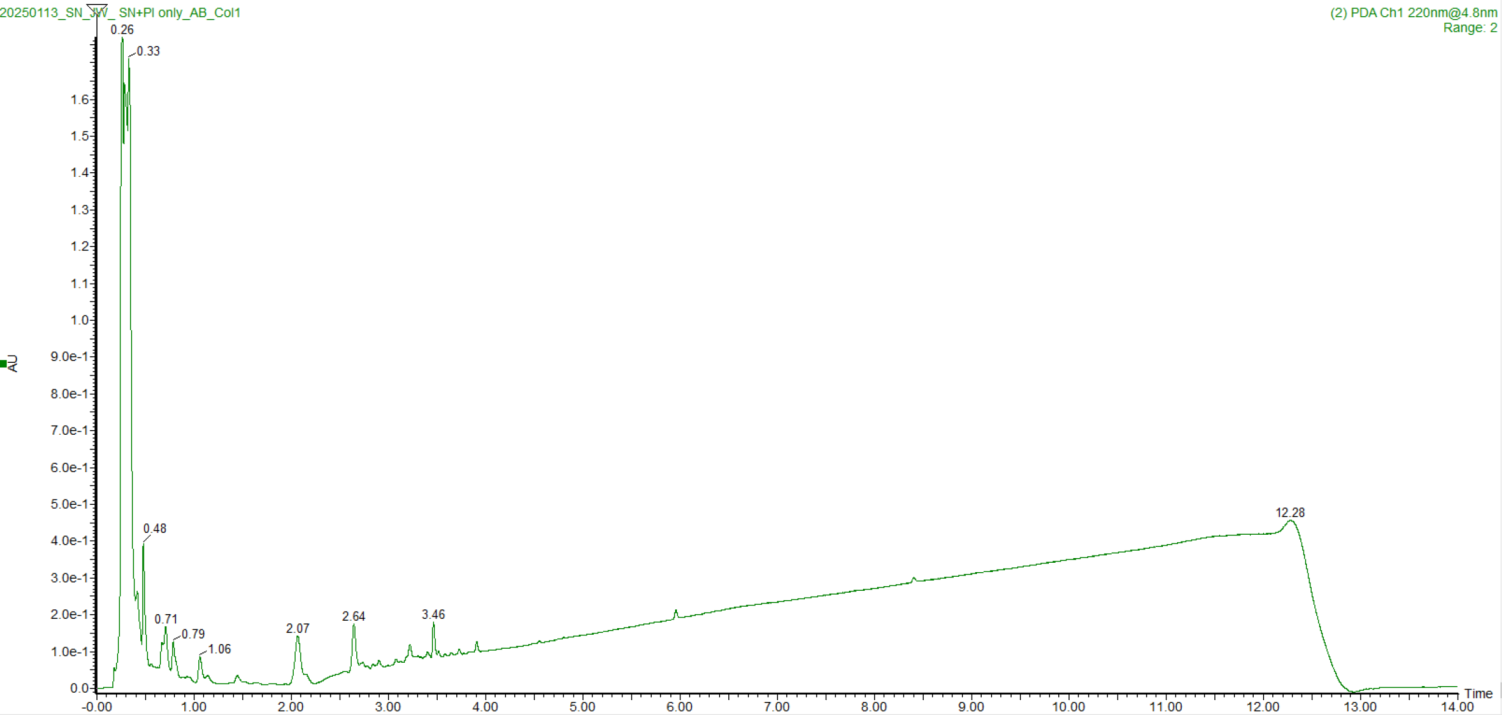

**Peptide 88 (MW: 2027.3)**

Media only

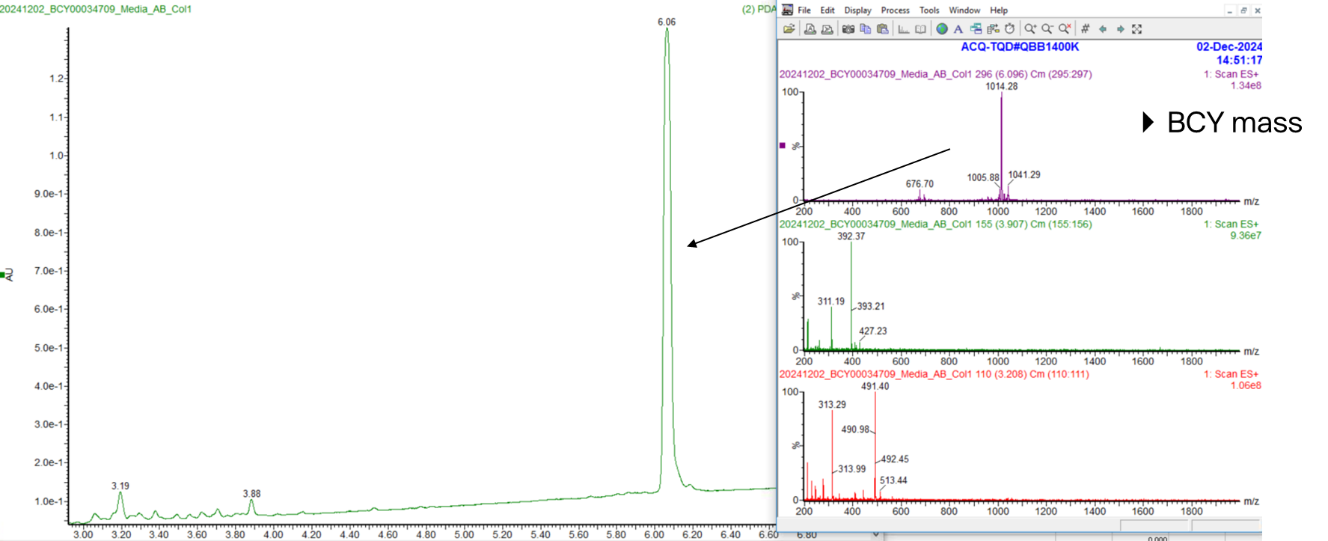

Peptide 88 + *S. aureus* supernatant + protease inhibitors

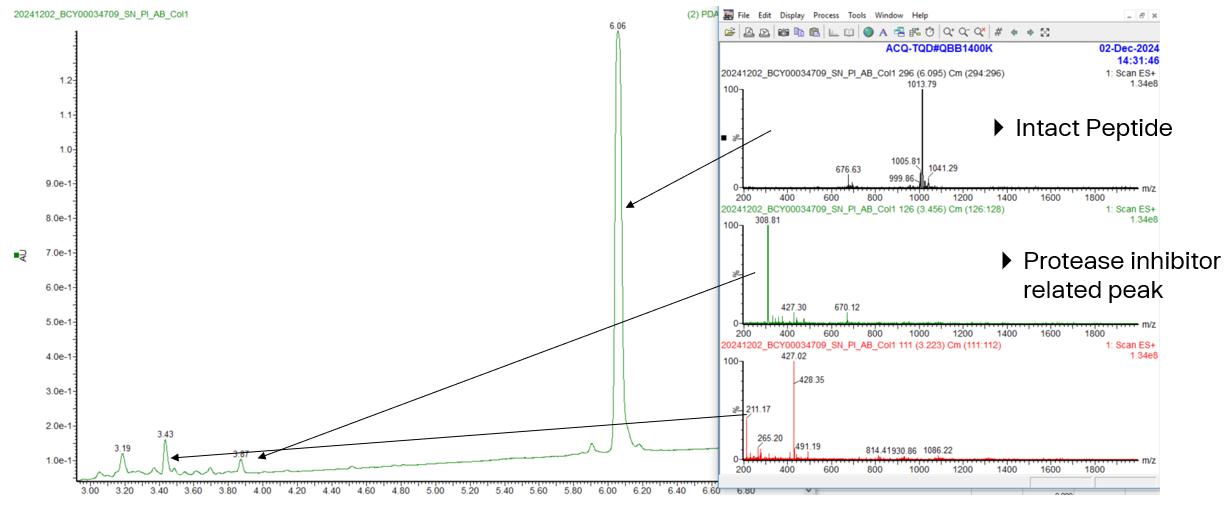

Peptide 88 + *S. aureus* supernatant

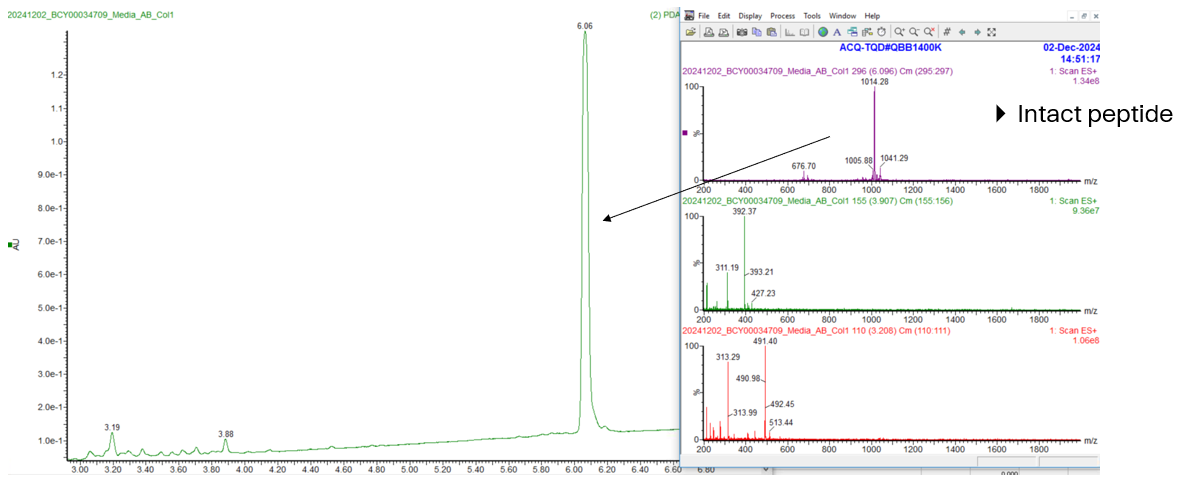

**Peptide 20 (MW: 2126.5)**

Peptide 20 + *S. aureus* supernatant

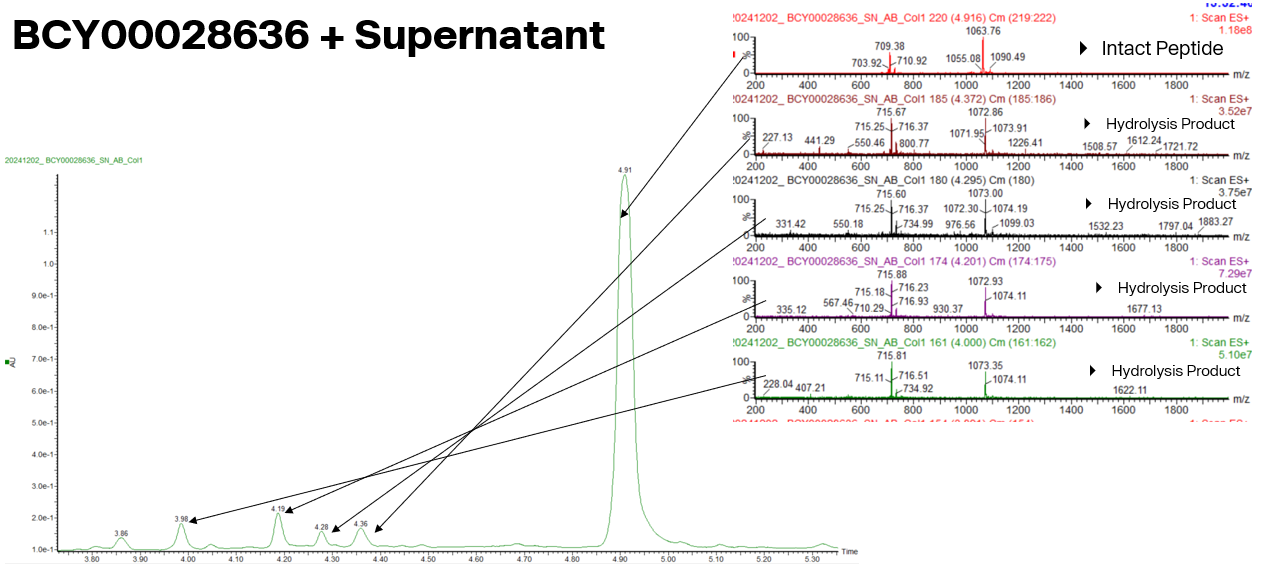

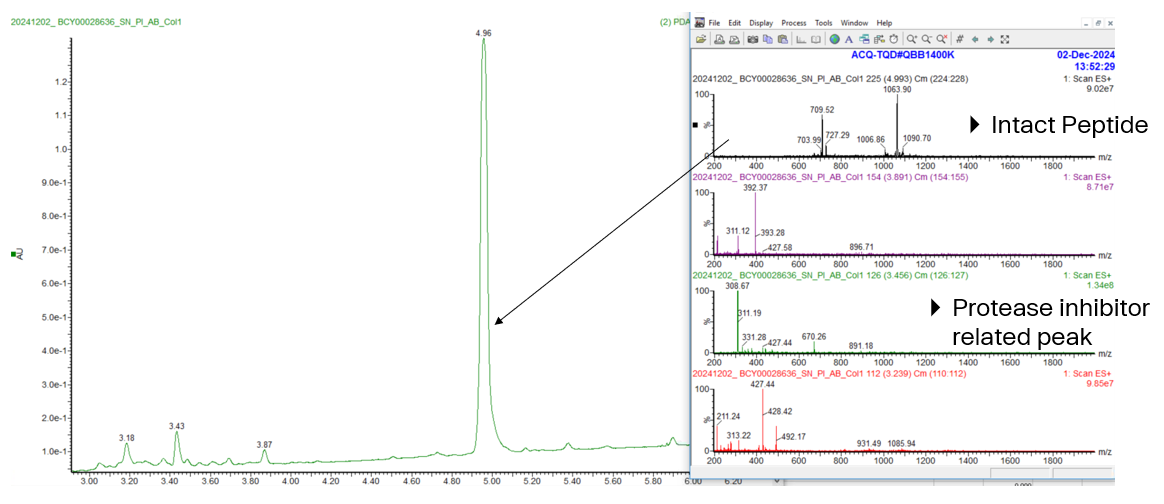
Peptide 20 + *S. aureus* supernatant + protease inhibitors

**Peptide 21 (MW: 2100.4)**

Peptide 21 + *S. aureus* supernatant

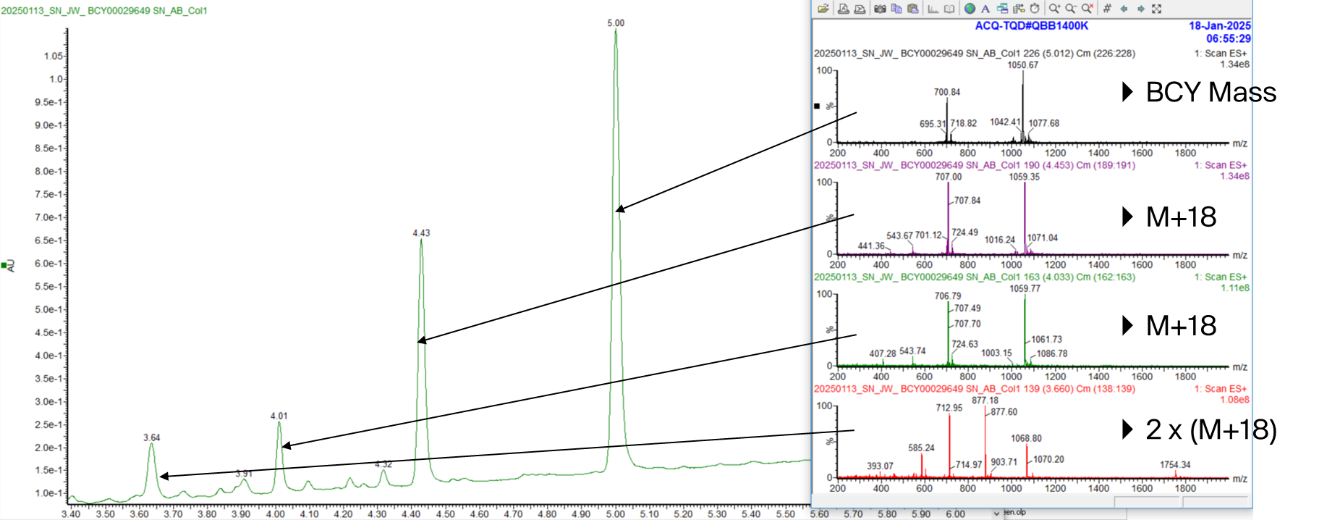
Peptide 21 + *S. aureus* supernatant + protease inhibitors

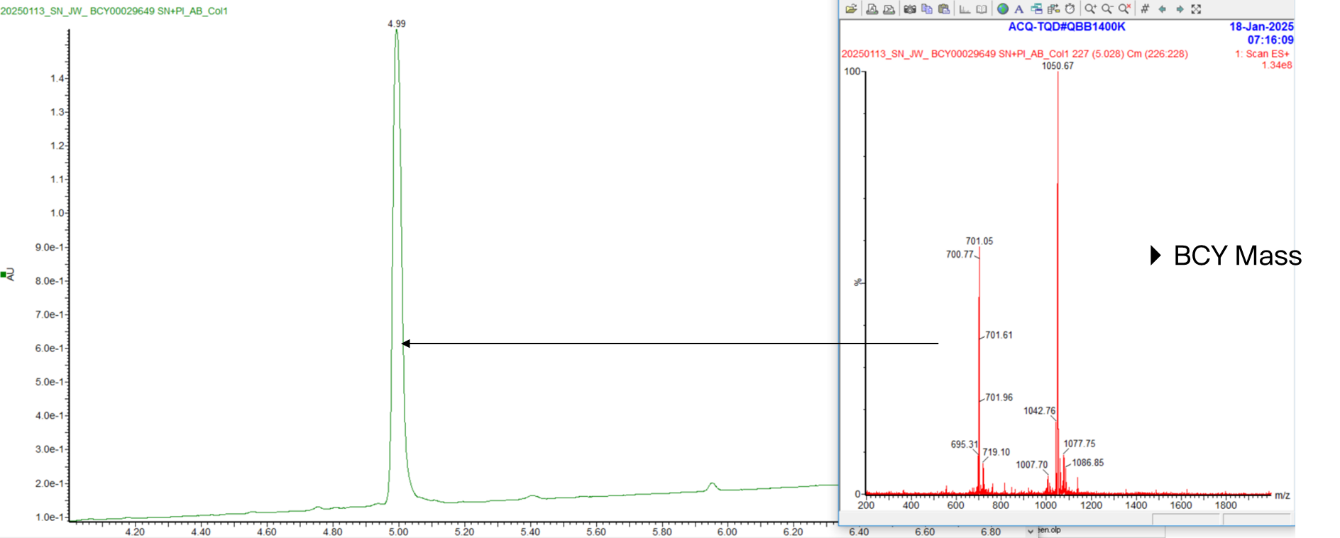

**Peptide 22 (MW: 2096.4)**

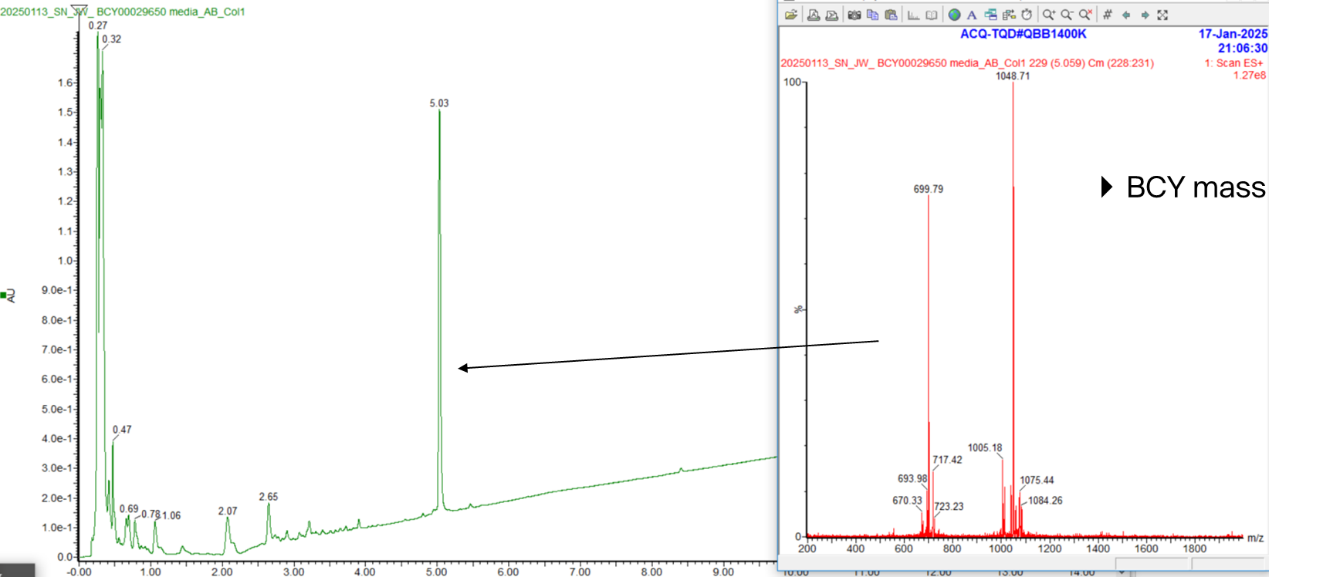
Peptide 22 + media

- BCY mass

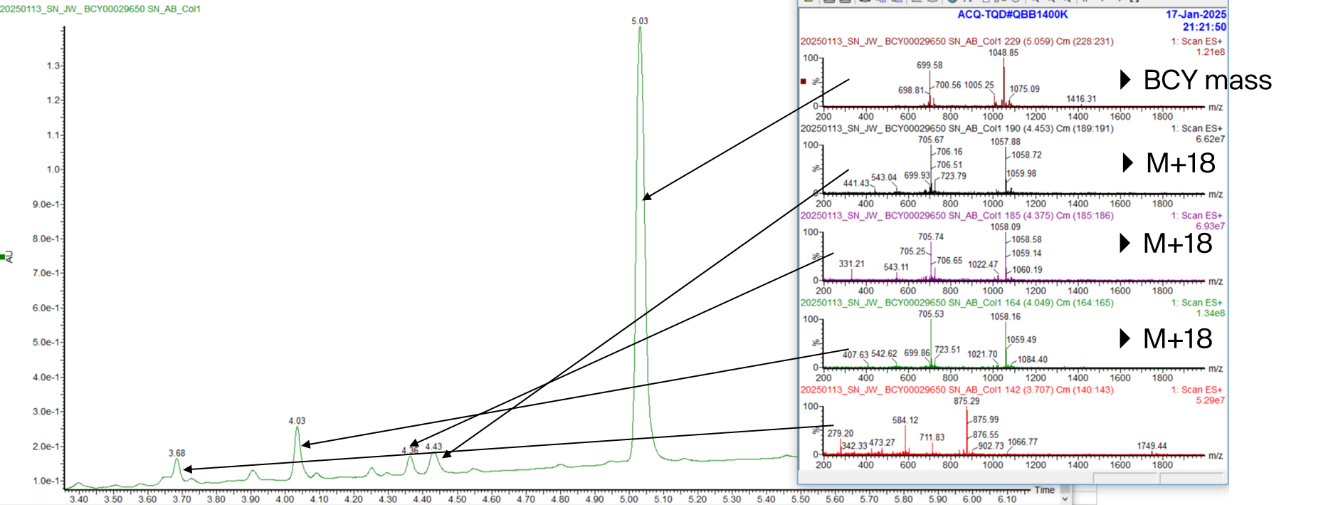
Peptide 22 + *S. aureus* supernatant

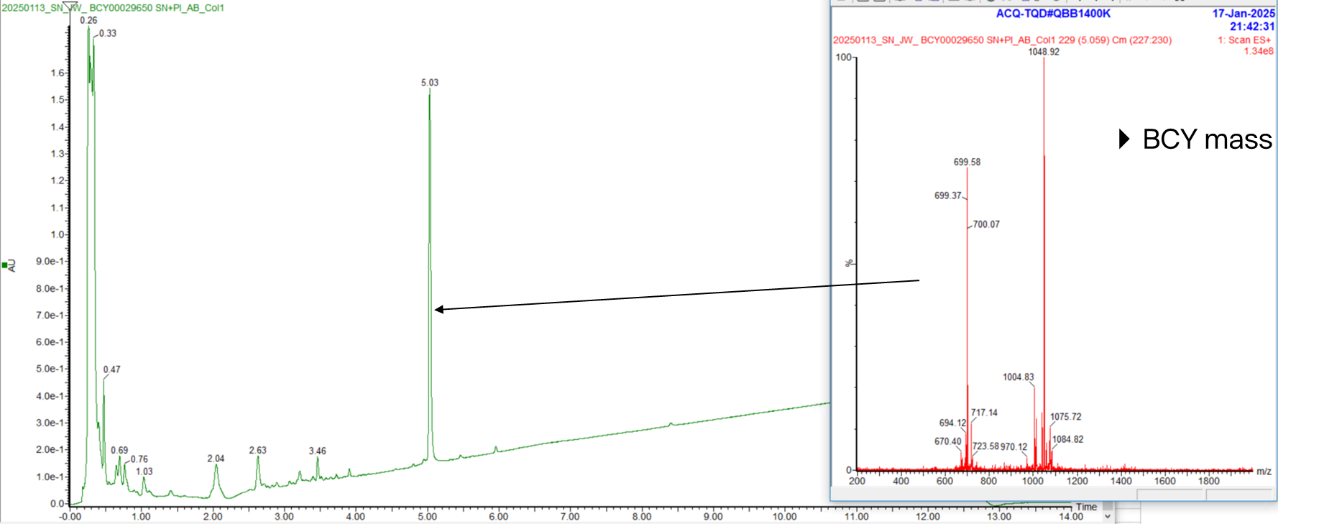
Peptide 22 + *S. aureus* supernatant + protease inhibitors

- BCY mass

**Peptide 23 (MW: 2084.4)**

Peptide 23 + media

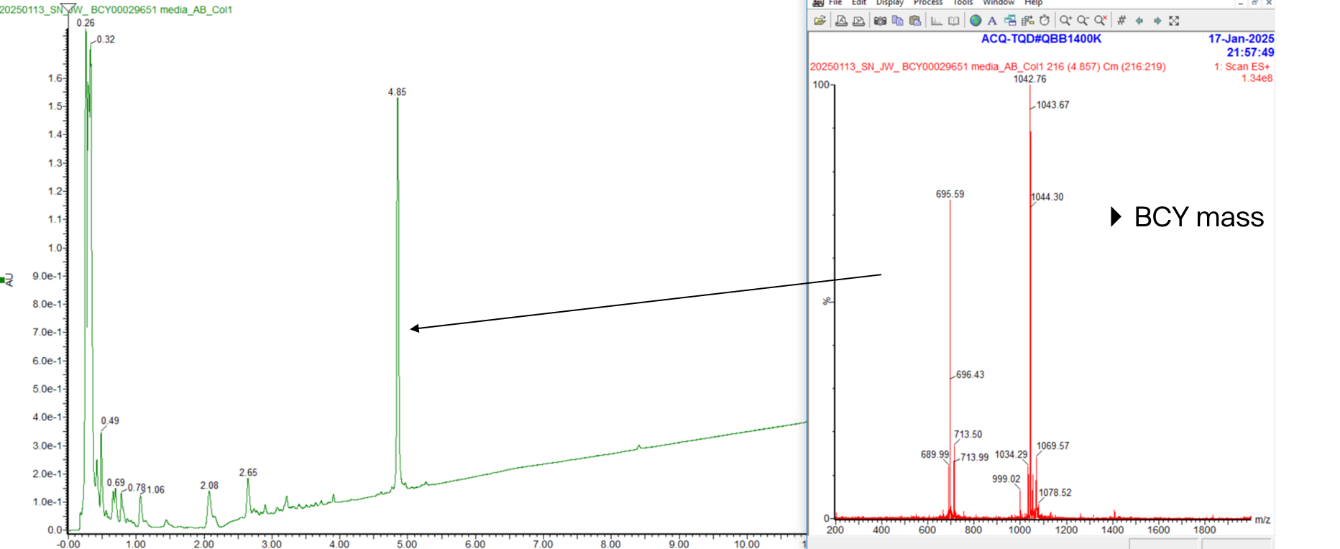

Peptide 23 + *S. aureus* supernatant

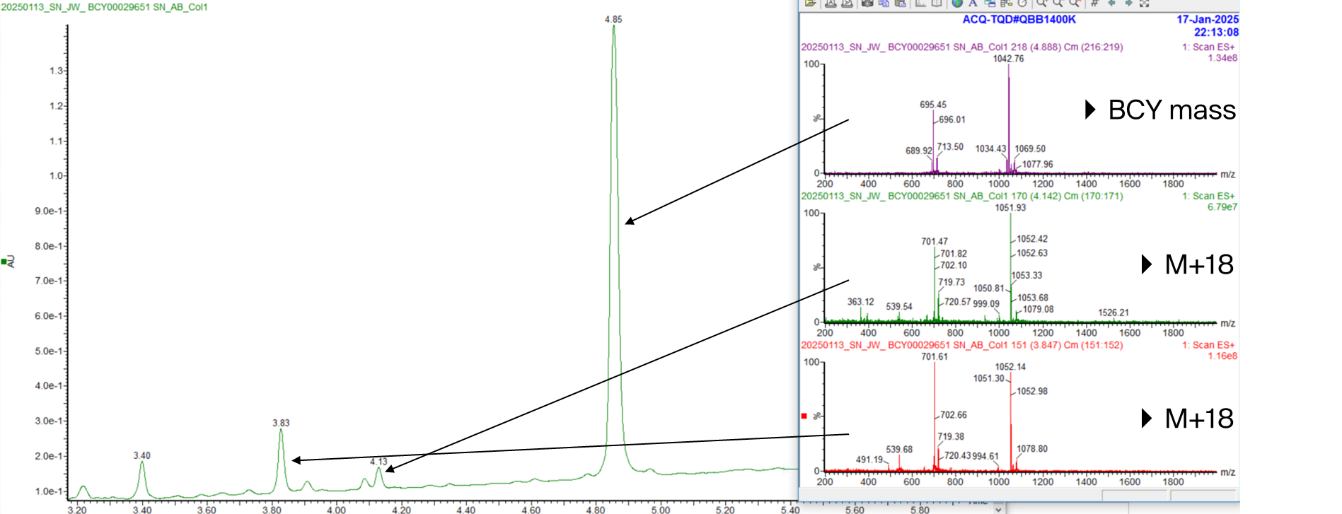

Peptide 23 + *S. aureus* supernatant + protease inhibitors

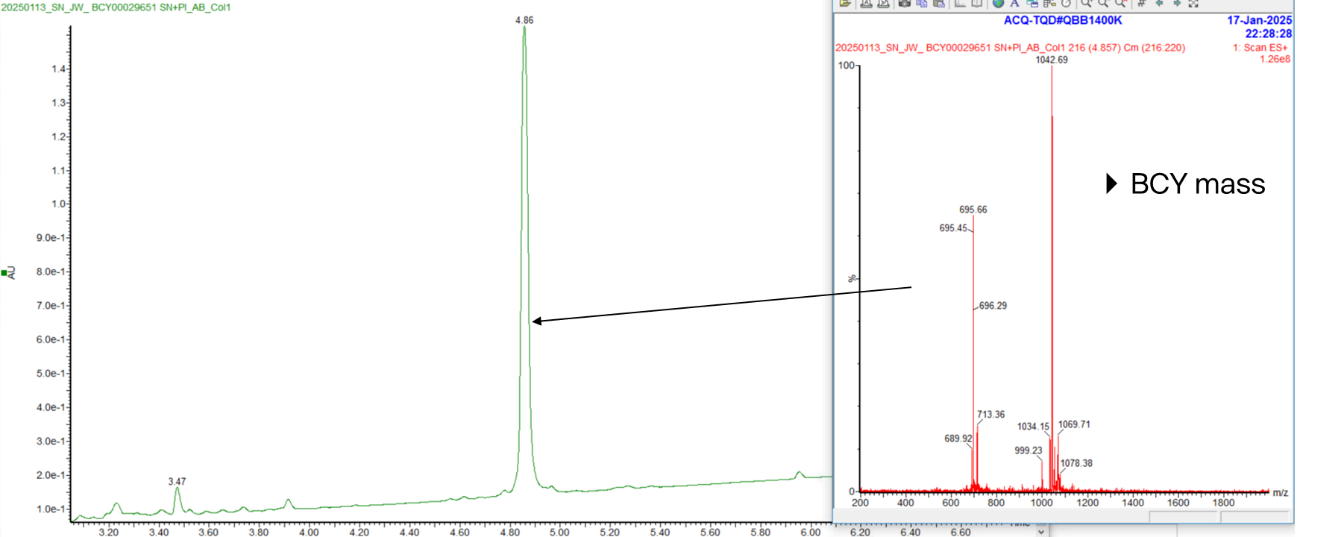

**Peptide 24 (MW: 2083.4)**

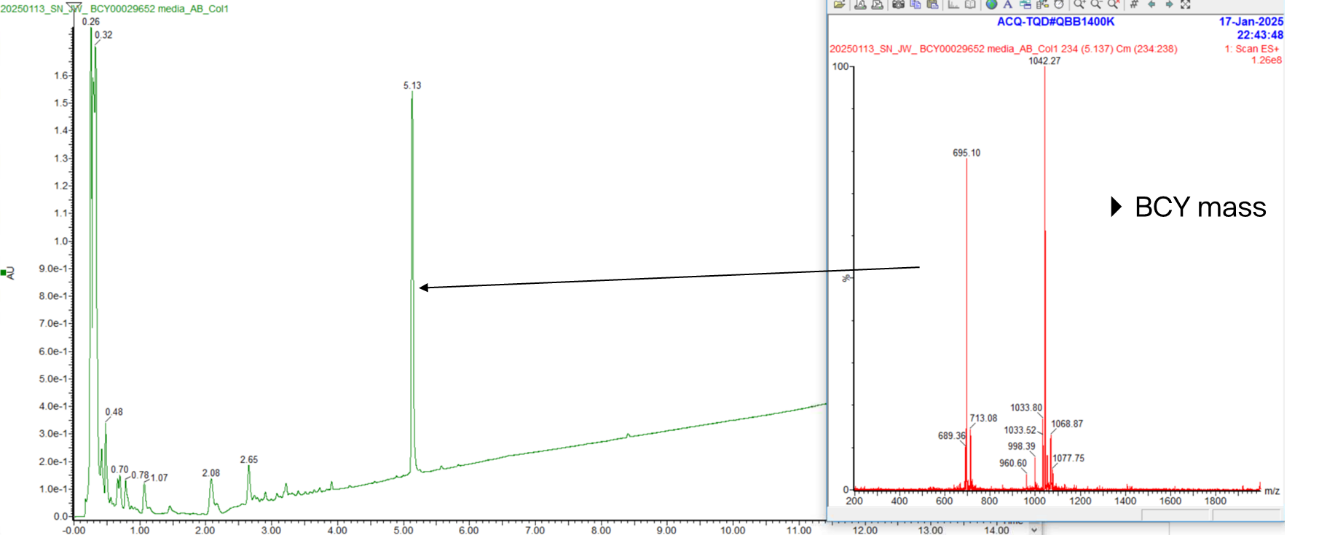
Peptide 24 + media

Peptide 24 + *S. aureus* supernatant

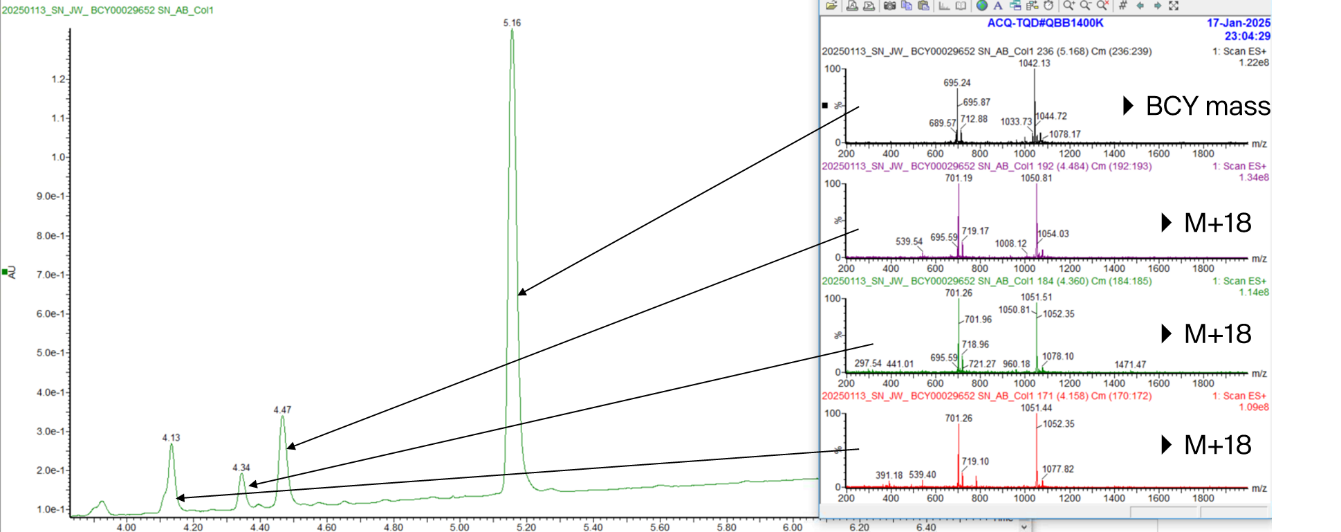

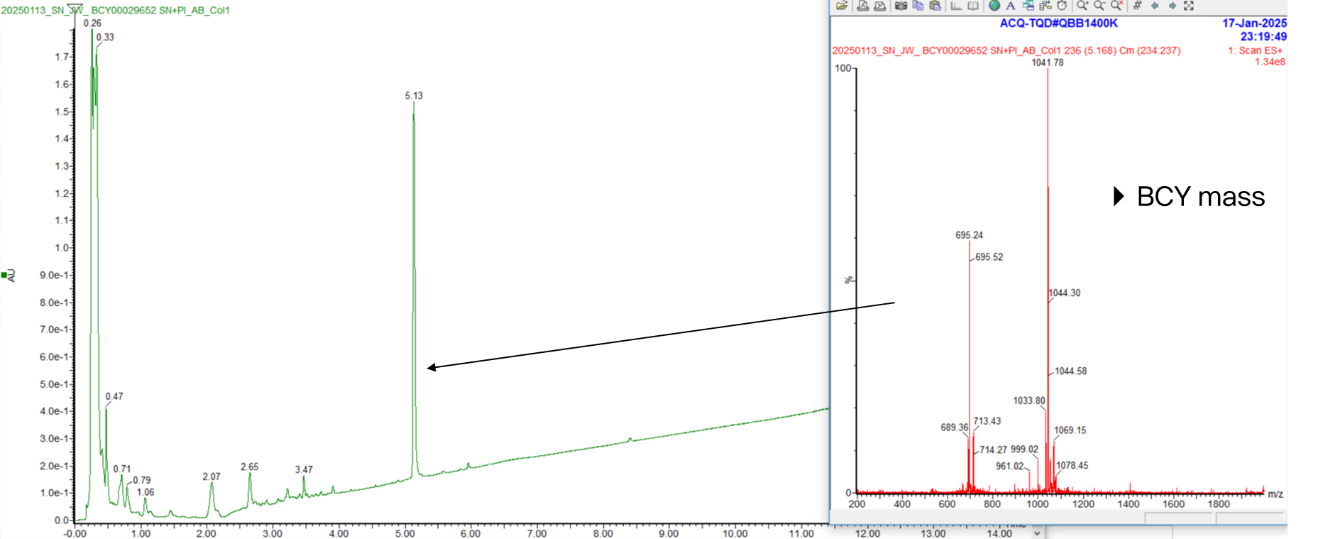
Peptide 24 + *S. aureus* supernatant + protease inhibitors

**Peptide 25 (MW: 2034.4)**

Peptide 25 + media

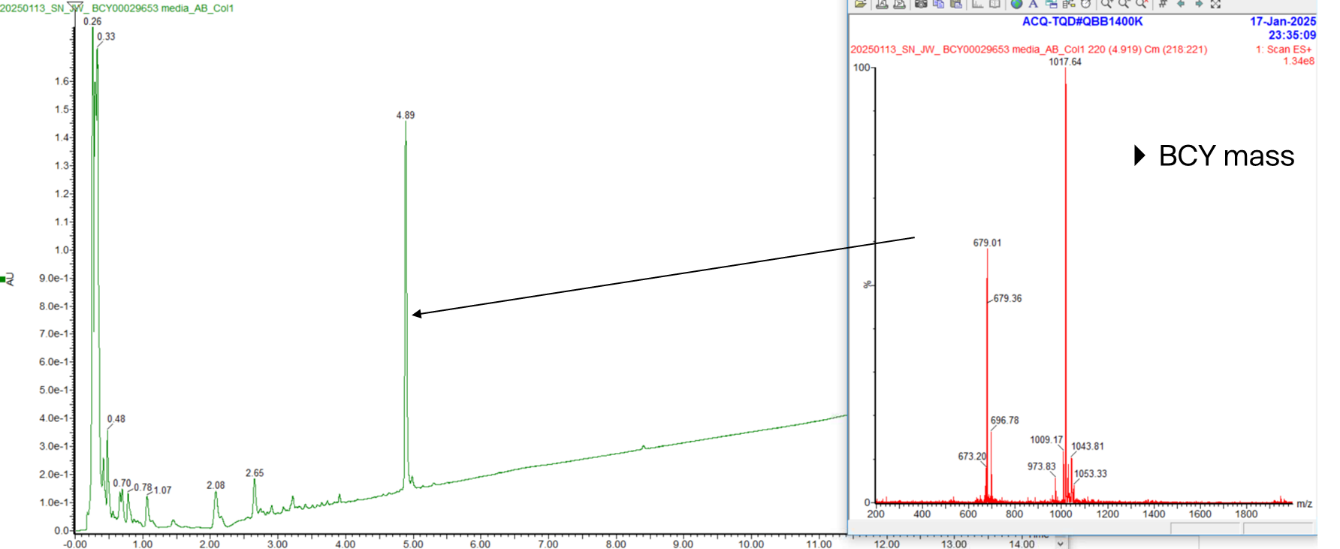

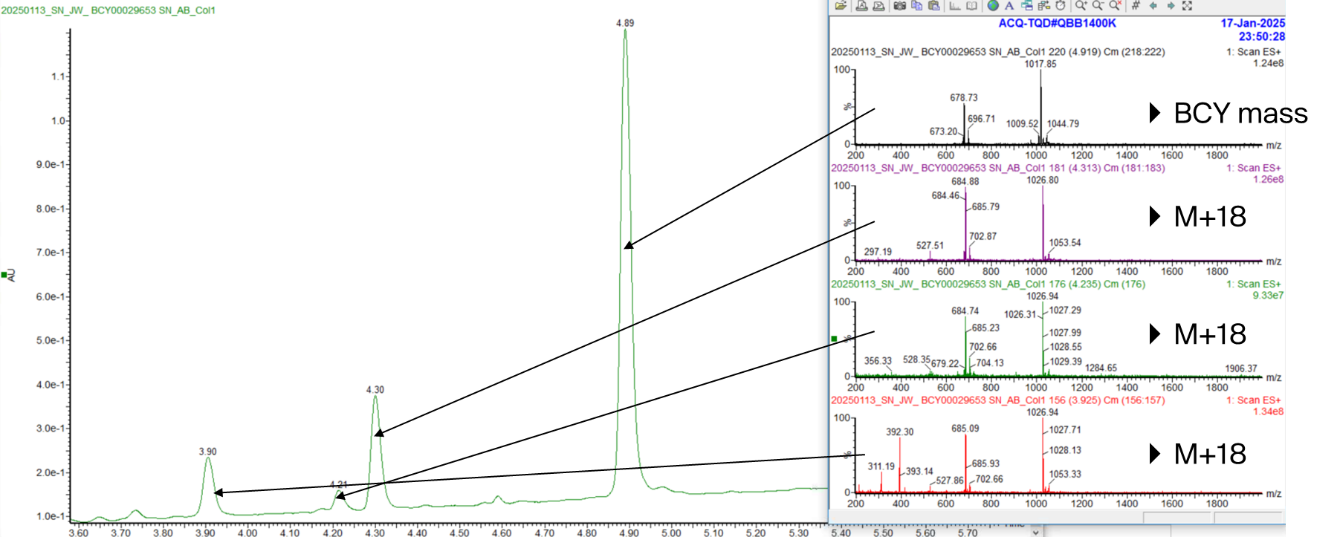
Peptide 25 + *S. aureus* supernatant

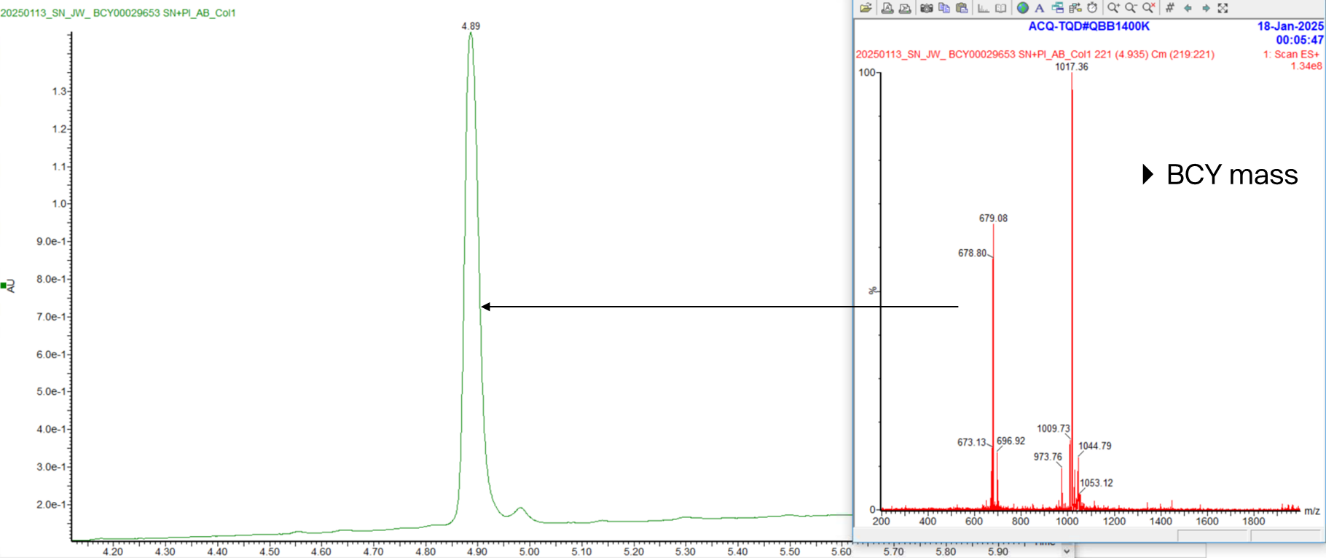
Peptide 25 + *S. aureus* supernatant + protease inhibitors

**Peptide 26 (MW: 2011.3)**

Peptide 26 + media

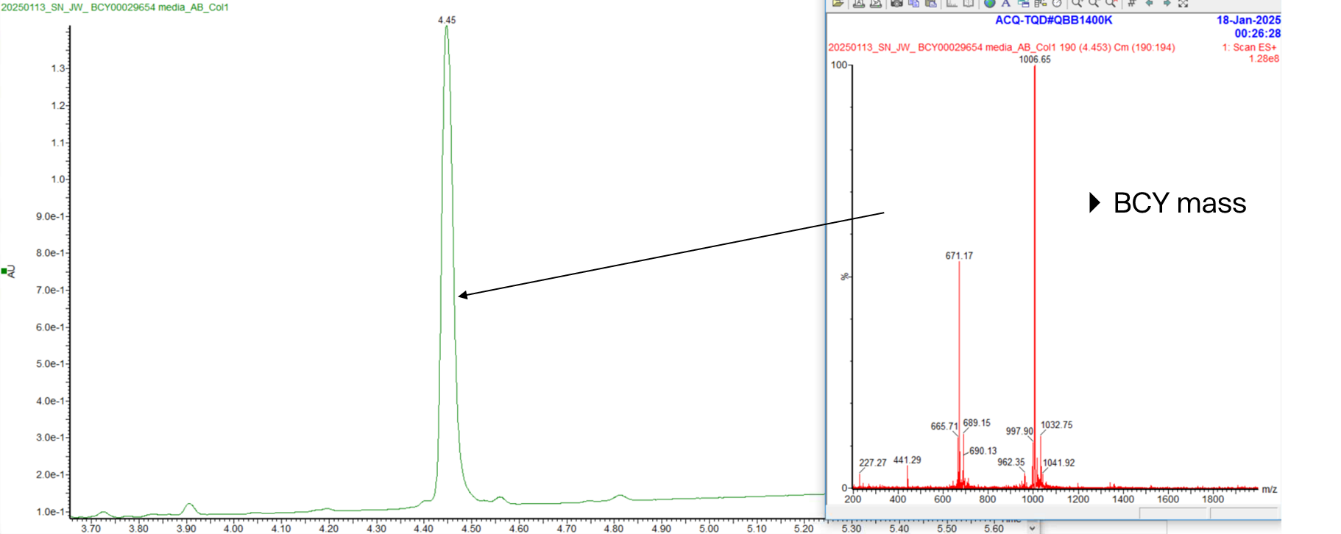

Peptide 26 + *S. aureus* supernatant

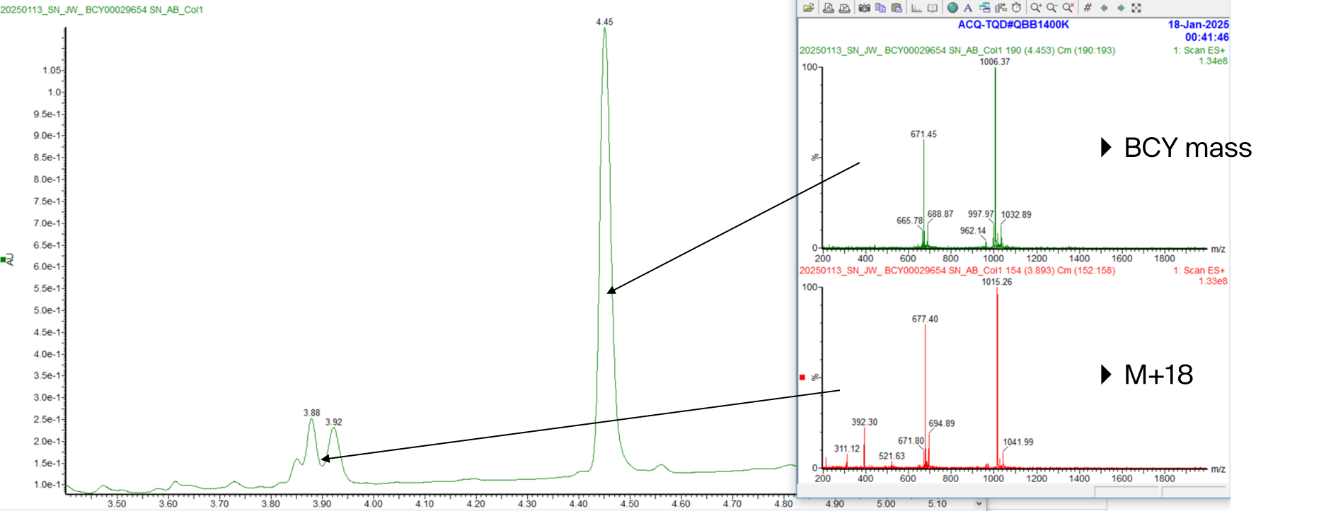

Peptide 26 + *S. aureus* supernatant + protease inhibitors

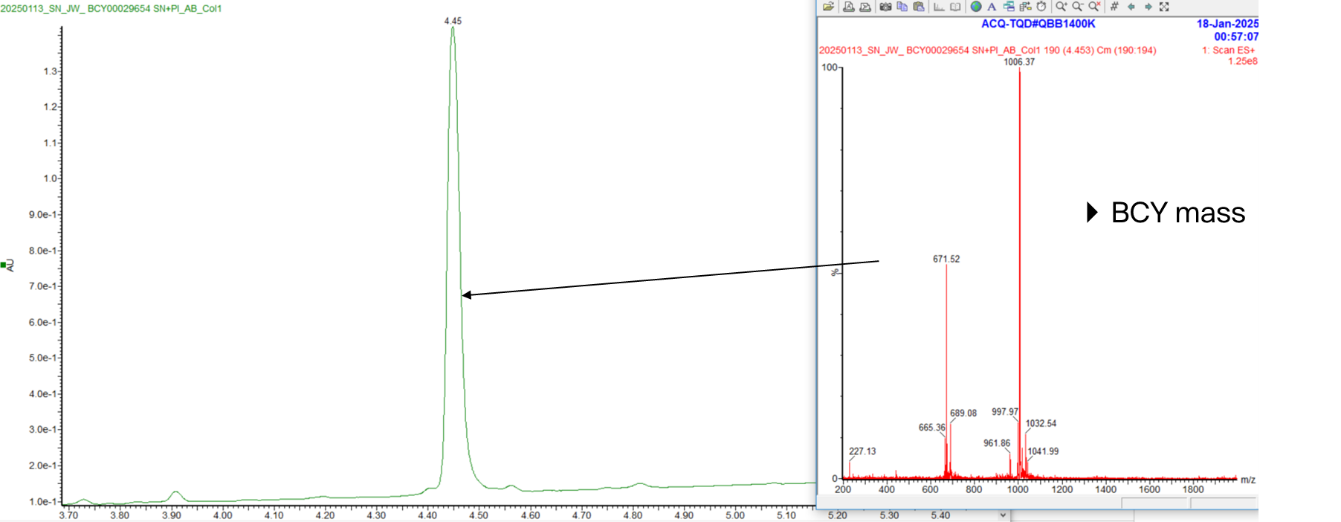

**Peptide 27 (MW: 2083.4)**

Peptide 27 + media

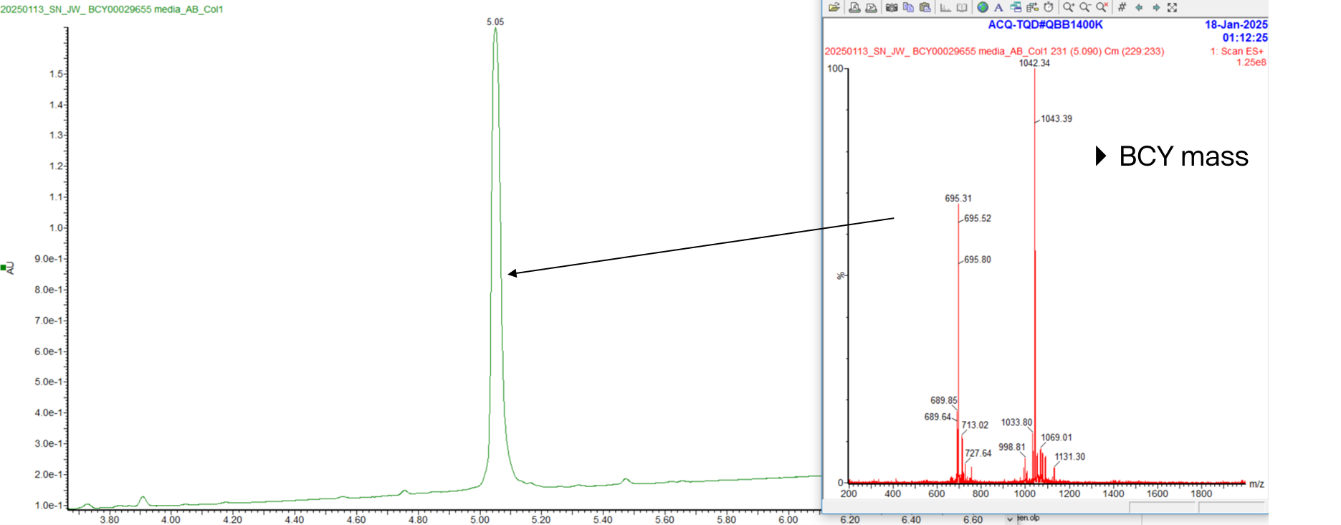

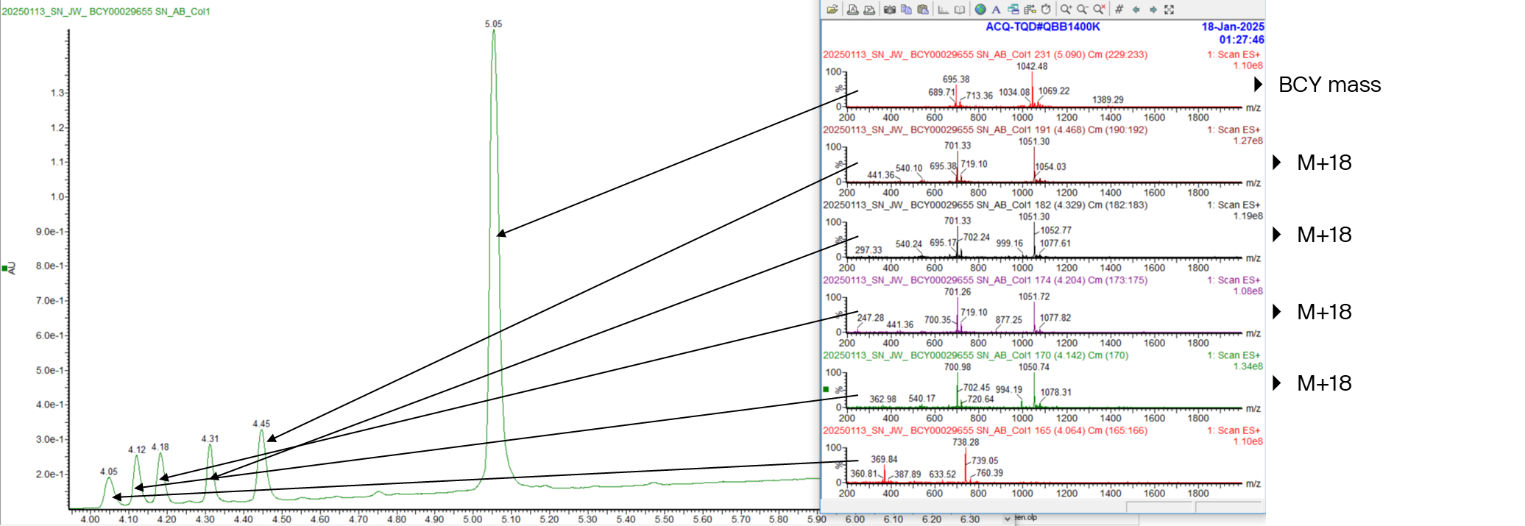
Peptide 27 + *S. aureus* supernatant

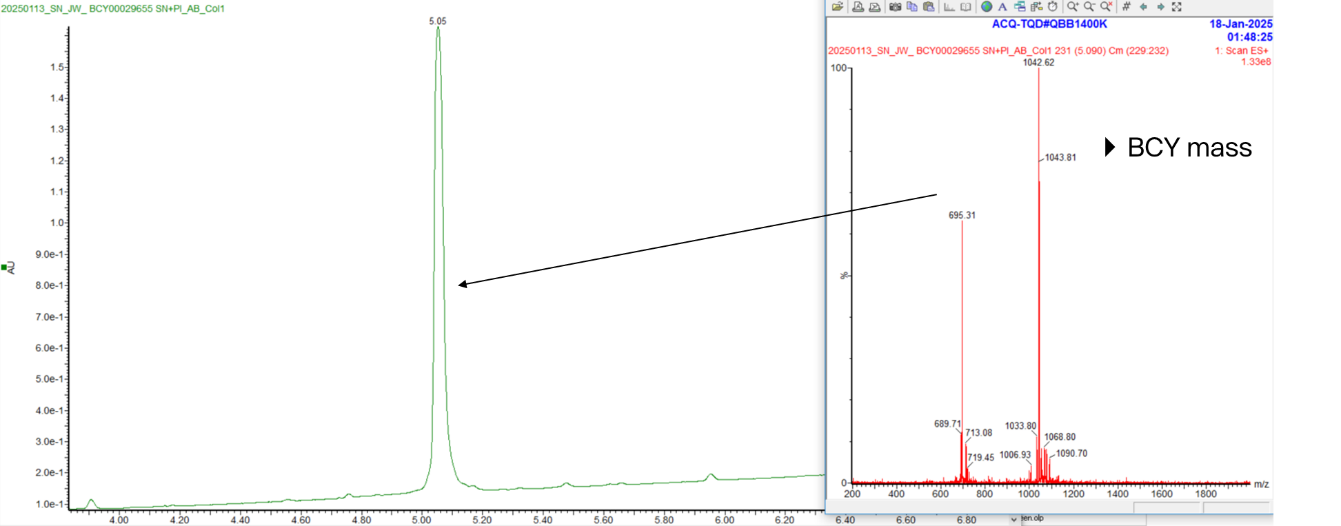
Peptide 27 + *S. aureus* supernatant + protease inhibitors

**Peptide 28 (MW: 2100.4)**

Peptide 28 + media

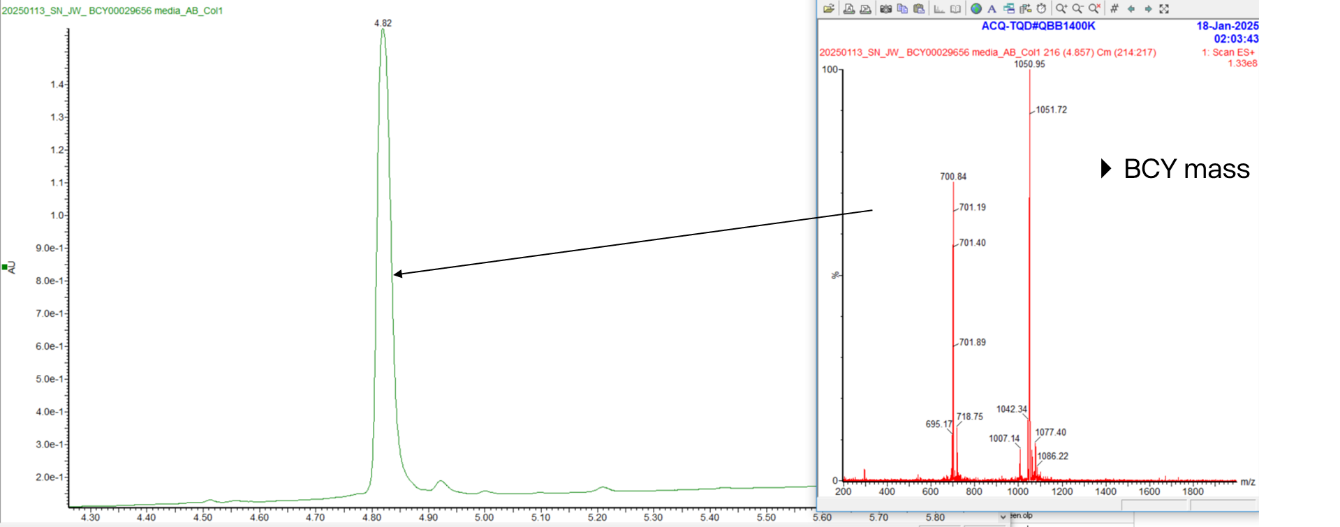

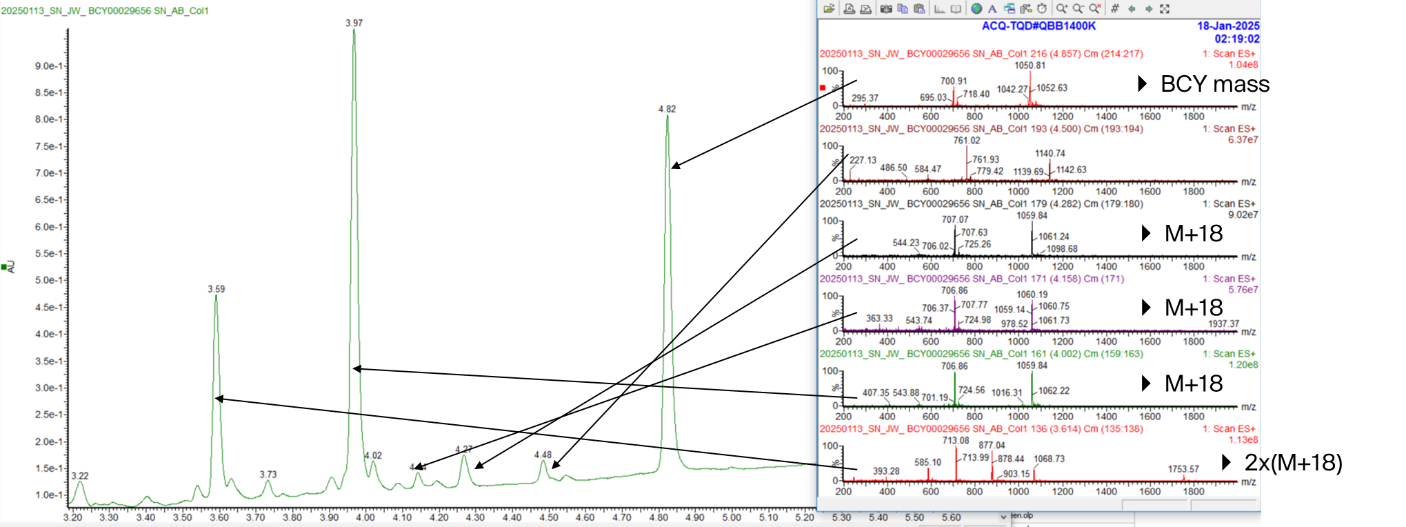
Peptide 28 + *S. aureus* supernatant

Peptide 28 + *S. aureus* supernatant + protease inhibitors

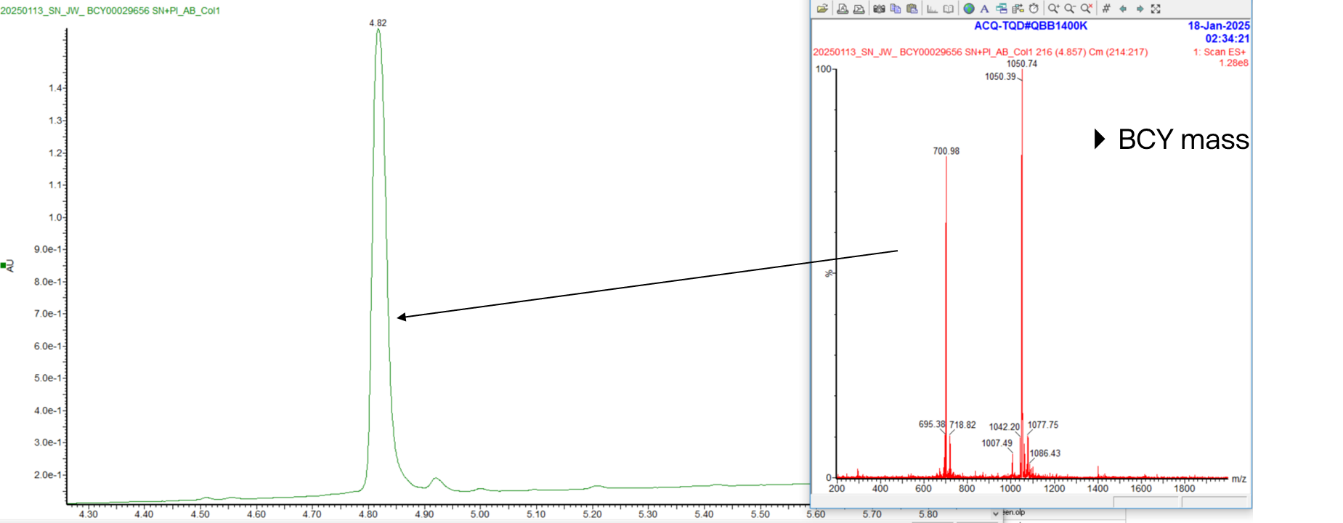

**Peptide 29 (MW: 2050.4)**

Peptide 29 + media

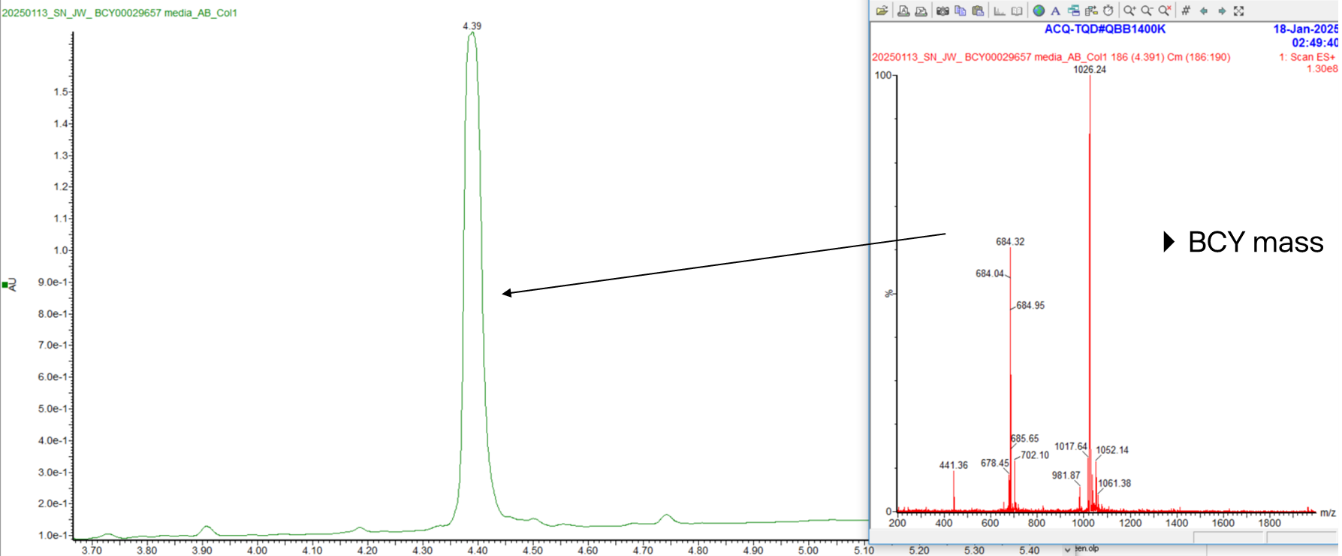

Peptide 29 + *S. aureus* supernatant

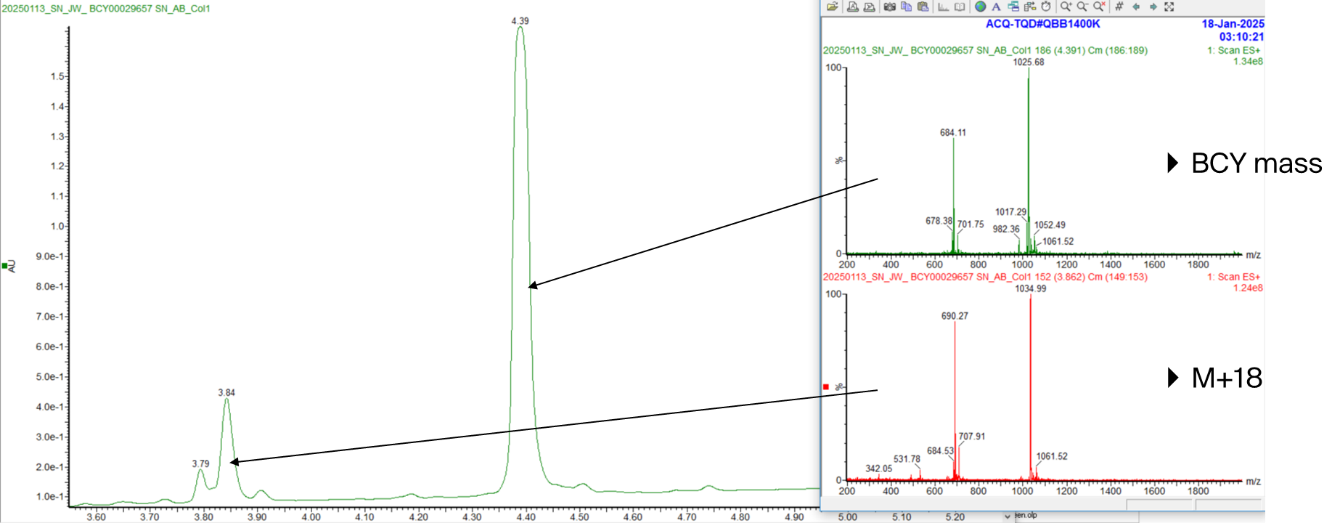

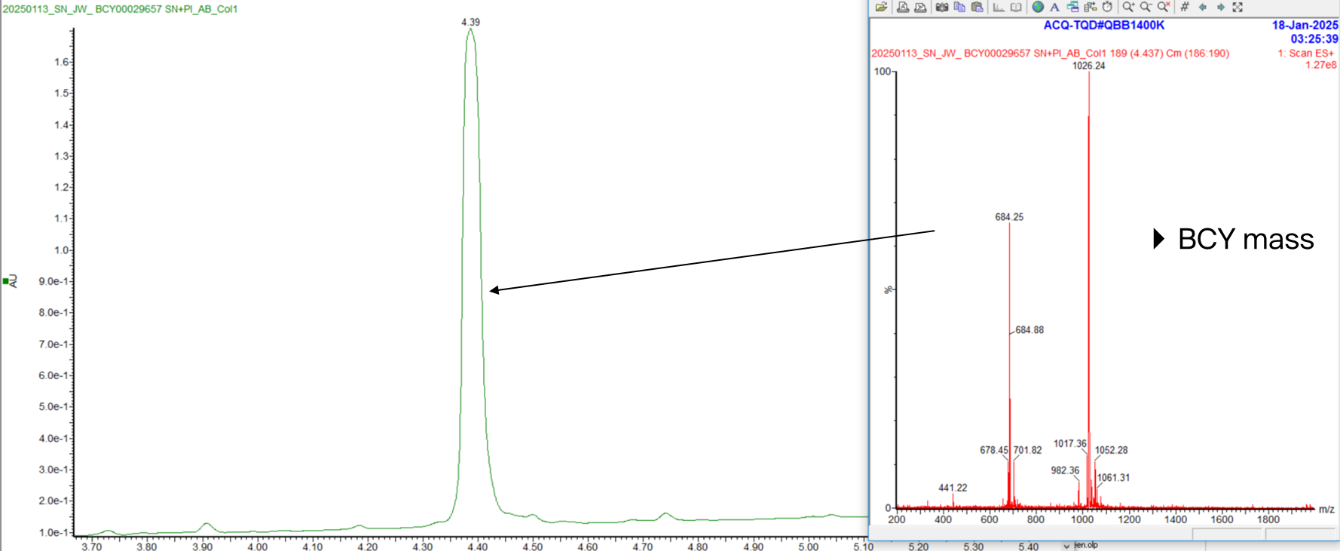
Peptide 29 + *S. aureus* supernatant + protease inhibitors

**Peptide 31 (MW: 2110.5)**

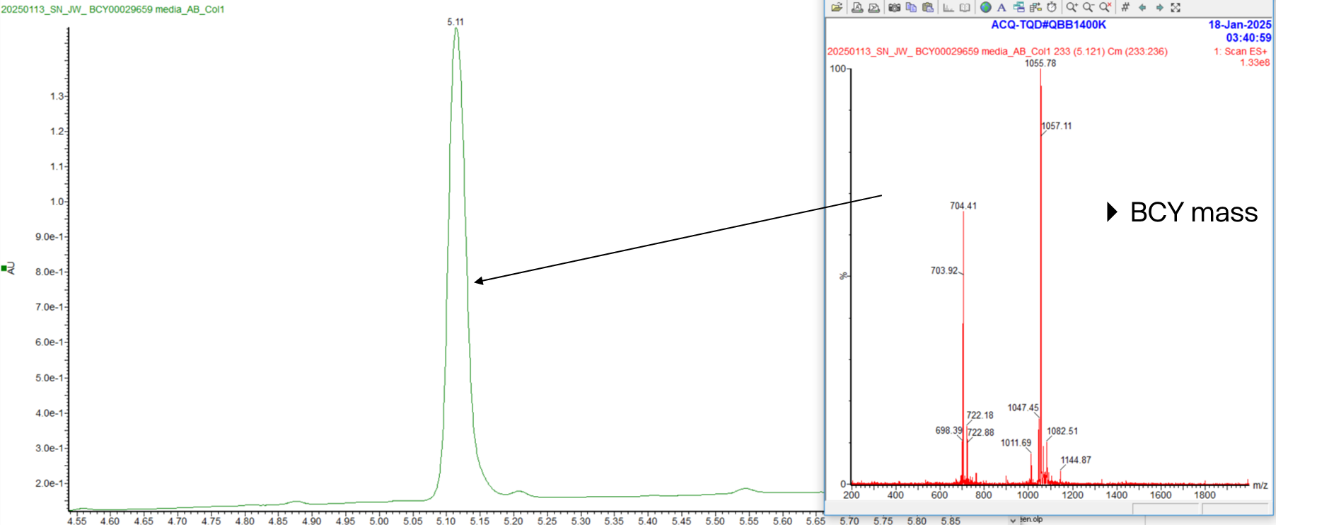
Peptide 31 + media

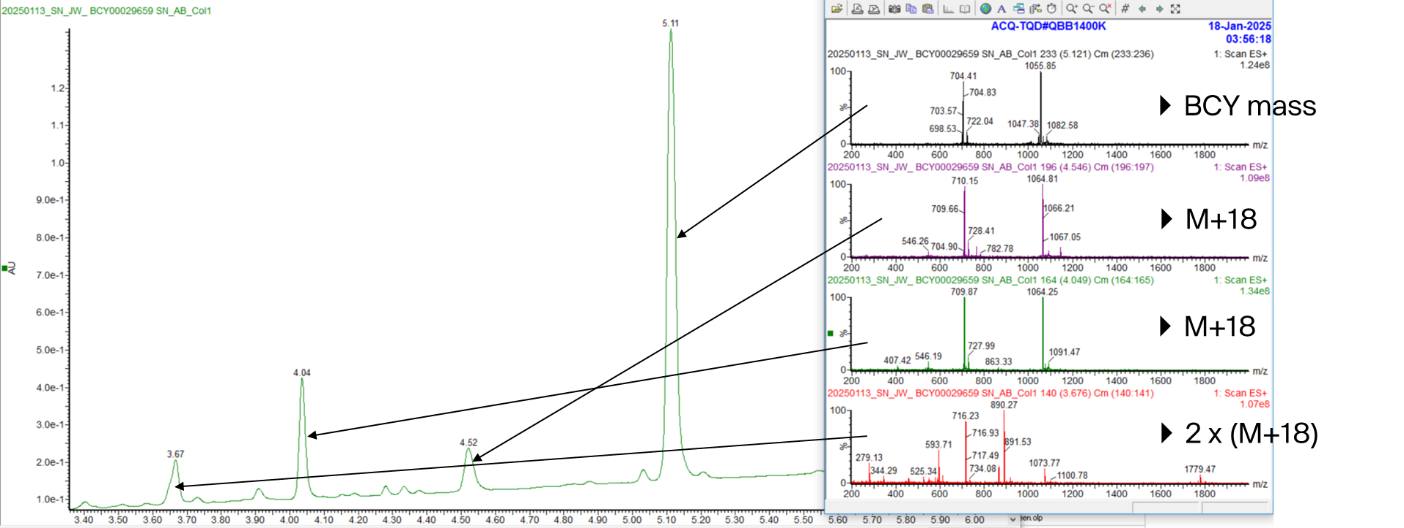
Peptide 31 + *S. aureus* supernatant

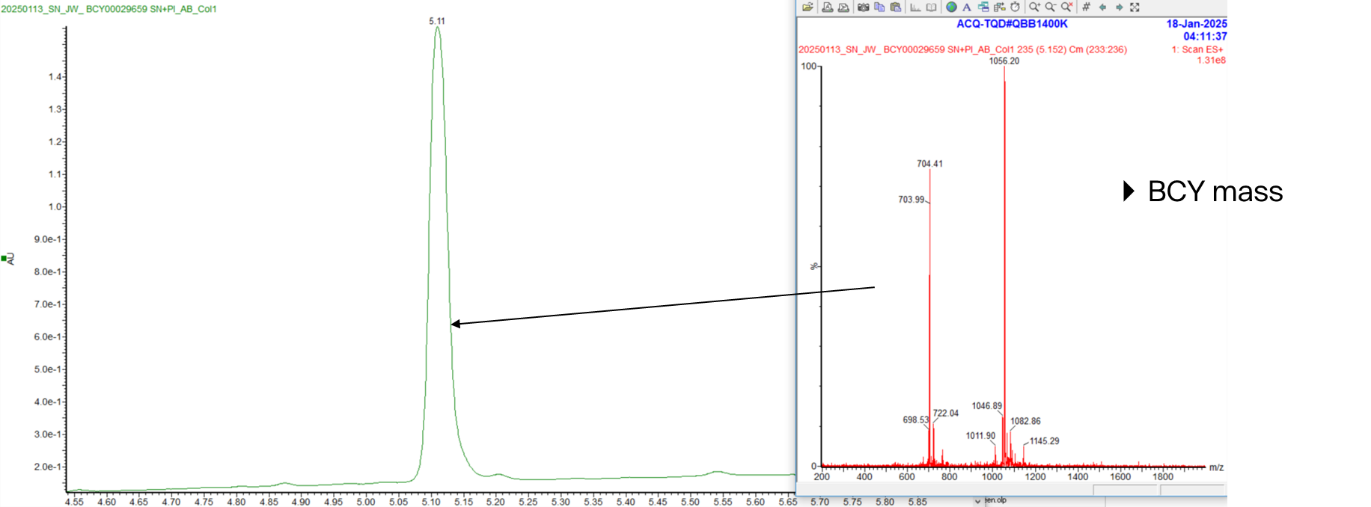
Peptide 31 + *S. aureus* supernatant + protease inhibitors

**Peptide 32 (MW: 2098.4)**

Peptide 32 + media

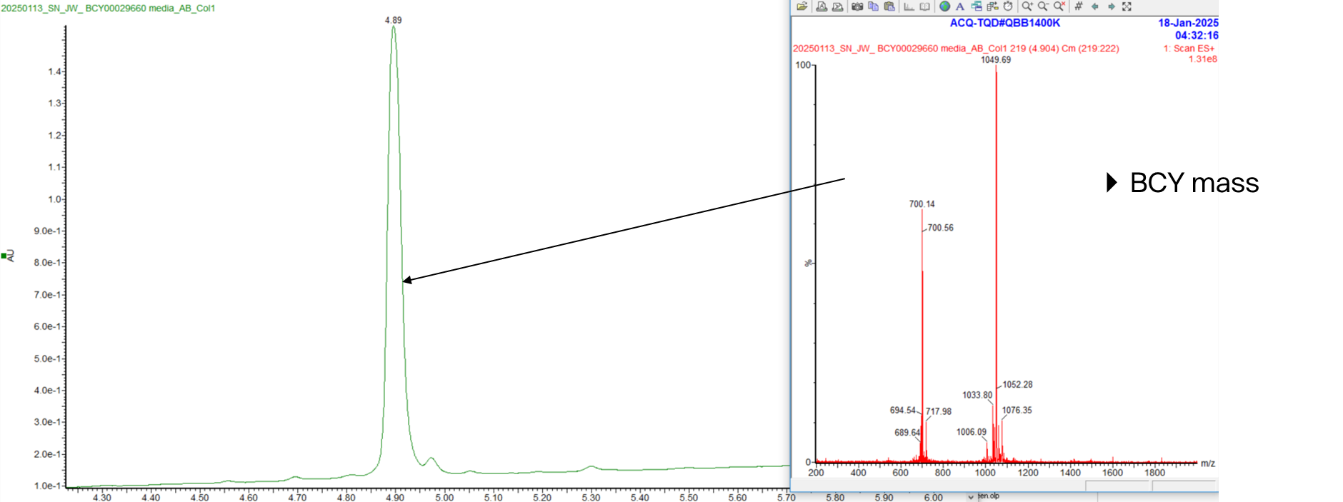

Peptide 32 + *S. aureus* supernatant

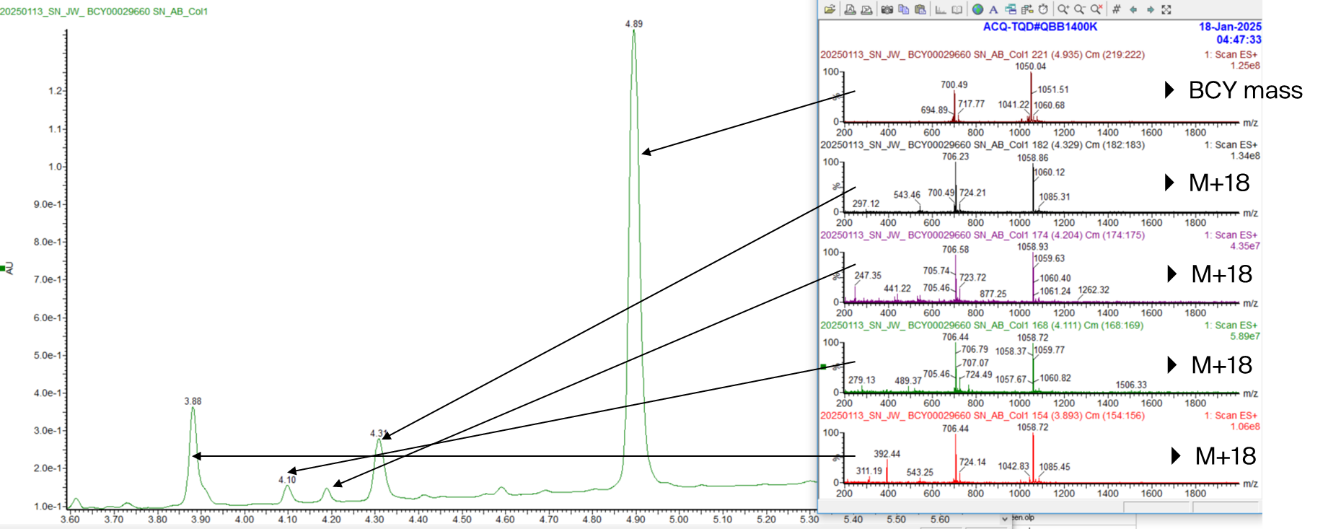

Peptide 32 + *S. aureus* supernatant + protease inhibitors

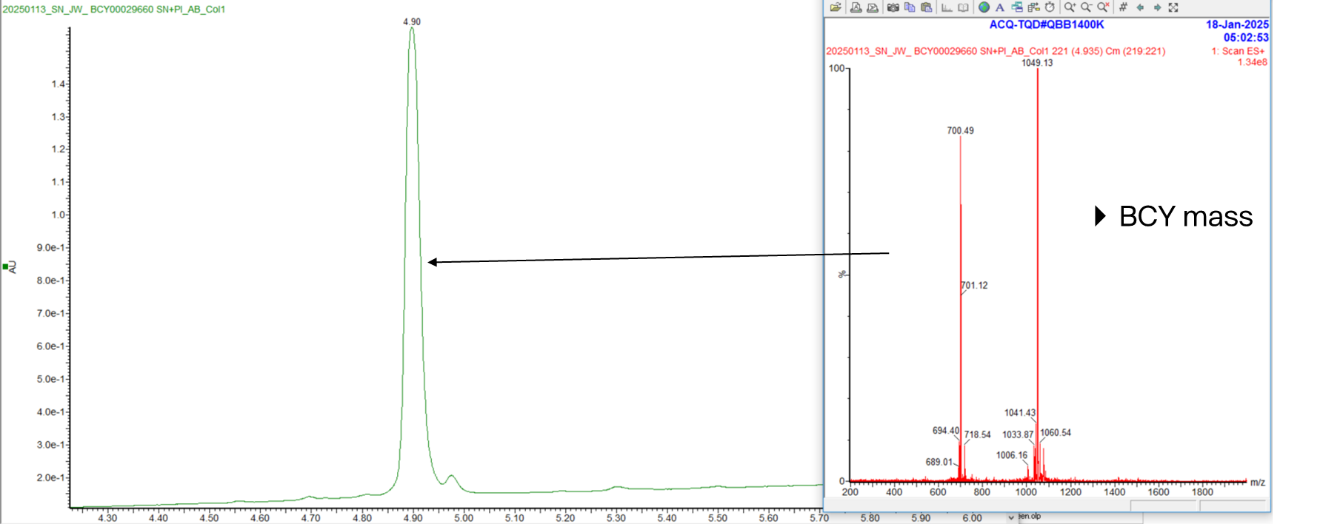

**Peptide 117 (MW: 2097.4)**

Peptide 117 + media

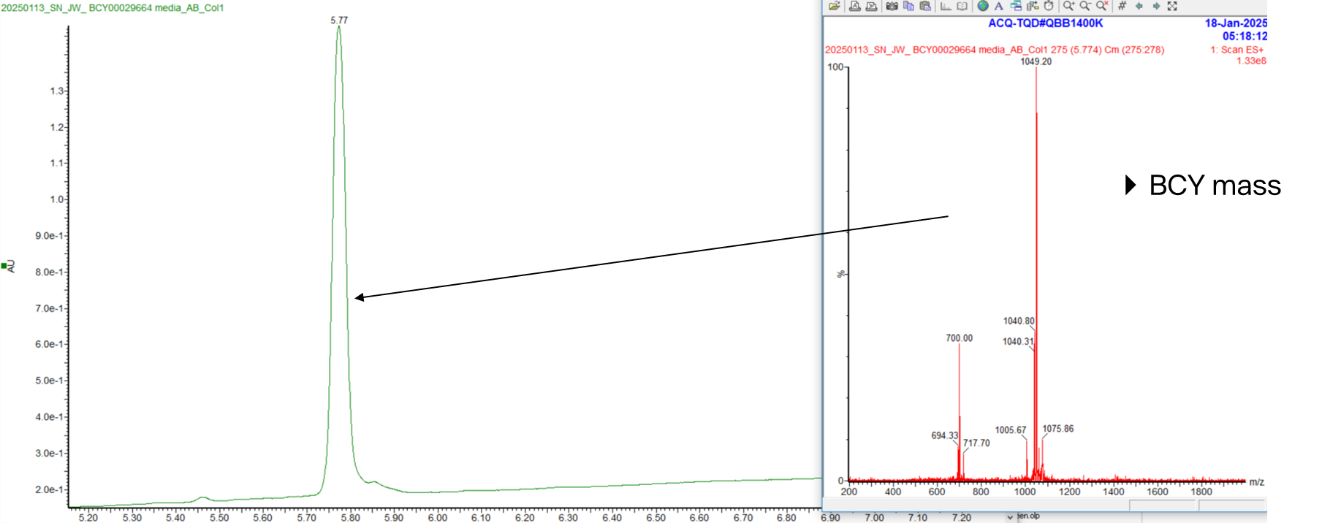

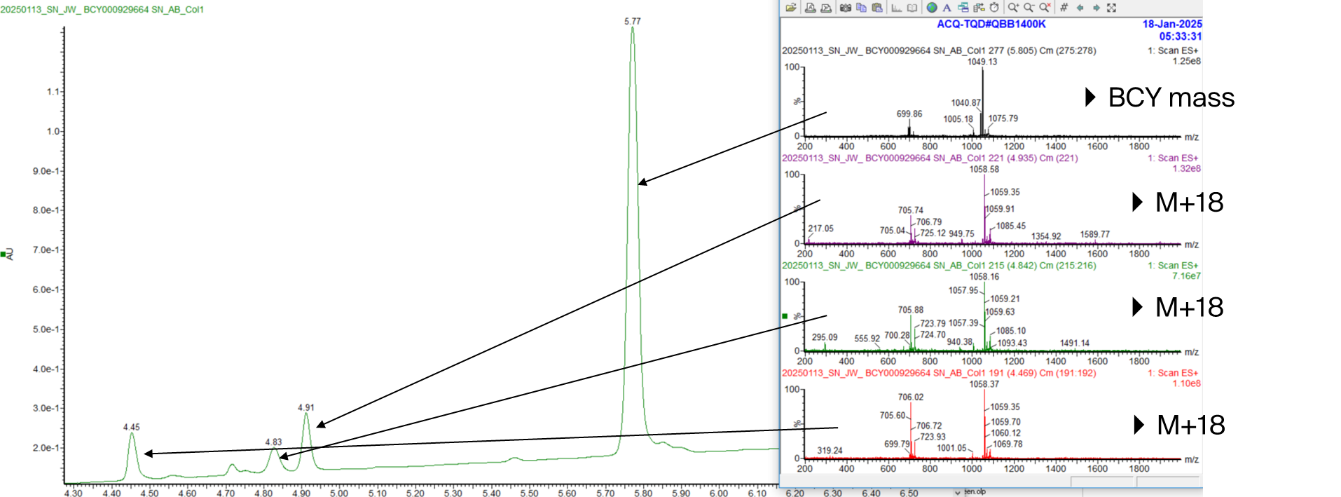
Peptide 117 + *S. aureus* supernatant

Peptide 117 + *S. aureus* supernatant + protease inhibitors

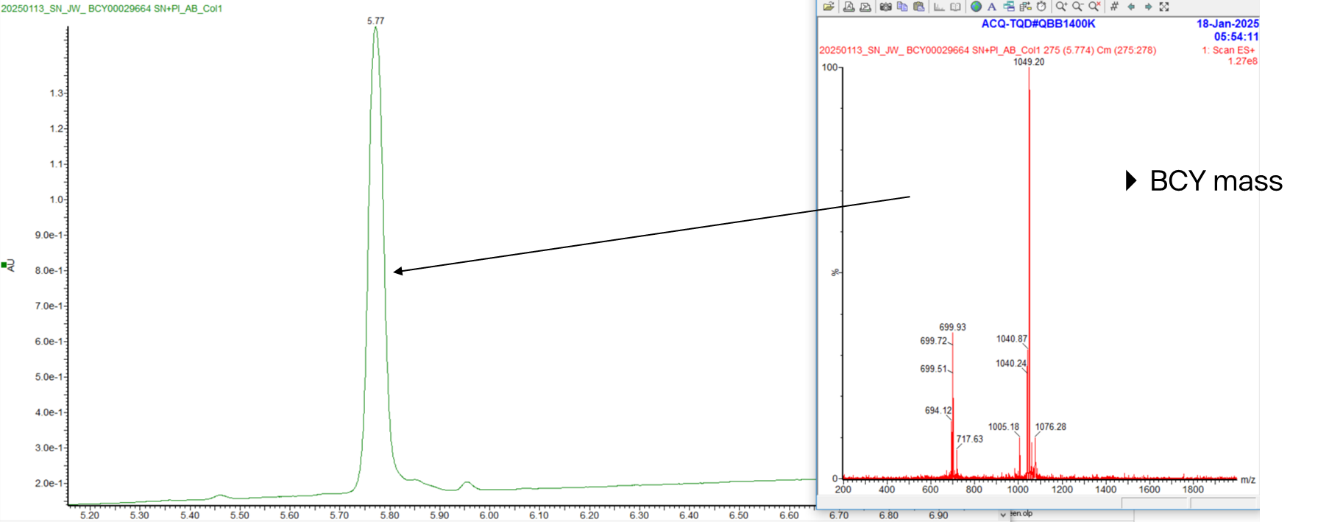

**Peptide 118 (MW: 1984.3)**

Peptide 118 + media

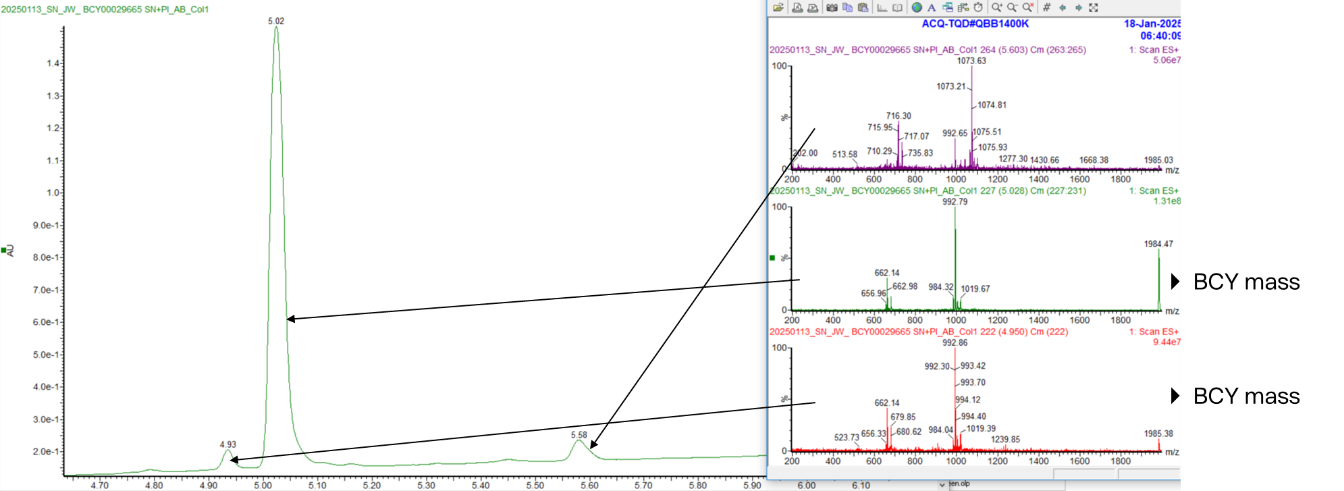

Peptide 118 + *S. aureus* supernatant

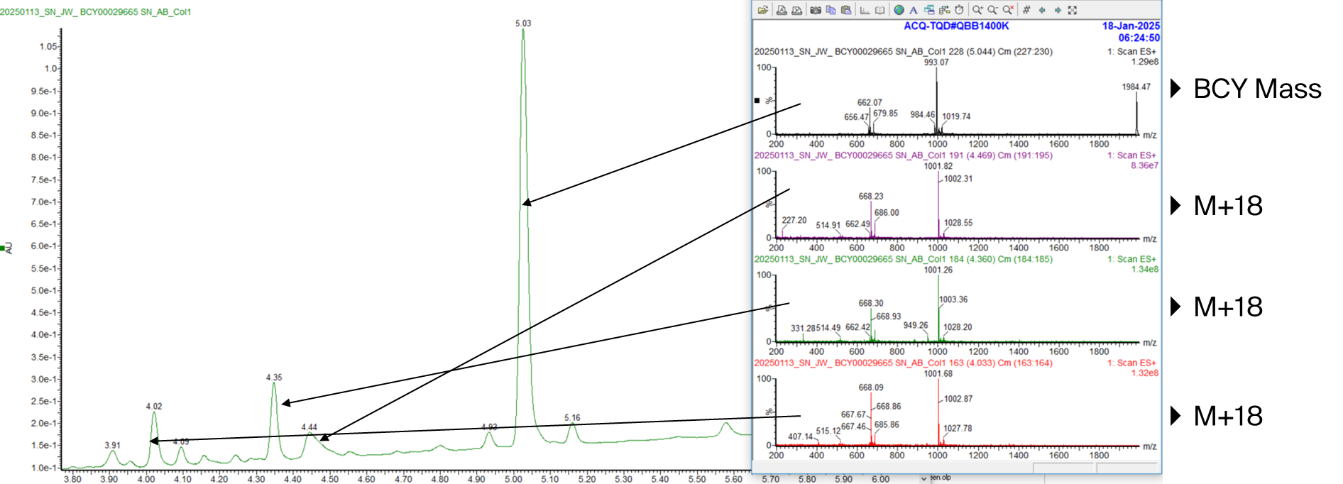

Peptide 118 + *S. aureus* supernatant + protease inhibitors

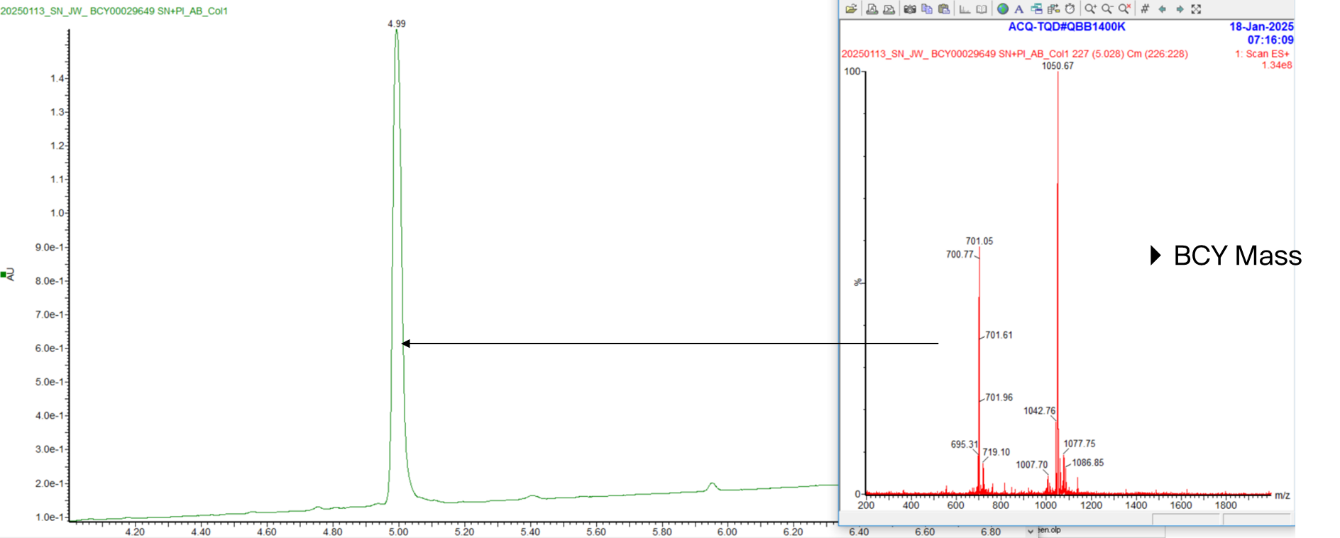
